# Role of channels in the O_2_ permeability of murine red blood cells II. Morphological and proteomic studies

**DOI:** 10.1101/2025.03.05.639962

**Authors:** Fraser J. Moss, Pan Zhao, Ahlam I. Salameh, Sara Taki, Amanda B. Wass, James W. Jacobberger, Dale E. Huffman, Howard J. Meyerson, Rossana Occhipinti, Walter F. Boron

## Abstract

In this second of three papers, we examine red blood cell (RBC) morphometry and RBC-membrane proteomics from our laboratory mouse strain (C57BL/6_Case_). In <u>paper #1</u>, using stopped-flow absorbance spectroscopy to ascertain the rate constant for oxyhemoglobin (HbO_2_) deoxygenation (*k*_HbO_2__), we find substantial *k*_HbO_2__ reductions with (1) membrane-protein inhibitors p-chloromercuribenzenesulfonate (pCMBS) or 4,4’-diisothiocyanatostilbene-2,2’-disulfonate (DIDS); (2) knockouts of aquaporin-1 (AQP1-KO), or Rhesus blood-group–associated A-glycoprotein (RhAG-KO), or double knockouts (dKO); or (3) inhibitor+dKO. In <u>paper #3</u>, reaction-diffusion mathematical modeling/simulations reveal that *k*_HbO_2__ could fall secondary to slowed intracellular O_2_/HbO_2_/Hb diffusion. Here in <u>paper #2</u>, blood smears as well as still/video images and imaging flow cytometry (IFC) of living RBCs show that ∼97.5% to ∼98.6% of control (not drug-treated) cells are biconcave disks (BCDs) across all genotypes. Pretreatment with pCMBS raises non-BCD abundance to ∼8.7% for WT and ∼5.7% for dKO; for DIDS pretreatment, the figures are ∼41% and ∼21%, respectively. Modeling (<u>paper #3</u>) accommodates for these shape changes. Light-scattering flow cytometry shows no significant difference in RBC size or shape among genotypes. IFC reveals minor differences among genotypes in RBC major diameter (Ø_Major_), which (along with mean corpuscular volume, <u>paper #1</u>) yields RBC thickness for simulations in paper #3. Label-free liquid chromatography/tandem mass spectrometry (LC/MS/MS) proteomic analyses of RBC plasma-membrane ghosts confirm the deletion of proteins targeted by our knockouts, and rule out changes in the 100 proteins of greatest inferred abundance. Thus, genetically induced changes in *k*_HbO_2__ must reflect altered abundance of AQP1 and /or the Rh complex.

**Key Points:**

- O_2_-offloading from red blood cells (RBCs) depends not only on membrane O_2_ permeability and oxyhemoglobin dissociation, but also on RBC size and shape. In this second of three papers, we use blood smears, still/video images of living RBCs, and imaging flow cytometry to examine morphometry of RBCs from <u>paper #1</u>.
- We find that mouse RBCs of all genotypes—wild-type, aquaporin-1 knockout (AQP1-KO), Rhesus blood group-associated A-glycoprotein knockout (RhAG-KO), and double knockout—are dominantly biconcave discs, with ∼1.4% to ∼2.5% poikilocytosis (shape change, SC). Drug pre-treatment increases %SC.
- Using label-free liquid chromatography/tandem mass spectrometry to assess apparent abundance of RBC-ghost proteins, we find no significant differences among genotypes for any of the ∼100 most abundant protein species except, as appropriate, AQP1, RhAG, or Rhesus blood group D antigen.
- Thus, the substantial effects observed in <u>paper #1</u> cannot be attributed to differences in morphometry or protein content.

## Introduction

The uptake of O_2_ by red blood cells (RBCs) in the pulmonary capillaries, the carriage of O_2_ to the systemic tissues, and the offloading of O_2_ from RBCs in the systemic capillaries to support oxidative metabolism is central to the life of every mammal. Some investigators, influenced by an implicit assumption underlying the mathematical formulation of Krogh’s cylinder (Krogh, 1919; Kreuzer, 1982), have believed that membranes, including RBC membranes, offer no resistance to O_2_ diffusion. Others, influenced by the work of Overton (for reviews, see Missner & Pohl, 2009; Boron, 2010), have contended that, while membranes do offer resistance to O_2_ diffusion, the rate is solely contingent upon O_2_ solubility in membrane lipids (Forstner & Gnaiger, 1983).

The present contribution is the second in a series of three interdependent papers ^1^ that investigate the extent to which murine aquaporin-1 (AQP1), the Rh complex (mouse Rh_Cx_ = Rhesus blood group-associated A-glycoprotein, RhAG + Rhesus blood group D antigen, RhD), and potentially other RBC membrane proteins contribute to the O_2_ permeability of the murine RBC membrane (*P*_M,O_2__).

In the first paper (i.e., <u>paper #1</u>; Zhao *et al*., 2025), we expose murine RBCs to an O_2_ scavenger (creating an O_2_-free environment) in a stopped-flow (SF) absorbance-spectroscopy apparatus to determine the rate constant for dissociation of oxyhemoglobin (*k*_HbO_2__). We find that *k*_HbO_2__ falls markedly if we pretreat RBCs with p-chloromercuribenzenesulfonate (pCMBS) or 4,4’-diisothiocyanatostilbene-2,2’-disulfonate (DIDS)—known inhibitors of CO_2_-conducting channels—or if we obtain the RBCs from our laboratory strain of mice (C57BL/6_Case_) genetically deficient in *Aqp1* (i.e., *Aqp1–/–* genotype), *Rhag* (*Rhag–/–* genotype), or both (*Aqp1–/–Rhag*–/– genotype). Of course, *k*_HbO_2__ depends not only on *P*_M,O_2__ but also on the diffusion of O_2_, oxyhemoglobin (HbO_2_), and hemoglobin (Hb) through the cytosol of the RBC, as well as the rate of dissociation of O_2_ from HbO_2_ per se. Thus, in <u>paper #1</u>, we measure mean corpuscular volume (MCV), mean corpuscular hemoglobin (MCH), and mean corpuscular hemoglobin concentration (MCHC), other automated-hematological parameters, and the rate constant (*k*_HbO_2_→Hb_) of the reaction HbO_2_ → Hb + O_2_ in fully hemolyzed RBCs (i.e., in the absence of intact membranes). <u>Paper #1</u> also provides a brief summary of the essential contributions of <u>paper #2</u> (the present paper) and <u>paper #3</u> (Occhipinti *et al*., 2025).

In this second (the present) paper, we assess RBC morphology at a microscopic scale— including blood smears, still and video images of living RBCs, and imaging flow cytometry (IFC). We ask whether the cells are mature RBCs and shaped as biconcave discs (BCDs) and, if they are not, determine the prevalence and other characteristics of these other cells (i.e., poikilocytes), collectively termed non-BCDs (nBCDs). The IFC approach allows us to determine RBC major diameter (Ø_Major_), which—together with the above MCV—allow us to compute RBC thickness— an important input for the mathematical simulations (see next paragraph). The present paper also includes a proteomic analysis of RBC ghosts to determine whether the knockout (KO) of *Aqp1* and/or *Rhag* produces unexpected effects in the protein composition of the RBC membranes. We find that the RBC membranes of mice genetically deficient in *Aqp1* and/or *Rhag* lack only the intended proteins among the 100 proteins with the greatest inferred abundance. Here in <u>paper #2</u>, we make frequent reference to relevant elements in <u>paper #1</u> and <u>paper #3</u>.

In <u>paper #3</u>, we report the development of a novel reaction-diffusion model of O_2_ unloading from a RBC and the resulting mathematical simulations of the experiments from the <u>paper #1</u>. Importantly, these simulations draw from the MCV, MCH, and MCHC data from <u>paper #1</u>, as well as the BCD, nBCD data and Ø_Major_ data from the present paper. The simulations agree rather well with the physiological data, allowing us to extract *P*_M,O_2__ values from the simulated *k*_HbO_2__ data. Our graphical user interface (Huffman *et al*., 2025) expands the power of the model, especially for non-experts.

Together, the three papers show that RBC membranes of wild-type (WT) mice offer considerable resistance to O_2_ diffusion, thereby ruling out the implicit assumption of the Krogh mathematical formulation. Nevertheless, the resistance would be ∼10-fold higher yet if it were not for the *P*_M,O_2__ contributions of AQP1 (∼22%), Rh_Cx_ (∼36%), and at least one unidentified non-AQP1/non-Rh_Cx_ pathway that is blocked by pCMBS (∼36%). These results also rule out the Overton hypothesis, at least for murine RBCs. Thus, the work of the three papers changes the way we think of RBC O_2_ handling, and raises the possibility of modulating *P*_M,O_2__ by manipulating membrane proteins^2^.

## Methods

### Ethical approval and animal procedures

All animal procedures are reviewed and approved by the Institutional Animal Care and Use Committee (IACUC) at Case Western Reserve University (CWRU).

### Mice

The mice and the methodologies concerning the mice are the same as in paper #1^3^.

### Genotyping

The genotyping methodologies are the same as in paper #1^4^.

### Physiological solutions

The physiological solutions are a subset of those summarized in <u>paper #1</u>^5^, and replicated here as Table 1. Although in the SF experiments of <u>paper #1</u>, we titrated solutions at 10°C, here in <u>paper #2</u>, we make pH measurements at room temperature (RT), using a portable pH meter (model A121 Orion Star, Thermo Fischer Scientific (TFS), Waltham, MA) fitted with a pH electrode (Ross Sure-Flow combination pH Electrode, TFS). Beakers containing pH calibration buffers (pH at 6, 8 and 10; TFS), physiological solutions to be titrated, and the pH electrode in its storage solution (Beckman Coulter, Inc. Brea, CA) are all equilibrated at RT. We adjust pH with NaOH (5 M) or HCl (5 M). Osmolality is measured using a vapor pressure osmometer (Vapro 5520; Wescor, Inc., Logan, UT) and, as necessary, adjusted upward by addition of NaCl.

**Table 1.**
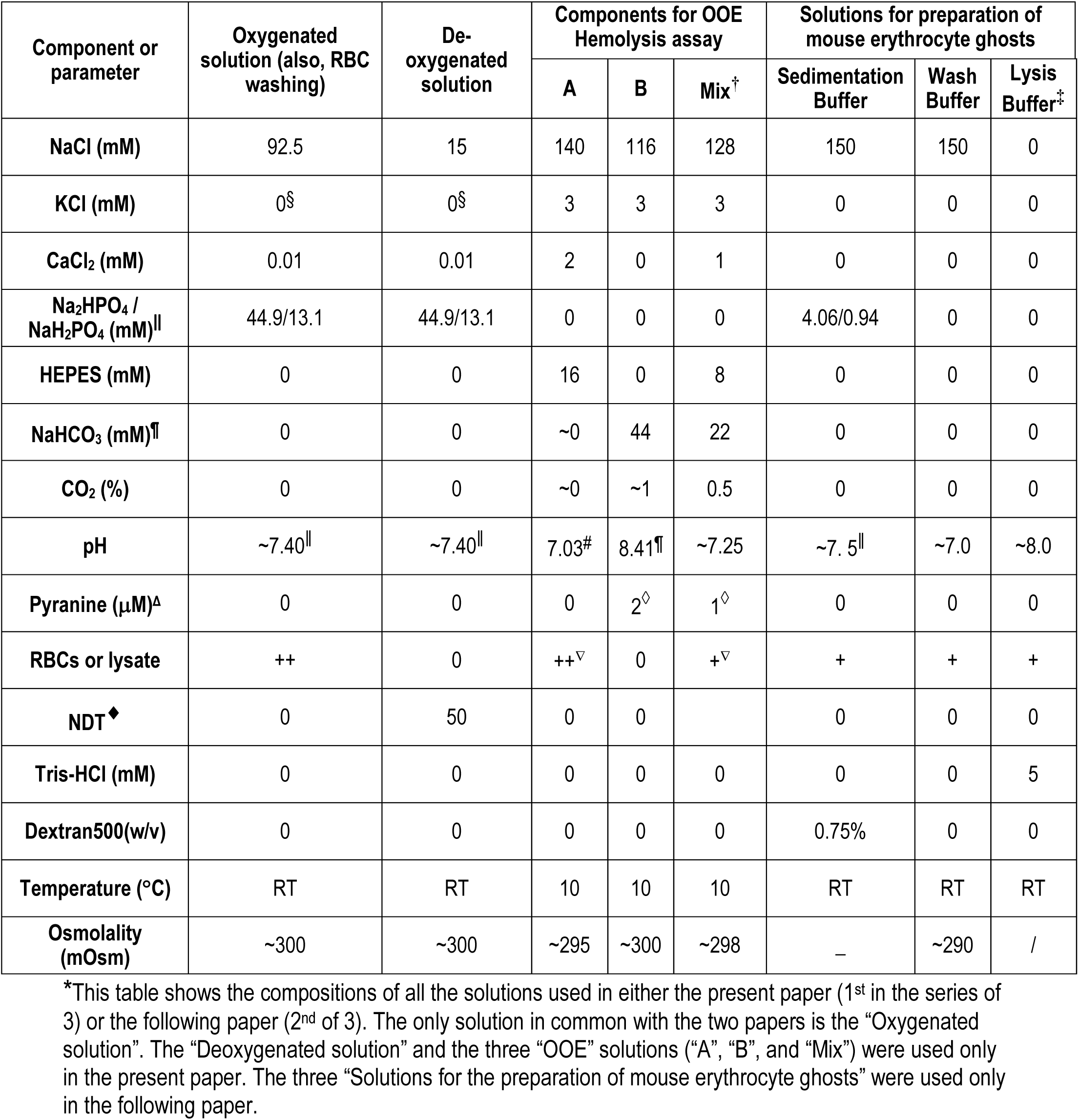
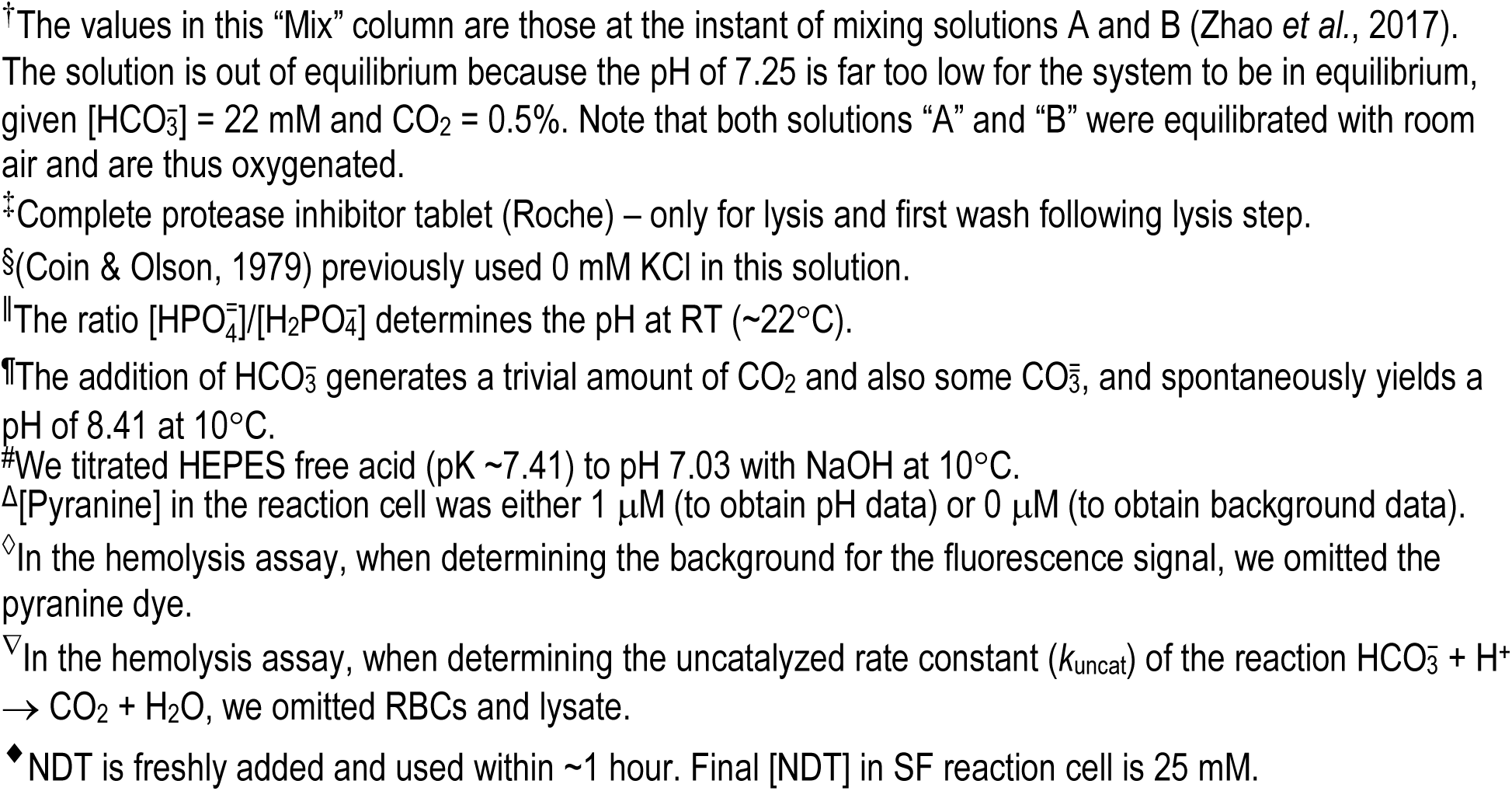
Physiological solutions*.

| Component or parameter | Oxygenated solution (also, RBC washing) | De-oxygenated solution | Components for OOE Hemolysis assay |  |  | Solutions for preparation of mouse erythrocyte ghosts |  |  |
| --- | --- | --- | --- | --- | --- | --- | --- | --- |
|  |  |  | A | B | Mix<sup>†</sup> | Sedimentation Buffer | Wash Buffer | Lysis Buffer<sup>‡</sup> |
| NaCl (mM) | 92.5 | 15 | 140 | 116 | 128 | 150 | 150 | 0 |
| KCl (mM) | 0<sup>§</sup> | 0<sup>§</sup> | 3 | 3 | 3 | 0 | 0 | 0 |
| CaCl<sub>2</sub> (mM) | 0.01 | 0.01 | 2 | 0 | 1 | 0 | 0 | 0 |
| Na<sub>2</sub> HPO<sub>4</sub> / NaH<sub>2</sub> PO<sub>4</sub> (mM) | 44.9/13.1 | 44.9/13.1 | 0 | 0 | 0 | 4.06/0.94 | 0 | 0 |
| HEPES (mM) | 0 | 0 | 16 | 0 | 8 | 0 | 0 | 0 |
| NaHCO<sub>3</sub> (mM) | 0 | 0 | ~0 | 44 | 22 | 0 | 0 | 0 |
| CO<sub>2</sub> (%) | 0 | 0 | ~0 | ~1 | 0.5 | 0 | 0 | 0 |
| pH | ~7.40 | ~7.40 | 7.03<sup>#</sup> | 8.41 | ~7.25 | ~7.5 | ~7.0 | ~8.0 |
| Pyranine (μM)<sup>Δ</sup> | 0 | 0 | 0 | 2<sup>◇</sup> | 1<sup>◇</sup> | 0 | 0 | 0 |
| RBCs or lysate | ++ | 0 | ++<sup>▽</sup> | 0 | +<sup>▽</sup> | + | + | + |
| NDT<sup>◆</sup> | 0 | 50 | 0 | 0 |  | 0 | 0 | 0 |
| Tris-HCl (mM) | 0 | 0 | 0 | 0 | 0 | 0 | 0 | 5 |
| Dextran500(w/v) | 0 | 0 | 0 | 0 | 0 | 0.75% | 0 | 0 |
| Temperature (°C) | RT | RT | 10 | 10 | 10 | RT | RT | RT |
| Osmolality (mOsm) | ~300 | ~300 | ~295 | ~300 | ~298 | — | ~290 | / |

\*This table shows the compositions of all the solutions used in either the present paper (1<sup>st</sup> in the series of 3) or the following paper (2<sup>nd</sup> of 3). The only solution in common with the two papers is the “Oxygenated solution”. The “Deoxygenated solution” and the three “OOE” solutions (“A”, “B”, and “Mix”) were used only in the present paper. The three “Solutions for the preparation of mouse erythrocyte ghosts” were used only in the following paper.
<sup>†</sup>The values in this “Mix” column are those at the instant of mixing solutions A and B (Zhao *et al.*, 2017). The solution is out of equilibrium because the pH of 7.25 is far too low for the system to be in equilibrium, given $[\text{HCO}_3^-] = 22 \text{ mM}$ and $\text{CO}_2 = 0.5\%$ . Note that both solutions “A” and “B” were equilibrated with room air and are thus oxygenated.
<sup>‡</sup>Complete protease inhibitor tablet (Roche) – only for lysis and first wash following lysis step.
<sup>§</sup>(Coin & Olson, 1979) previously used 0 mM KCl in this solution.
<sup>||</sup>The ratio $[\text{HPO}_4^{2-}]/[\text{H}_2\text{PO}_4^-]$ determines the pH at RT ( $\sim 22^\circ\text{C}$ ).
<sup>¶</sup>The addition of $\text{HCO}_3^-$ generates a trivial amount of $\text{CO}_2$ and also some $\text{CO}_3^{2-}$ , and spontaneously yields a pH of 8.41 at $10^\circ\text{C}$ .
<sup>#</sup>We titrated HEPES free acid ( $\text{pK} \sim 7.41$ ) to pH 7.03 with NaOH at $10^\circ\text{C}$ .
<sup>Δ</sup>[Pyranine] in the reaction cell was either $1 \text{ } \mu\text{M}$ (to obtain pH data) or $0 \text{ } \mu\text{M}$ (to obtain background data).
<sup>◇</sup>In the hemolysis assay, when determining the background for the fluorescence signal, we omitted the pyranine dye.
<sup>∇</sup>In the hemolysis assay, when determining the uncatalyzed rate constant ( $k_{\text{uncat}}$ ) of the reaction $\text{HCO}_3^- + \text{H}^+ \rightarrow \text{CO}_2 + \text{H}_2\text{O}$ , we omitted RBCs and lysate.
<sup>◆</sup>NDT is freshly added and used within $\sim 1$ hour. Final [NDT] in SF reaction cell is 25 mM.

### Inhibitors

The inhibitors and their sourcing are the same as <u>paper #1</u>^6^. We freshly prepare stock solutions of 2 mM pCMBS and 2 mM DIDS by dissolving the agents directly into our “Oxygenated solution” (see Table 1), taking care to shield the solutions from light.

#### Preparation of RBCs

<u>Paper #1</u>^7^ provides an overview of our approaches for collecting and preparing RBCs for various assays. Here we allude to general methodology for the sake of context, but focus on the methods particularly relevant to this second (the present) paper.

##### Collection of blood from mice

We collect fresh blood from WT or KO mice for seven assays: (1) SF, (2) still microphotography, (3) microvideography, (4) flow cytometry, (5) automated hematology, (6) blood smears, and (7) proteomics/mass spectrometry. For the work in the present paper, we collect blood (∼250 μl) exclusively using the submandibular-bleed method (Golde *et al*., 2005) with a 3-, 4-, or 5-mm point-length sterile animal lancet (MEDIpoint, Inc., Mineola, NY). We return to the same mouse no sooner than 72 h from a previous blood sample, using the contralateral side.

##### Processing of RBCs for assays 1–4

For the first four assays in the above list—SF (in <u>paper #1</u>^8^) as well as still microphotography, microvideography, and flow cytometry (the last three in the present paper)—we collect blood into 1.7-ml microcentrifuge tubes that are previously rinsed with 0.1% sodium heparin (H4784, Sigma-Aldrich, St. Louis, MO), and then centrifuged in a Microfuge 16 Microcentrifuge (Beckman Coulter, Inc.) at 600 × g for 10 min. We aspirate and discard the resulting supernatant and buffy coat. To remove residual extracellular Hb, we resuspend the pelleted RBCs in our oxygenated solution (Table 1) to a hematocrit (Hct) of 5% to 10%, centrifuge at 600 × g for 5 min. After four such washes, we resuspend RBCs in oxygenated solution to a final Hct of 25% to 30%, and maintain on ice (for up to ∼6 h) for experiments. At this point, the RBCs are directed to studies of SF (see <u>paper #1</u> ^9^), still microphotography and microvideography (see <u>below</u>^10^), or flow-cytometry (see <u>below</u>^11^).

##### Automated hematology

For automated hematological studies (see paper #1^12^), we collect whole blood into 20-μl plastic Boule MPA Micro pipettes (EDTA-K2, Boule Medical AB, Stockholm, Sweden).

##### Blood smears

For blood-smear studies (see <u>below</u>^13^), we collect whole blood into 2 ml K2E K2EDTA VACUETTE® tubes (Grelner Bio-One North America Inc., Monroe, NC).

##### Proteomic analysis

For proteomic analysis/mass spectrometry (see Proteomic analysis^14^) we collect whole blood into 1.7-ml heparinized microcentrifuge tubes.

### Blood smears

#### Preparation of RBCs

We collect fresh blood from three mice of each genotype (see <u>above</u>^15^) into 2-ml K2E K2EDTA VACUETTE® tubes (Monroe, NC 28110, USA). Blood smears are prepared using microscope slides (Fisher scientific, Pittsburgh, USA), and stained using Wright’s stain on a Sysmex SP-10 autostainer (Kobe, Japan). Air-dried smears are placed in staining cassettes, and staining performed per manufacturer’s specifications. Blood smears images (1000× magnification) are taken on an Olympus BH-2 microscope (Tokyo, Japan) with a DP73 digital camera attachment (Olympus), visualized with cellSens software (Olympus), and reviewed by a board-certified hematopathologist (H.J.M.).

#### Use of inhibitors in blood-smear studies

After two washes, we resuspend packed RBCs into our oxygenated solution (see Table 1; [Hb] = 5 μM, Hct ≅ 0.3%) without drugs or with either pCMBS (1 mM for 15 min) or DIDS (200 µM for 1 h). After centrifuging RBCs at 600 × g for 5 min, we aspirate and discard the resulting supernatant to remove inhibitors. RBCs are then resuspended in our oxygenated solution to make blood smears, following our standard procedure, outlined in the previous paragraph. We repeat this on three different days, for a total of three mice of each genotype.

#### Still microphotography and microvideography of living RBCs

We perform experiments (still or video imaging) on fresh blood, collected and processed as described in <u>Methods</u>^16^, one sample per mouse for each of four genotypes, all on the same day. We repeat this on three different days, for a total of three mice of each genotype, for both still and video studies (i.e., a total of 3+3 mice/genotype). After four washes and dilution of RBCs to an Hct of 25% to 30%, we suspend the cells in our oxygenated solution (see Table 1) to a final Hct of 0.5% to 1%, and store on ice for 30 – 120 min before imaging.

In preparation for the experiment, we acid-wash coverslips by immersing them in 2 M HCl for 30 min, performing 3 × 5 min washes with DNAase-/RNAase-free water, and finally immersed them in 100% ethanol for 30 min before final air drying. A droplet containing suspended RBCs from one mouse (see previous paragraph) is placed on one of these acid-washed glass coverslips that served as the bottom of a recording chamber. The chamber is then mounted on Olympus IX-81 inverted microscope equipped for differential interference contrast (DIC) studies, using either of two oil-immersion objectives (60× objective, NA 1.42 for still micrographs or 40× objective, NA 1.35 for microvideomicroscopy) with a 1.5× magnification selector. The light is detected with an intensified EM-CCD camera (C9100-13, Hamamatsu Corporation, Bridgewater, NJ) with 512 × 512 pixels, and data acquired using SlideBook 5.0 software (Intelligent Imaging Innovation, Denver, CO) for the Hamamatsu camera. We record still micrographs or microvideos (1 frame per 5 s) of the RBC droplet as RBCs fall freely through the plane of focus, toward the coverslip surface.

##### Use of inhibitors in still photomicrography

After 4 washes, as noted in the previous paragraph, we resuspend packed RBCs in our oxygenated solution (see Table 1) without drugs or with either pCMBS (1 mM for 15 min) or DIDS (200 µM for 1 hour) to a final Hct of 1% to 2%, and direct the RBCs to still-photomicrography studies.

#### Flow cytometry (workflow #6 & #6′, #15 & #15′)

Figure 1 summarizes the workflow for the three papers in the project, including input from the present paper. All flow cytometry experiments are performed at the Case Comprehensive Cancer Center Cytometry and Microscopy Share Resource at CWRU (CMSR).

**Figure 1.**
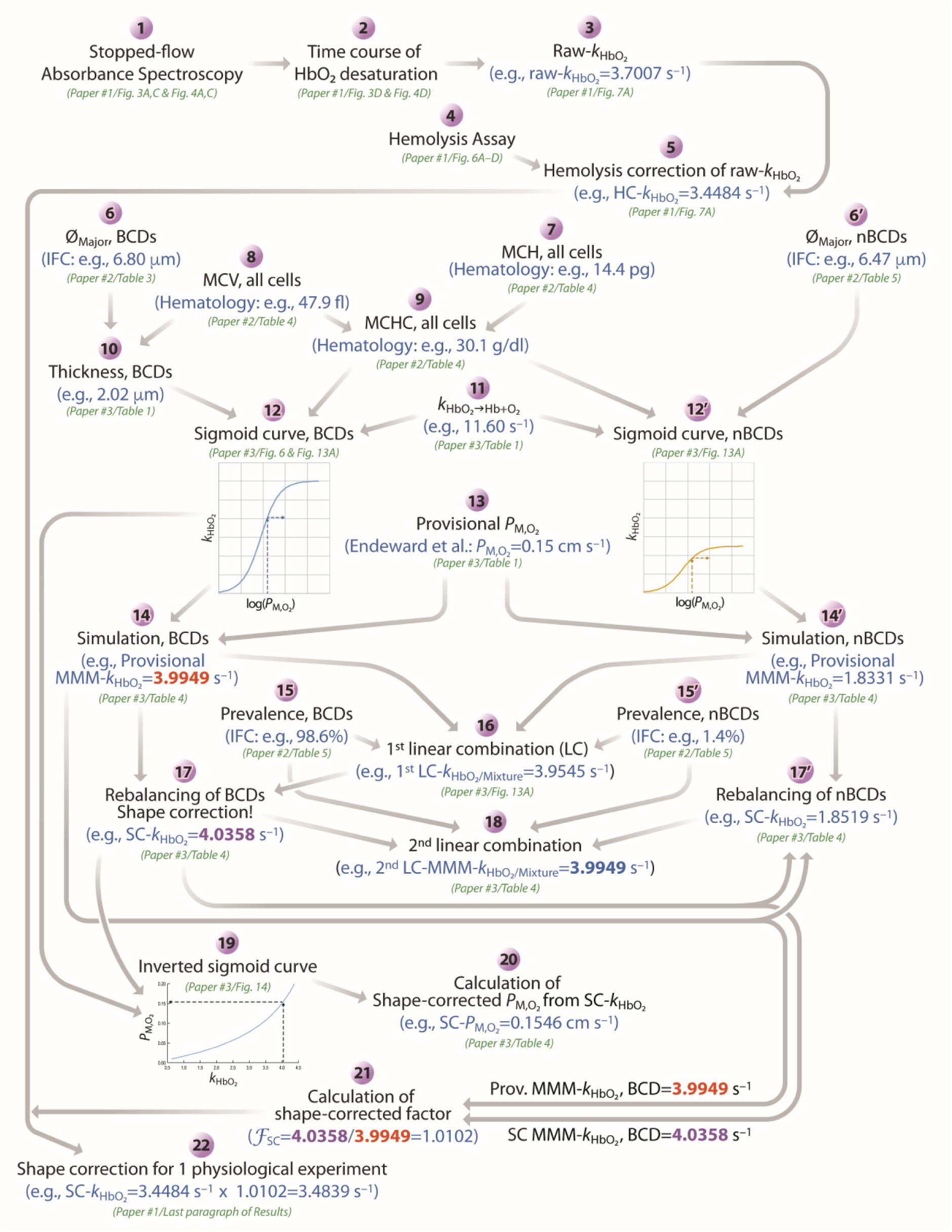
Workflow to obtain hemolysis-corrected and shape-corrected values of *P*_M,O_2__. The numerals 1 through 22 indicate the steps summarized in in <u>paper #1</u>—and presented in detail, as appropriate, in the present paper (<u>paper #2</u>), <u>paper #1</u>, or <u>paper #3</u>—to arrive, first, at a hemolysis-corrected (HC) *k*_HbO_2__ and, ultimately, at a shape-corrected (SC) *k*_HbO_2__. At each step, we provide example values, if possible, and example figure/table numbers referenced to <u>paper #1</u>, the present paper, or <u>paper #3</u>. We repeat workflow steps 1–5 for each experiment, ultimately arriving at a HC-*k*_HbO_2__ value for each. Step 6 applies to biconcave disks (BCDs), whereas step 6′ applies to non-biconcave disks (nBCDs). The same is true for steps 12 vs. 12′, 14 vs. 14′, 15 vs. 15′, and for 17 vs. 17′. *ℱ*_SC_, shape-correction factor; Hb, hemoglobin; IFC, imaging flow cytometry; MCH, mean corpuscular hemoglobin; *k*_HbO_2__, rate constant for deoxygenation of HbO_2_ within intact RBCs; *k*_HbO_2_→Hb+O_2__, rate constant for deoxygenation of HbO_2_ in free solution; LC, linear combination; MCH, mean corpuscular hemoglobin; MCHC, mean corpuscular hemoglobin concentration; MCV, mean corpuscular volume; MMM-k_HbO_2__, macroscopic mathematically modeled rate constant for deoxygenation of intracellular HbO_2_; *P*_M,O_2__; membrane permeability to O_2_; Prov., provisional; Ø_Major_, major diameter (of BCD or nBCD); SC-k_HbO_2__, shape-corrected rate constant for deoxygenation of intracellular HbO_2_.

##### Sample preparation

On a single day, we collect fresh blood (see <u>above</u>^15^) from 2 mice of each genotype. We repeat experiments on another day, for a total of four mice per genotype. To permit gating of viable RBC precursors, we dilute 100-μl samples of cells to a 1% Hct in our oxygenated solution (see Table 1) containing 1 μM Calcein Violet (CV; viability marker; excitation/emission 405/450 nm; TFS, C3099), 0.1 μg/ml Thiazol Orange (TO; to stain RNA; 490/530 nm; Sigma-Aldrich 390062), and 5 μM DRAQ-5 (to stain DNA; 647/683 nm; TFS, 62252), and then incubate for 20 min at RT in the dark. We then wash the dye-loaded RBC samples ×3 in 1 ml of our oxygenated solution, centrifuging at 600×g for 5 min between washes, and maintained stained cells on ice for up to ∼2 h on ice for experiments. The Hct was either 0.06% (∼2 million cells/ml) for light scattering on an LSRII flow cytometer (BD Biosciences; San Jose, CA) or 1% Hct for imaging on the ImageStreamX (Amnis Corporation, Seattle, WA) imaging flow cytometer.

##### Light-scattering flow cytometry

The first step is to generate a histogram—over a single sample of cells—that describes the relative number of cells (y-axis) vs. amount of scattered light (x-axis) (Loken *et al*., 1977). Forward scatter—measured by photodiodes positioned in line with the incident light source—is proportional to cell size. Forward-scatter intensity area (FSC-A) is the cumulative integration of the forward-scatter signal for the full period a cell traverses the laser path, specifically the total area underneath the corresponding histogram. The FSC-A parameter provides overall size and shape information for the cell, utilized alongside forward scatter pulse height (FSC-H), which quantifies the amplitude of the detected light scatter signal as cells traverse the laser beam, indicating cell size, and forward-scatter intensity width (FSC-W), which measures the duration of the cell’s passage through the laser beam, indicating size variability and aiding in the differentiation between individual cells and aggregates.

FSC-A, FSC-H, FSC-W, as well as the fluorescence of CV, TO, and DRAQ-5, are measured with the LSRII flow cytometer. To exclude aggregates and debris we set an FSC-A/FSC-H gate (R1, Figure 6A). To focus on the erythrocytic cells (RBCs and reticulocytes), we set gates on the dim or negative cells in TO vs CV (R2, Figure 6B) and TO vs DRAQ-5 (R4, Figure 6C) plots. R4 is a contour gate including 99% of events. We combine the gates (Boolean AND) R1, R2, R4 and exclude R3 (Boolean NOT) that has been set on the TO^+^ reticulocyte population. See <u>Results</u>^17^ and the legend to Figure 6 for additional details on analysis.

##### Imaging flow cytometry (IFC)

The ImageStreamX analyzes individual cells—in flow, by brightfield and multiple fluorescence parameters—as they flow past a 60× microscope objective. The device collects data as images, namely, two-dimensional spatial grids, with a third intensity dimension captured for each pixel. We load RBCs with CV (for viability), TO (for RNA), and DRAQ-5 (for DNA), as described <u>above</u> ^18^, and establish gating schemes in order to discriminate mature RBCs from other cell classes, measure Ø_Major_ for biconcave disks and non-biconcave disks, and determine the proportion of the RBCs that are BCDs vs. nBCDs (see <u>Results</u>^19^).

Gating schemes (e.g., see Figure 7A) that we establish allow size analysis of individual RBCs and RBC precursor types, separately from one another. Measurements of Ø_Major_ are steps #6 (for BCDs) and #6′ (for nBCDs) in our workflow (see Figure 1), and determinations of prevalence are steps #15 (for BCDs) and #15′ (for nBCDs).

##### Use of inhibitors in imaging flow cytometry

As detailed in <u>paper #1</u>^20^, we collect fresh blood from WT or KO mice, centrifuge, remove the buffy coat, suspend the pelleted RBCs in our oxygenated solution (see Table 1) to a hematocrit (Hct) of 5% to 10%, and then centrifuge at 600 × g for 5 min (see <u>above</u>^21^). After two such washes, we resuspend RBCs in oxygenated solution without drugs or with either pCMBS (1 mM for 15 min) or DIDS (200 µM for 1 hour), to a final Hct of 25%. After centrifuging RBCs at 600 × g for 5 min, we aspirate and discard the resulting supernatant to remove inhibitors, and load the blood cells with CV, TO and DRAQ-5, as described <u>above</u>^18^.

After implementing gating schemes (Figure 7A) to sort mature RBCs from their precursors, we then sort the normal BCD RBCs from the nBCD cells, employing the Lobe count feature in the Ideas software, using the H Variance Mean function (granularity assigned as 5). This sorting procedure yields four bins of cells identified as possessing 1, 2, 3, or 4 lobes. Regardless of the lobe bin (i.e., 1, 2, 3, or 4 lobes), the majority of nBCD cells have H Variance Mean values <10.

We manually inspect each cell in each bin of sorted cells to verify correct assignment of BCD (i.e., normal) vs. nBCD cells. We performed experiments on blood samples from three age-matched pairs of WT vs dKO mice.

#### Proteomic analysis

##### Preparation of RBC ghosts

We collect blood samples (∼250 μl) into heparinized microcentrifuge tubes from three mice for each genotype, and then place the tubes on ice, pending immediate processing into ghosts. We generate erythrocyte ghosts as previously described (Bennett, 1983), with the following changes to buffer composition (see Table 1): RBC lysis and post-lysis wash buffers contain 5 mM Tris-HCl/pH 8.0 and complete Protease Inhibitor Cocktail tablets (cPIC; Roche, 04693116001), as opposed to sodium-phosphate buffer/pH 7.5 and PMSF and pepstatin A—modifications required to obtain good mass-spectrometry data. Ghosts are then flash-frozen and held at −80 °C prior to mass-spectrometry analysis.

##### Mass spectrometry

We perform mass spectrometry experiments at the CWRU Center for Proteomics and Bioinformatics. Briefly, RBC ghost samples are lysed with 2% SDS and cPIC, using pulse sonification, and removing SDS by filter-aided sample preparation (Wiśniewski, 2017) detergent cleanup. We determine total protein concentration using the Bio-Rad Protein assay kit (Bio-Rad, Hercules CA), digesting a 10-µg sample with LysC/Trypsin, and analyzing 300 ng of the resulting product via 24-hr label-free liquid chromatography–tandem—mass spectrometry (LC/MS/MS) using a UHPLC Vanquish LC system and a Orbitrap Exploris 480 Mass Spectrometer (TFS) following a previously described protocol (Schlatzer et al., 2009). We extract raw data from each run to generate MS/MS peak lists for identification purposes, as well as intensity-based profile peak lists for quantification using Peaks software (Bioinformatics Solutions, Ontario Canada). We perform one-way ANOVA to test for statistically significant differences in abundance (see <u>below</u>^22^).

##### Relative abundance

We determine relative abundance by comparing the mean area under curve (AUC) for all detected peptides for each protein, from three mice of each genotype. For example, with anion exchanger 1 (AE1; band 3; SLC4A1), we detect 57 different peptides. Therefore, in samples from three animals, the relative abundance of AE1 is the mean AUC of the 57 × 3 = 171 total peptides.

### Statistical analysis

We report results as mean ± SD. In each figure legend, we report which statistical tests is performed, from among the following, to generate unadjusted *P*-values: (1) paired two-tailed t-test, (2) unpaired two-tailed Welch’s t-test (Welch, 1947) or (3) one-way analysis of variance (ANOVA). For comparisons of two means, we performed #1. For comparisons between more than two means, we performed #2 or #3 and then, to control for type-I errors across multiple means comparisons, we applied the Holm-Bonferroni correction, setting the familywise error rate (FWER) to α = 0.05. Briefly, we order the unadjusted *P*-values for all N comparisons in each dataset from lowest to highest. For the first test, we compare the lowest unadjusted *P*-value to the first adjusted α value, α/N. If the null hypothesis is rejected, then we compare the second-lowest *P*-value to the second adjusted α value, α/(N–1), and so on. If, at any point, the unadjusted *P*-value is ≥ the adjusted α, the null hypothesis is accepted and all subsequent hypotheses in the test group are considered null.

#### Data availability

The data supporting the findings of this study are available within the paper and its Supplementary Information files. Any further relevant data are available from the corresponding author upon reasonable request.

## Results

As described in <u>Methods</u>^23^, we prepared RBCs from WT, AQP1-KO, RhAG-KO, and dKO mice to assess the impact of each genetic deletion on RBC shape, other morphometric parameters, and the RBC-membrane proteome. This approach allows us to correlate physiological data (i.e., SF) and automated-hematology results from <u>paper #1</u>^24^ with mathematical modelling/simulations from <u>paper #3</u>^25^, and thereby estimate *P*_M,O_2__.

### Morphometry

#### Blood smears

##### Effect of gene deletions

Expert review^26^ of 12 blood smears for control cells (i.e., not treated with drugs) reveals unremarkable RBC morphology (Figure 2) of each of four genotypes: WT, *Aqp1*–/–, *Rhag*–/–, and *Aqp1*–/–/*Rhag*–/–, all on a C57BL/6_Case_ background. We observed only occasional target cells and stomatocytes, with no differences among genotypes. Previous authors noted normal RBC morphology for RhAG-KO mice (Goossens *et al*., 2010).

**Figure 2.**
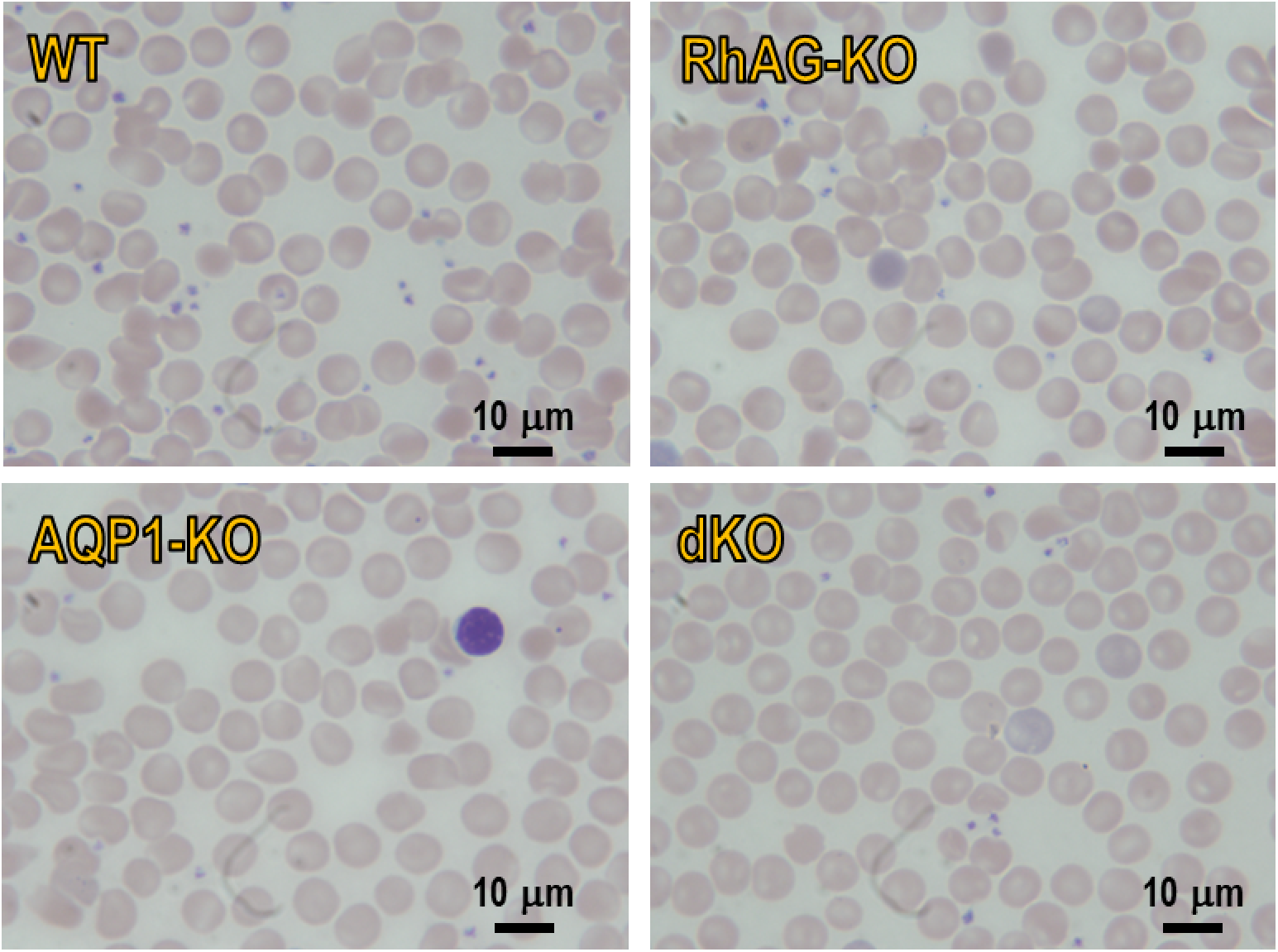
Representative blood smears from each of four genotypes. We generated 12 Wright-Giemsa stained air-dried peripheral blood smears from WT, AQP1-KO, RhAG-KO and dKO mice (i.e., 4 smears from each of 3 separate mice), for a total of 48 smears. No differences in red cell morphology are noted among knockout strains and WT mice. Photos are taken at 1000× magnification.

##### Effect of drugs

Previous reports indicate that exposing RBCs to pCMBS does not change RBC shape (Kuchel *et al*., 1997), whereas DIDS exposure can cause alterations in cell morphology (Mosior *et al*., 1992; Blank *et al*., 1994; Hoefner *et al*., 1994; Al-Samir *et al*., 2025).

In the present study, in which we evaluated RBCs from WT and dKO mice, we determine (1) whether pCMBS or DIDS impact morphology of RBCs used in our SF experiments in <u>paper #1</u>^27^, and (2) the extent to which drug-induced shape changes correlate with WT vs. dKO mice.

At the level of blood smears, we observe no remarkable drug-induced changes in RBC morphology for WT or dKO (Figure 3). In the particular case of pCMBS-treated cells, we note a faint halo of light-purple around cells. Occasionally, we observed vacuoles at the periphery of RBCs—outside of the cell membrane—in some of the pCMBS-treated samples (Figure 3, center panels). Expert review deems these changes to be artifactual, and likely to be related to the effect of the added pCMBS on the chemistry of the staining reactions.

**Figure 3.**
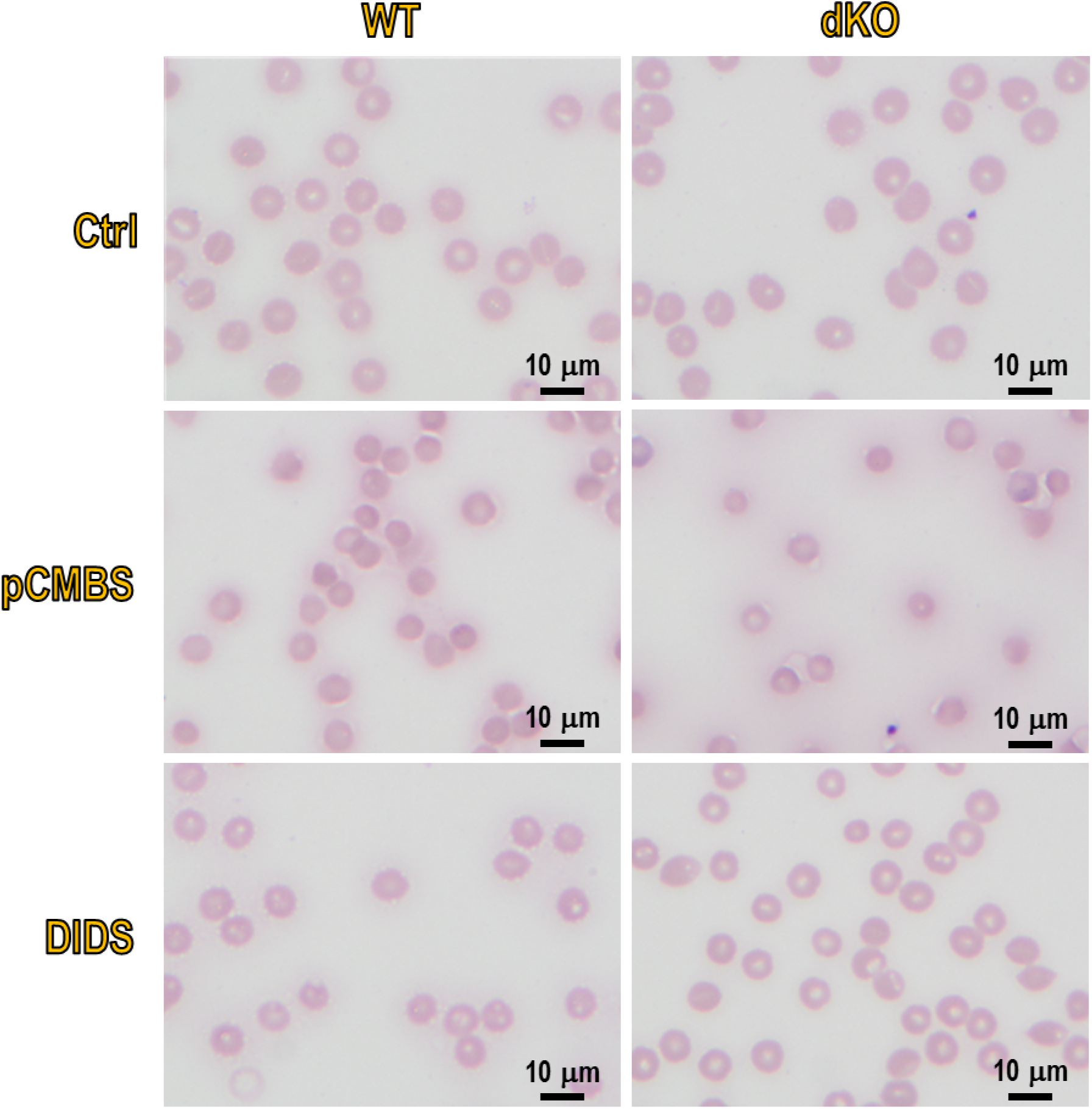
Representative blood smears from WT and dKO mice, of cells treated with inhibitors. Representative images (1000× magnification) of Wright-Giemsa stained air-dried peripheral blood smears using blood obtained from WT mice (left column row) and dKO mice (right column), either in control conditions (Ctrl; top row) or with a 15 min pretreatment with 1 mM pCMBS (middle row) or a 1 h pretreatment with 200 μM DIDS (bottom row). No significant differences in red cell morphology were noted between WT and dKO mice under any tested conditions.

#### Imaging of living, tumbling RBCs

The preceding analyses were performed on air-dried blood smears. In order to evaluate RBC morphology in an environment more closely approximating that of the SF reaction cell during *k*_HbO_2__ experiments, we also use DIC microscopy to observe living RBCs—suspended in our oxygenated solution (see Table 1) at an initial Hct of 25–30%—as we drop the RBC suspension onto glass coverslips overlaid with the same oxygenated solution (for a final Hct of 0.5–1%; see <u>Methods</u>^28^). We record both still and video micrographs of living RBCs as, driven by gravity, as they tumble through the focal plane and toward the coverslip surface, specifically avoiding acquisition of images at the coverslip surface because many cells quickly deform on contact with the acid-washed glass.

##### Effect of gene deletions

Still images from DIC microscopy (Figure 4) reveal that virtually all cells, regardless of the genotype of the donor mouse, are normal BCDs. In addition, videos of living RBCs tumbling through our oxygenated solution (see Table 1)—see Video 1, Video 2, Video 3, and Video 4—confirm that nearly all RBCs of all four genotypes are normal BCDs.

**Figure 4.**
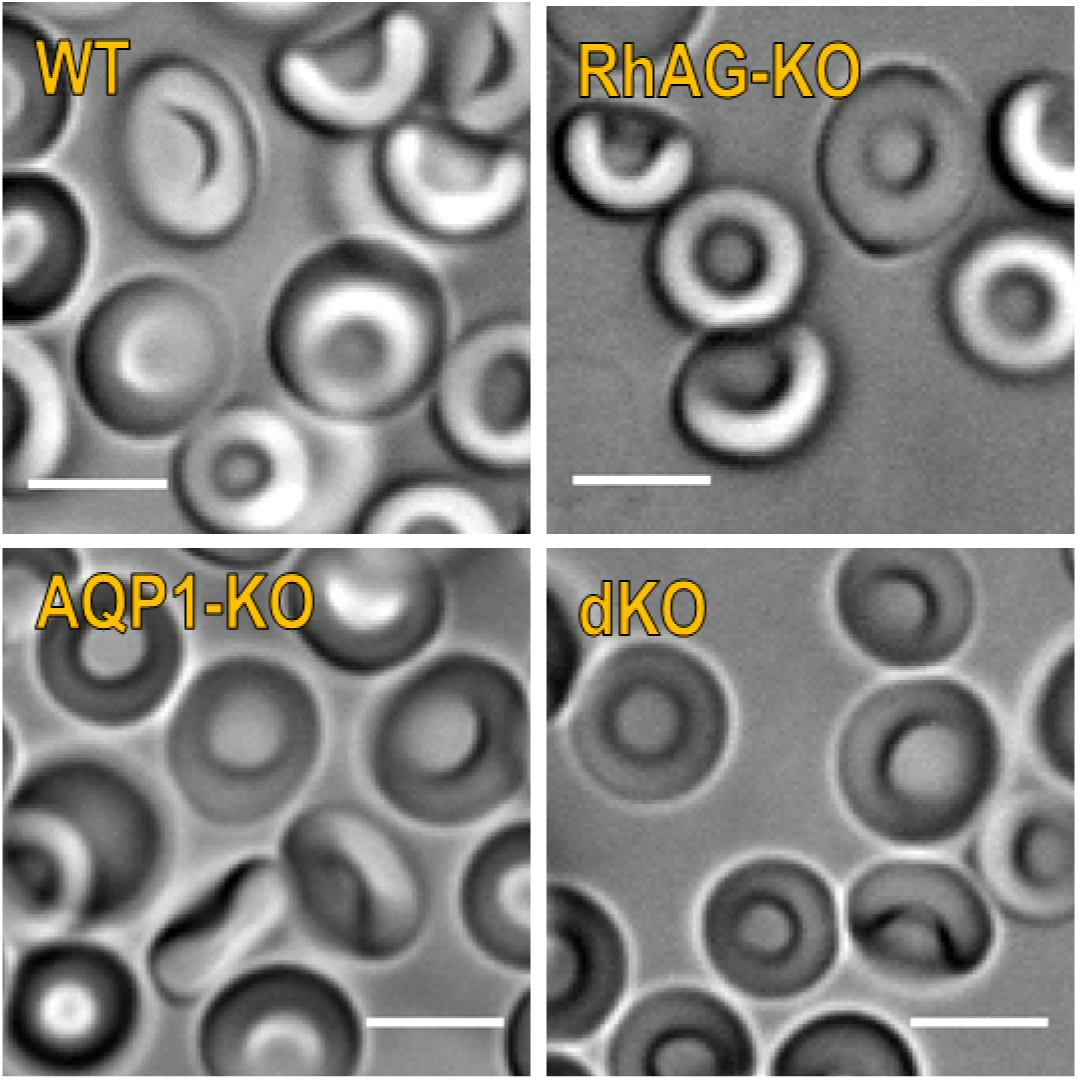
Representative DIC still micrographs of living RBCs. DIC micrographs of fresh RBCs from WT and the three KO mouse strains (annotated in the top left corner of each panel), tumbling through plane of focus and revealing normal biconcave disks. On a given day, we reviewed 4 to 5 samples (i.e., droplets) of RBCs suspended in oxygenated solution for 1 mouse of each genotype. We executed this protocol on 3 separate days, for a total of 3 separate mice/genotype (i.e., a total of 12 to 15 samples/genotype). Red cell morphology is similar in all groups and is unremarkable, with no differences noted among knockout strains and WT mice. The bars represents 5 μm. See (Supporting Videos Supporting video 1, Supporting video 2, Supporting video 3, and Supporting video 4) for a representative DIC video clip of each genotype (i.e., WT, AQP1-KO, RhAG-KO, dKO); each clip shows RBCs tumbling through the plane of focus.

##### Effect of drugs

DIC microscopy of inhibitor-treated RBCs (Figure 5 reveals that pCMBS pretreatment causes the occasional appearance of small, more spherical cells with spicules (Figure 5; yellow arrows), which we classify as nBCDs. The nBCDs were more prevalent in DIDS-treated than in pCMBS-treated cells.

**Figure 5.**
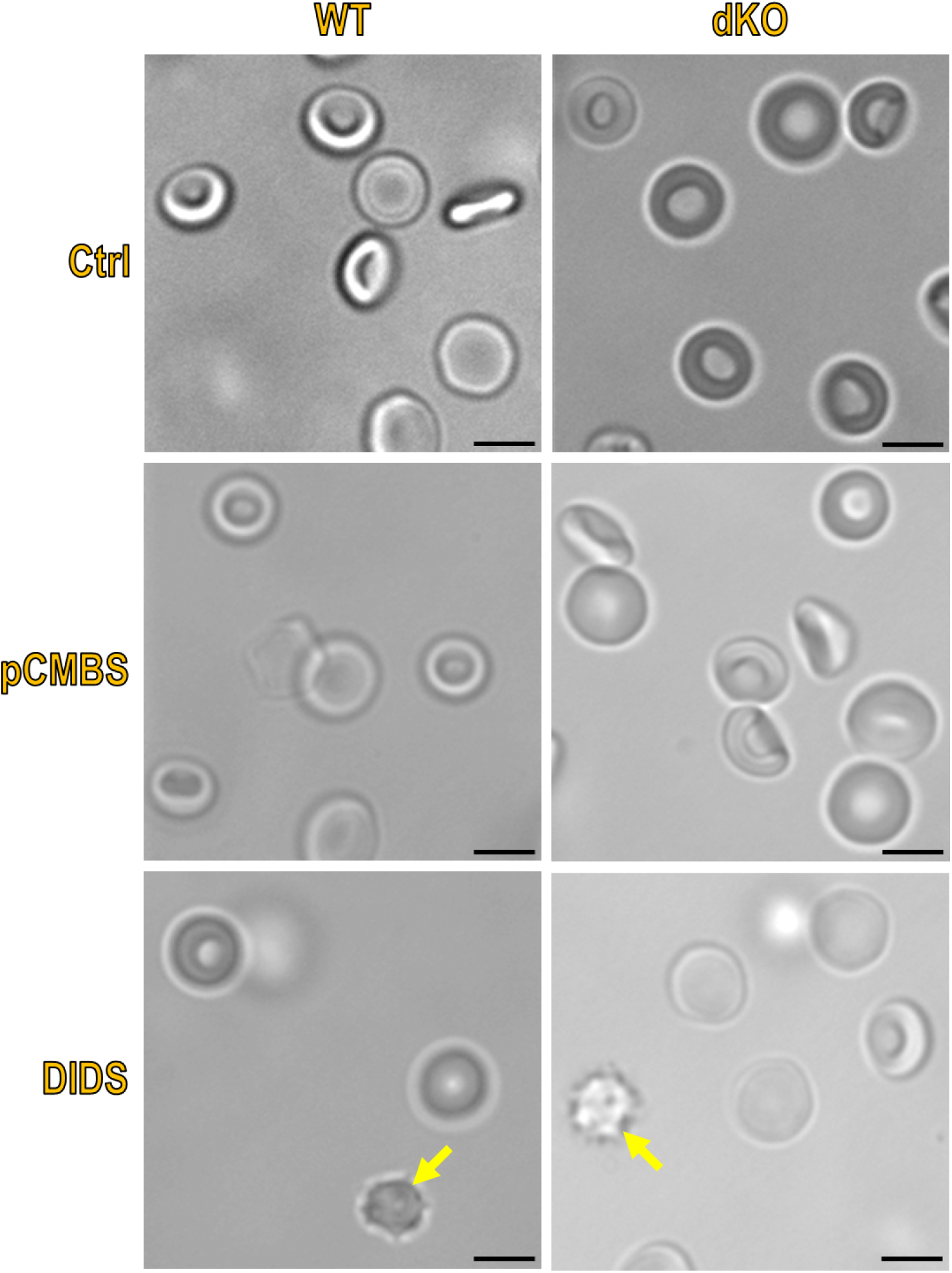
Representative DIC still micrographs of living RBCs, treated with inhibitors. On a given day, we reviewed four to five samples (i.e., droplets) of RBCs suspended in oxygenated solution for one mouse of each of genotype. We executed this protocol on three separate days, for a total of three separate mice/genotype (i.e., a total of 12 to 15 samples/genotype). Red cell morphology is similar in control groups and is unremarkable, with no differences noted among dKO and WT mice. For both WT and dKO mice, treatment of RBCs with inhibitors tended to cause the appearance of small, spherical cells with spicules (yellow arrows), more so in DIDS-treated than in pCMBS-treated cells. We can also identify these tumbling spherocytes by microvideography (not shown). The scale bars represents 5 μm.

Note that, in SF experiments on drug-treated cells, the RBCs usually entered the SF device within 5 min of the completion of the pretreatment. For DIC experiments, this delay was typically 30-90 min. Thus, to the extent that formation of nBCDs is time dependent, the nBCD abundance observed by DIC would overestimate the actual abundance in SF experiments.

#### Light-scattering flow cytometry

Flow cytometry provides a more efficient and less subjective approach than microscopy for assessing the morphology of large numbers of living cells in an environment that resembles the SF reaction cell. In light-scattering flow cytometry, the amount of light scattered depends on both the size and internal intricacy of a single cell as it passes through the laser beam. Simultaneous forward and side scatter have long been used in flow cytometry to discriminate among RBCs and various type of leukocytes.

##### Gating

To discriminate RBCs from very small particles on the one hand and aggregates on the other, we gate the light-scattering data—representing a total of >500,000 cells per run—by plotting FSC-A vs FSC-H. In Figure 6A, the thin cluster of gray points nearest the origin y-axis represents particulates (“P”), whereas the larger, diffuse cluster of gray points centered around FSC-A ≅ 10^−5^ represent aggregates (“A”). We discriminate among RBCs, reticulocytes, and nucleated precursors by using gating schemes based upon staining with the fluorescent markers CV, DRAQ-5 and TO (see <u>Methods</u>^29^). The tear-drop area (“R1”) comprises the “gated” cells. For all four genotypes at least 99.9% of the cells within “R1” are CV positive (i.e., viable). As described next, the plots in Figure 6B and Figure 6C allow us to identify (1) reticulocytes (arbitrarily-colored red pixels) and (2) nucleated precursors or other nucleated contaminant cells (arbitrarily-colored cyan pixels). We show these red and cyan pixels (retroactively classified) in Figure 6A—all in area “R1”—as well as in panels *B* and *C*.

**Figure 6.**
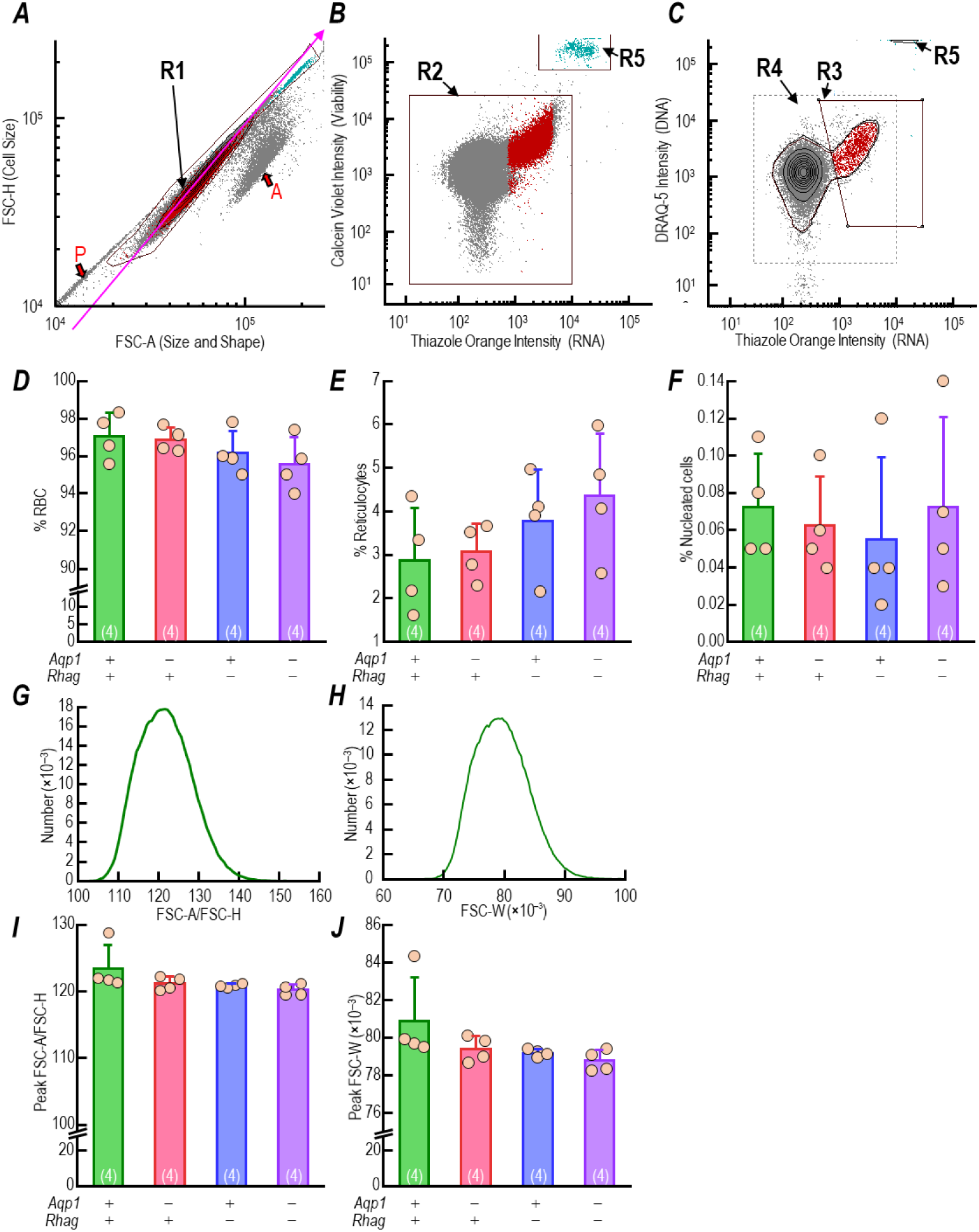
Light-scattering flow cytometry. We sort mature RBCs from their precursors in the LSRII device in order to perform size and shape analyses. In panels A-C and G and H we display representative data from one WT sample. For definitions, see Methods^52^. *A,* Singlet cells in the R1 region of the FSC-A/FSC-H plot are individual viable blood cells. An arrow pointing from the red letter “P” annotates the gray coordinates representing small particulates and the arrow pointing from the red letter “A” annotates coordinates for cell aggregates. The red and dark-cyan colored pixels represent reticulocytes (identified by the R3 region, panel *C*) and nucleated cells (identified by the R5 region, panel *B & C*) respectively, and the coloring is applied using Boolean logic) after the regions in panels *B* and *C* are applied. *B,* The cells within R1 from panel *A* are plotted in a TO vs. CV intensity plot confirming that all are intact and alive. Two distinct clusters are observed and are set as the regions R2 and R5. R5 included cells with high TO and CV intensity are also observed in Panel *C* and classified there. *C,* When analyzing the R1 cells on a TO vs. DRAQ-5 intensity plot we see the R5 cluster also has the highest TO^+^ and DRAQ-5^+^ signals. These are the CV^+^ nucleated cells that are colored dark-cyan in Panels *A-C*. A region (R3) isolates reticulocytes (red pixels) within R4. The R4 contour captures 99% of the enucleated cells, representing the erythrocytic populations (RBCs and reticulocytes). Cells in panel *B* were already gated in panel *A* (i.e., they are all single cells). The mature RBCs are both TO^−^ and DRAQ-5^−^. Reticulocytes (red points) are TO^+^ but DRAQ-5 ^dim^. Nucleated cells are the brightest TO^+^ and DRAQ-5^+^ cells. . *D,* Summary of the mean %RBCs from 4 samples each of WT, AQP1-KO, RhAG-KO and dKO blood. The break in the y-axis is from 12-89%. We measure no significant differences between genotypes. All *P*-values for one-way ANOVA, followed by the Holm-Bonferroni correction (see <u>Methods</u>^35^) are presented in Statistics Table 1. *E,* Summary of the mean %reticulocytes from 4 samples each of WT, AQP1-KO, RhAG-KO and dKO blood. We measure no significant differences between genotypes. All *P*-values for one-way ANOVA, followed by the Holm-Bonferroni correction are presented in Statistics Table 2. *F,* Summary of the mean %nucleated cells from 4 samples each of WT, AQP1-KO, RhAG-KO and dKO blood. We measure no significant differences between genotypes. All *P*-values for one-way ANOVA, followed by the Holm-Bonferroni correction are presented in Statistics Table 3. In Panels *D-F*, mouse genotype is indicated by a + or – for *Aqp1* and *Rhag* below each of the columns, which are colored green for WT, pink for AQP1-KO, blue for RhAG-KO and purple for dKO. Individual data points from each sample are plotted on top of the columns and error bars represent SD. *G,* Representative RBC-only Boolean-gated single parameter frequency distribution of forward-scatter intensity area (FSC-A)/forward scatter pulse height (FSC-H; see <u>Methods</u>^52^ from the WT sample in panel *A* (This plot is the equivalent of traveling up the (magenta) diagonal arrow of the FSC-H v FSC-A plot in panel *A* with the nucleated cells (R5) and reticulocytes (R3) excluded. *H,* Representative RBC-only Boolean-gated single parameter frequency distribution of forward-scatter intensity width (FSC-W; see <u>Methods</u>^52^) from the WT sample in panel *A*. *I,* Summary of the mean peak FSC-A/FSC-H ratio from 4 samples each of WT, AQP1-KO, RhAG-KO and dKO blood. Individual data points from each sample are plotted on the bars. The break in the y-axis is from 28-99. We measure no significant differences between genotypes. All *P*-values for one-way ANOVA, followed by the Holm-Bonferroni correction are presented in Statistics Table 4. *J,* Summary of the mean peak FSC-W value from 4 samples each of WT, AQP1-KO, RhAG-KO and dKO blood. The break in the y-axis data is between 28×10^−3^ to 75×10^−3^. We measure no significant differences between genotypes. All *P*-values for one-way ANOVA, followed by the Holm-Bonferroni correction are presented in Statistics Table 5. In Panels *G-J*, genotypes and columns are annotated and colored as in panels *D-E.* Individual data points from each sample are plotted on top of the columns and error bars represent SD. CV, calcein violet (viability); TO, thiazole orange (RNA); and DRAQ-5, Deep Red Anthraquinone 5 (DNA).

In Figure 6B, we plot CV intensity (i.e., viability) vs. TO intensity (i.e., RNA) for the cells from “R1” in Figure 6A. As noted above, >99.9% of the cells are above the x-axis and thus viable. Moreover, the cells colored red and cyan have relatively high RNA levels.

In Figure 6C, we plot DRAQ-5 intensity (i.e., DNA) vs. TO for the “R1” cells in Figure 6A. The small cluster of cyan pixels in the upper right of the plot are nucleated (DRAQ-5^+^) cells and are gated as “R5” in panels *B* and *C* because they are not only viable, but have strong TO signals (i.e., they have endoplasmic reticulum, ER) and DRAQ-5 signals (i.e., they have nuclei). Returning to Figure 6B, nearly all of the cells that are not “R5” are gated as “R2”. In Figure 6C, we then use Boolean gating to subtract the manually set region “R3” from the contour-fit of the gray pixels in “R4” (dashed box) to isolate the reticulocytes (red pixels), which are viable (see red pixels in Figure 6B), have high TO signals (i.e., they possess residual ER), but are DRAQ-5^dim30^ (i.e., they lack nuclei). The remaining cells are mature RBCs that lack both ER and nuclei, 99% of which are captured within the contour-fit within “R4”.

Using these criteria we conclude that 97.1 ± 1.2% of the cells in the samples from the 4 WT animals in Figure 6D (green column) are mature RBCs. Figure 6D also shows that the mean %RBC in AQP1-KO, RhAG-KO, and dKO samples are, respectively, 96.9 ± 0.6% (pink column), 96.2 ± 1.2% (blue column) and 95.6 ± 1.4% (purple column). A statistical analyses reveals no significant differences between WT and any of the three KO strains for the percentages of mature RBCs (Figure 6D), reticulocytes (Figure 6E), or nucleated cells (Figure 6F).

##### Size and shape analysis

We next confine our light-scattering analyses to include only the gated, mature RBC population (as defined in Figure 6A-C). Our goal is to identify any unexpected strain-dependent changes in size and/or shape that could potentially impact *k*_HbO_2__, independent of *P*_M,O_2__. Figure 6G is a representative Boolean-gated single-parameter frequency distributions of FSC-A/FSC-H from the single WT sample in Figure 6A. The distribution in Figure 6G is equivalent to passing along the magenta diagonal arrow through the mature RBC coordinates within “R1” of the FSC-A vs. FSC-H plot in Figure 6A. Figure 6H is an analogous frequency distribution for FSC-W.

We performed a total of 16 experiments like that shown in Figure 6G (4 mice/strain × 4 strains). Figure 6I summarizes the FSC-A/FSC-H ratios at which peak values were achieved in these experiments. A statistical analysis reveals no significant differences among the four genotypes. We also performed a total of 16 experiments like that shown in Figure 6H, and Figure 6J summarizes the FSC-W at which peak values were achieved. Again, a statistical analysis reveals no significant differences among the four genotypes. From panel *I* and *J*, we conclude that RBCs from WT, AQP1-KO, RhAG-KO, and dKO mice have similar shape distributions. The small sample-to-sample variations among replicates (from one genotype) may be due to flow nuances (e.g., slight variation in flow rates), or environmental variables (e.g., small differences in cell density).

#### Imaging flow cytometry (IFC)

Whereas the light-scattering/fluorescence experiments, just discussed, provide a semiquantitative measure of cell size and shape (FSC-A), cell size (FSC-H), size variability (FSC-W), and cell classification (gated by fluorescent markers), IFC provides similar basic information as well as the high-resolution, multi-spectral imaging capability (see <u>Methods</u>^31^). As a result, we can observe the size and shape characteristics of individual RBCs, in flow. Similar to light-scattering flow cytometry, IFC allows the sorting (i.e., gating) of multiple cell classes based upon combinations of fluorescent markers, but adds concurrent brightfield image acquisition of individual cells as they pass a 60× microscope objective lens. Thus, the investigator can analyze cell size and shape, with substantially increased throughput and considerably reduced opportunity for subjective bias during analysis versus DIC microscopy, micrographs, or videography.

##### Effect of gene deletions

Using the same staining protocols as for the light-scattering flow cytometry (see <u>Methods</u>^32^), we label blood samples with CV, TO and DRAQ-5, and then establish gating schemes to sort the mature RBCs from reticulocytes, and nucleated precursors or other nucleated contaminant cells. Figure 7A is a frequency histogram for a one of four IFC runs on WT material, representing all CV^+^ events (i.e., viable cells), plotted as a function of TO (i.e., RNA) intensity. We represent the TO^+^ events—about 17,000 (i.e., ∼4.5% of the total events), distributed across a span of 999 bins along the x-axis—as peach dots and line segments. The ∼360,000 (i.e., >95% of the total events) TO^−^ events fall within a single bin (green).

**Figure 7.**
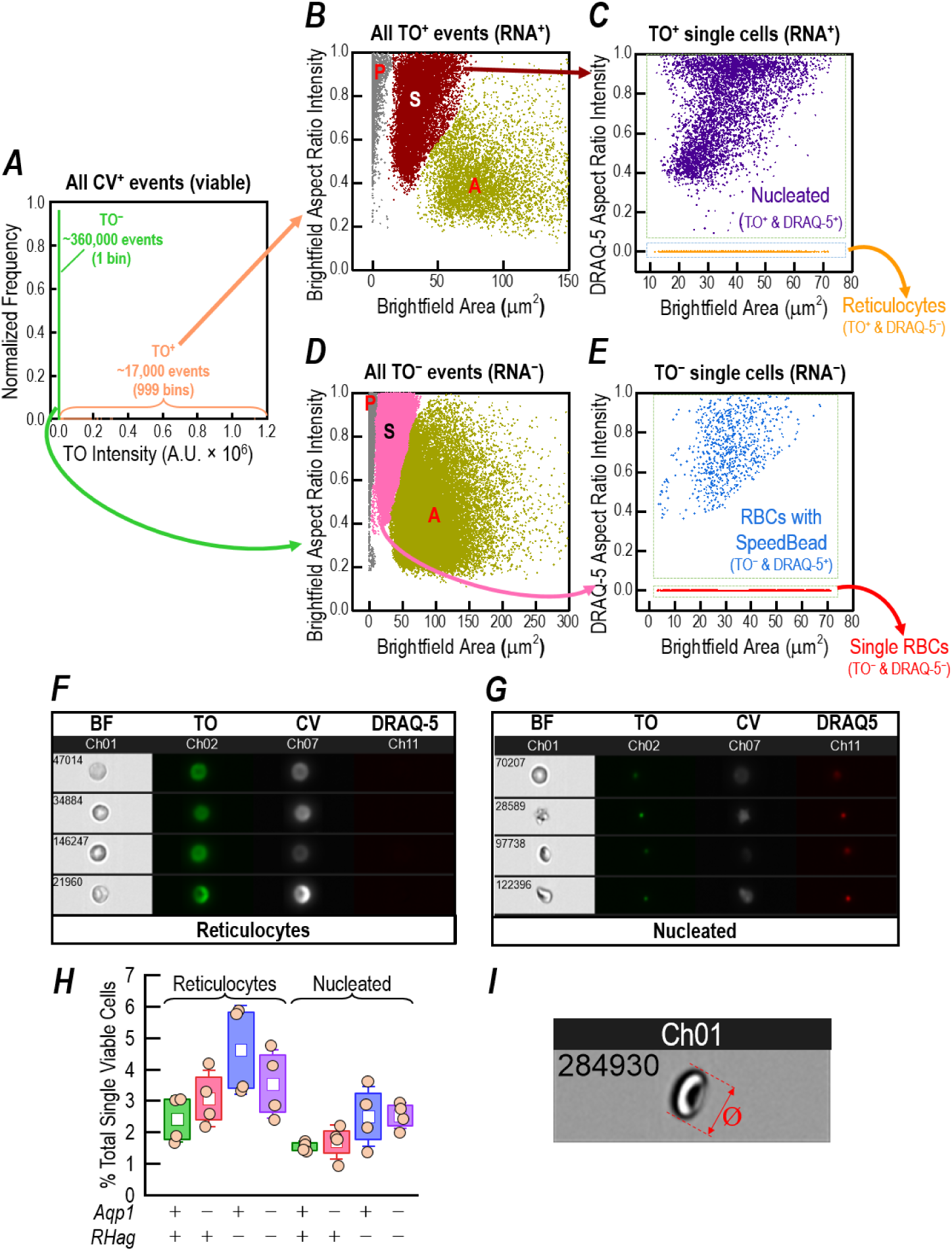
Imaging flow cytometry (IFC) IFC gating schemes established to allow analysis of individual mature RBCs and other non-mature RBC types. We sort mature RBCs from their precursors and nucleated cells by IFC in order to perform further size and shape analyses. We begin in panel *A,* with all CV-positive cells (CV^+^) and within this population distinguish TO^+^ (peach) from TO^−^ cells (green). In panel *B*, which plots the brightfield area vs. the brightfield aspect ratio of the TO^+^ cells sorted from panel *A,* we isolate the single TO^+^ cells (S; burgundy data points) from small particles (P; gray data points) or aggregated cells (A; drab-olive data points). In panel *C,* plots of the brightfield area vs. DRAQ-5 aspect ratio intensity for the single TO^+^ cell population from panel *B*, further isolates the nucleated TO^+^ and DRAQ-5^+^ cells (purple data points) from reticulocytes (TO^+^ DRAQ-5^−^; orange data points). In panel *D*, which plots the brightfield area vs. the brightfield aspect ratio of the TO^−^ cells sorted from panel *A,* we isolate the single TO^+^ cells (S; pink data points) from small particles (P; gray data points) or aggregated cells (A; drab-olive data points). In panel *E,* plots of the brightfield area vs. DRAQ-5 aspect ratio intensity for the single TO^−^ cell population from panel *D*, further isolates the TO and DRAQ-5^+^ population that represents RBCs with a contaminating speed bead in the same image (blue data points) from single mature RBCs (TO and DRAQ-5^−^; red data points). Panel *F* displays example images of cells classified as reticulocytes according their CV^+^, TO^+^ and DRAQ-5^−^ fluorescent signature. Panel *G* displays example images of cells classified as nucleated according their CV^+^, TO^+^ and DRAQ-5^+^ fluorescent signature, which have a heterogeneous morphology. In both panels *F* and *G* the channels are assigned in order of increasing fluorescence excitation wavelength.*H,* Percent of total cell count represented by cells not defined as mature RBCs for each genotype, identified by TO and DRAQ-5 staining as in Panel *C*. Each mouse genotype is color-coded green (WT), pink (AQP1-KO), blue (RhAG*-*KO), or purple (dKO). The presence (+) or absence (–) of *Aqp*1 or the *RhAG* gene is annotated at the base of each box plot. The cell classification is annotated above the braces grouping each set of four box plots. The mean for each group is plotted as an open square, the boxes represent the interquartile range, and the whiskers represent SD. The four individual data points (beige circles) overlaying each box pot report the normalized percentage total cells represented by each cell classification in a single blood sample. One-way ANOVA, followed by the Holm-Bonferroni (Holm, 1979) correction (see <u>Methods</u>^35^) determines that there are no significant differences vs. WT (see Statistics Table 7 and Statistics Table 8). *I,* After gating in IFC according to their fluorescence signature (Panels *A*→*E*), the Ø_Major_ of RBCs and non-RBCs were determined by measuring the length of the longest axis (red box) of the cells in the corresponding BF images. Every cell is annotated with a unique numeric identifier in the top left of the image. In *A*–*E*, CV, calcein violet (viability); TO, thiazole orange (RNA-stain); DRAQ-5, Deep Red Anthraquinone 5 (DNA-stain). BF, brightfield image.

In Figure 7B, we analyze just the TO^+^ events by plotting projection area of the imaged event vs. bright-field aspect-ratio intensity^33^. The burgundy events represent single TO^+^ cells (“S”). The gray events in the upper left are particulates (“P”), whereas the dull-olive events to the center/right are aggregates (“A”).

In Figure 7C, we analyze just the single cells that are TO^+^ by plotting the brightfield area of the imaged event vs. DRAQ-5 aspect ratio intensity^34^. The events colored purple are single TO^+^ (RNA^+^)/DRAQ-5^+^ (DNA^+^) cells that could be nucleated RBC precursors or other nucleated cells (e.g., contaminating white blood cells), neither of which would contain hemoglobin (i.e., they would not affect SF experiments in paper #1). The orange events that appear to from a line at y ≅ 0 are TO^+^ (RNA^+^) but DRAQ-5^−^ (DNA^−^). These are reticulocytes.

Returning to Figure 7A, we now analyze just the TO^−^ (RNA^−^) events (green) by plotting in Figure 7D brightfield area (analogous to Figure 7B) vs. the brightfield aspect ratio intensity. The events colored pink in Figure 7D represent TO^−^ single cells (“S”) that lack RNA (i.e., RBCs). The gray events in the upper left are TO^−^ particulates (“P”), whereas those colored dull-olive to the center/right are TO^−^ aggregates (“A”).

Finally, in Figure 7E, we analyze just the single cells that are TO^−^ (RNA^−^) by plotting the fluorescence aspect ratio intensity vs. the brightfield area of the imaged event. The blue-colored events represent single, mature RBCs in which a SpeedBead™ appears in the same image. Finally, the red events that appear to form a line at y ≅ 0 represent single, mature RBCs that are uncontaminated by SpeedBeads™.

Of the ∼375,000 images that generate the data for the WT sample in Figure 7A*–E*, Figure 7F*–G* are exemplars of two classes, reticulocytes and nucleated cells. Figure 7F is a data collage of four reticulocytes, and Figure 7G is the same for nucleated cells. In each panel, each of the four data rows contains comprises a brightfield image of the cell in flow, a TO image (green, RNA^+^), a CV image (gray, viable), and DRAQ-5 image (red, DNA^+^).

The data summarized in Figure 7A*–E* are one of four IFC runs on WT mice. The top data row in Table 2 summarizes data for all four WT runs—the abundance of single mature RBCs, reticulocytes, and nucleated cells.

**Table 2.** Prevalence mature erythrocytes vs. reticulocytes and nucleated cells present in mouse blood, determined using IFC.

| Mouse | % Mature RBCs |  | % Reticulocytes |  | % Nucleated cells |  |
| --- | --- | --- | --- | --- | --- | --- |
|  | Mean | SD | Mean | SD | Mean | SD |
| **WT** | 96.06 | 0.58 | 2.40 | 0.73 | 1.54 | 0.16 |
| **AQP1-KO** | 95.25 | 0.67 | 3.06 | 0.88 | 1.69 | 0.54 |
| **RhAG-KO** | 92.90 | 0.66 | 4.60 | 1.41 | 2.50 | 0.96 |
| **dKO** | 93.94 | 1.46 | 3.53 | 1.10 | 2.53 | 0.41 |

Data rows 2–4 in Table 2 summarize comparable data for the three KO strains. We see a trend for the abundance of mature RBCs to slightly decrease in the order WT > AQP1-KO > dKO > RhAG-KO with an almost reciprocal increase in the reticulocyte percentages in the order WT < AQP1-KO < dKO < RhAG. We also see small increases in the percentage of nucleated cells in the order WT < AQP1-KO < RhAG-KO < dKO. The box-plot in Figure 7H displays in more detail the individual results from each of the four runs for each genotype summarized in Table 2. Figure 7H highlights the trend for increased reticulocyte (left-grouped box plots) or nucleated cell (right-grouped box plots) percentage in each of the KO strains. However, the increases neither in the percentage of reticulocytes nor in the proportion of nucleated cells in KO blood are statistically significant compared with WT. Only if the minor increases in the reticulocytes and nucleated cells are combined do the slight reciprocal decreases in RBC prevalence become statistically significant in RhAG-KO or dKO vs. WT IFC samples (Statistics Table 6).

The major advantage that IFC offers over conventional flow cytometry is the opportunity for the user, examining brightfield images, to assess individually the morphology of hundreds of thousands of RBCs and directly measure their Ø_Major_ (Figure 7I, red box). We find that Ø_Major_ is slightly smaller in KOs than WT (Table 3) for either mature RBCs alone, or “all cell types” (i.e. mature RBCs + reticulocytes + nucleated cells). These results imply, together with increased MCVs for the KO strains (Table 4), that KO cells are slightly thicker. Due to the extremely high number of individual cells analyzed, one-way ANOVA with a Holm-Bonferroni means comparison (Holm, 1979; see <u>Methods</u>^35^) shows that differences between all pairs of mean Ø_Major_ values are statistically significant (*P*<0.001 for all comparisons). However, even the greatest difference (i.e., between WT and RhAG-KO) among only mature RBCs is ∼5% (i.e., 6.79 μm vs. 6.47 μm), and for “all cell types” (i.e., the sum of mature RBCs + reticulocytes + nucleated cells) is <5% (i.e., 6.80 μm vs. 6.53 μm). In our mathematical modeling in <u>paper #3</u>, we take as the diameter, the diameter of “all cell types.” This approach reflects the reality of an SF experiment (<u>paper #1</u>), which necessarily includes all cell types. Note that the inclusion of larger-diameter RBC precursor cells in the size analyses, does not substantially increase the mean major diameter of blood cells because these precursors represent only 4% (WT) to 7% (RhAG-KO) of the total cells in the sample.

**Table 3.** Quantification of major diameter (Ø_Major_) by IFC, among various blood-cell types, from four mouse genotypes*.

| Mouse | Cell types | n total | Mean ($\mu\text{m}$) | SD | Mode ($\mu\text{m}$) | Median ($\mu\text{m}$) | IQR |
| --- | --- | --- | --- | --- | --- | --- | --- |
| WT | Mature RBCs | 1156720 | 6.79 | 0.93 | 6.67 | 6.67 | 1.33 |
|  | Reticulocytes | 29317 | 7.06 | 1.15 | 6.67 | 7.00 | 1.33 |
|  | Nucleated cells | 18492 | 7.36 | 1.05 | 7.00 | 7.33 | 1.33 |
|  | Retics. + Nucleated | 47809 | 7.18 | 1.12 | 7.00 | 7.00 | 1.33 |
|  | ALL CELL TYPES | 1204530 | 6.80 | 0.94 | 6.67 | 6.67 | 1.33 |
| AQP1-KO | Mature RBCs | 1139890 | 6.68 | 0.90 | 6.67 | 6.67 | 1.33 |
|  | Reticulocytes | 36806 | 6.77 | 1.15 | 6.67 | 6.67 | 1.33 |
|  | Nucleated cells | 20044 | 7.14 | 1.01 | 7.00 | 7.00 | 1.00 |
|  | Retics. + Nucleated | 56850 | 6.90 | 1.12 | 6.67 | 7.00 | 1.33 |
|  | ALL CELL TYPES | 1196740 | 6.69 | 0.92 | 6.67 | 6.67 | 1.33 |
| RhAG-KO | Mature RBCs | 1077350 | 6.47 | 0.92 | 6.00 | 6.33 | 1.33 |
|  | Reticulocytes | 53283 | 7.18 | 1.22 | 7.00 | 7.00 | 1.67 |
|  | Nucleated cells | 29162 | 7.46 | 1.13 | 7.33 | 7.33 | 1.33 |
|  | Retics. + Nucleated | 82445 | 7.28 | 1.20 | 7.33 | 7.33 | 1.33 |
|  | ALL CELL TYPES | 1159800 | 6.53 | 0.97 | 6.00 | 6.33 | 1.00 |
| dKO | Mature RBCs | 1166840 | 6.52 | 0.92 | 6.33 | 6.33 | 1.00 |
|  | Reticulocytes | 44110 | 6.94 | 1.21 | 6.67 | 7.00 | 1.67 |
|  | Nucleated cells | 31462 | 7.32 | 1.04 | 7.33 | 7.33 | 1.33 |
|  | Retics. + Nucleated | 75572 | 7.10 | 1.16 | 7.00 | 7.00 | 1.33 |
|  | ALL CELL TYPES | 1242420 | 6.55 | 0.95 | 6.33 | 6.33 | 1.00 |

\*Blood samples were taken from four age-matched mice of each genotype. The $\emptyset_{\text{Major}}$ of all RBCs, reticulocytes (Retics.) and nucleated cells from blood of four strains of mice were analyzed after gating according to their fluorescence signature by IFC. One-way ANOVA with a Holm-Bonferroni means comparison (Holm, 1979) (see Methods) determined that differences between all pairs of mean $\emptyset_{\text{Major}}$ are statistically significant ( $P < 0.001$ for all comparisons). See Supporting file 2 for $P$ -values for all comparisons.

**Table 4.**
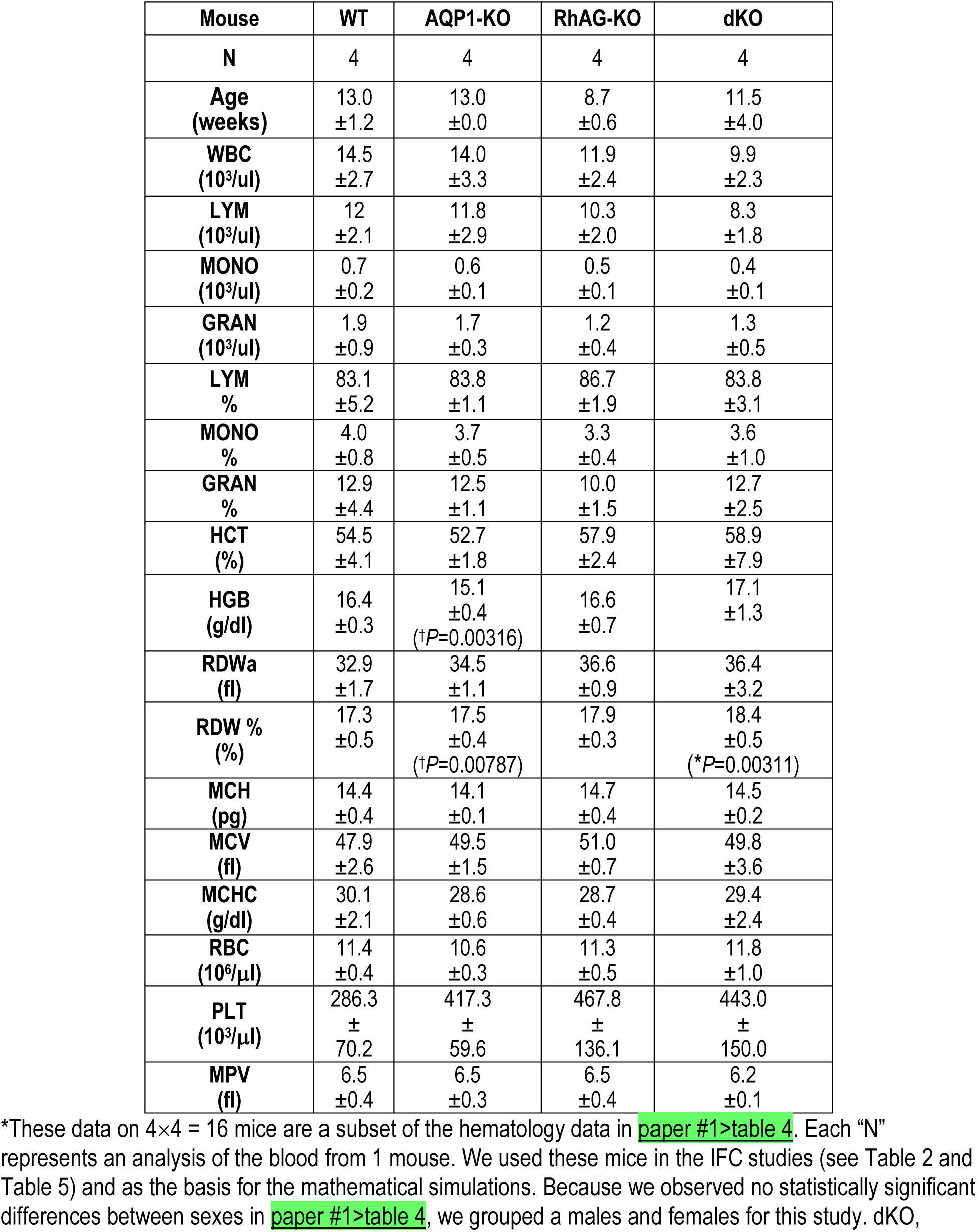
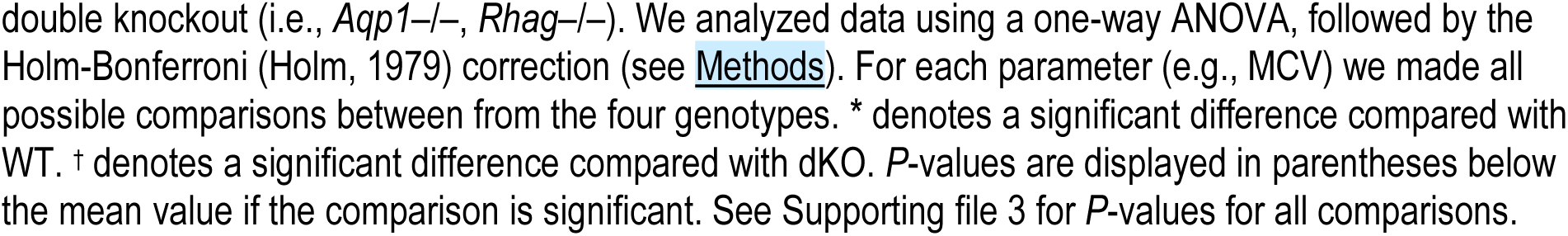
Summary of hematological data obtained from the subset of mice used in the IFC analysis and as basis for mathematical simulations*.

| Mouse | WT | AQP1-KO | RhAG-KO | dKO |
| --- | --- | --- | --- | --- |
| N | 4 | 4 | 4 | 4 |
| Age (weeks) | 13.0<br>±1.2 | 13.0<br>±0.0 | 8.7<br>±0.6 | 11.5<br>±4.0 |
| WBC (10<sup>3</sup>/ul) | 14.5<br>±2.7 | 14.0<br>±3.3 | 11.9<br>±2.4 | 9.9<br>±2.3 |
| LYM (10<sup>3</sup>/ul) | 12<br>±2.1 | 11.8<br>±2.9 | 10.3<br>±2.0 | 8.3<br>±1.8 |
| MONO (10<sup>3</sup>/ul) | 0.7<br>±0.2 | 0.6<br>±0.1 | 0.5<br>±0.1 | 0.4<br>±0.1 |
| GRAN (10<sup>3</sup>/ul) | 1.9<br>±0.9 | 1.7<br>±0.3 | 1.2<br>±0.4 | 1.3<br>±0.5 |
| LYM % | 83.1<br>±5.2 | 83.8<br>±1.1 | 86.7<br>±1.9 | 83.8<br>±3.1 |
| MONO % | 4.0<br>±0.8 | 3.7<br>±0.5 | 3.3<br>±0.4 | 3.6<br>±1.0 |
| GRAN % | 12.9<br>±4.4 | 12.5<br>±1.1 | 10.0<br>±1.5 | 12.7<br>±2.5 |
| HCT (%) | 54.5<br>±4.1 | 52.7<br>±1.8 | 57.9<br>±2.4 | 58.9<br>±7.9 |
| HGB (g/dl) | 16.4<br>±0.3 | 15.1<br>±0.4<br>(†P=0.00316) | 16.6<br>±0.7 | 17.1<br>±1.3 |
| RDW<sub>a</sub> (fl) | 32.9<br>±1.7 | 34.5<br>±1.1 | 36.6<br>±0.9 | 36.4<br>±3.2 |
| RDW % (%) | 17.3<br>±0.5 | 17.5<br>±0.4<br>(†P=0.00787) | 17.9<br>±0.3 | 18.4<br>±0.5<br>(*P=0.00311) |
| MCH (pg) | 14.4<br>±0.4 | 14.1<br>±0.1 | 14.7<br>±0.4 | 14.5<br>±0.2 |
| MCV (fl) | 47.9<br>±2.6 | 49.5<br>±1.5 | 51.0<br>±0.7 | 49.8<br>±3.6 |
| MCHC (g/dl) | 30.1<br>±2.1 | 28.6<br>±0.6 | 28.7<br>±0.4 | 29.4<br>±2.4 |
| RBC (10<sup>6</sup>/μl) | 11.4<br>±0.4 | 10.6<br>±0.3 | 11.3<br>±0.5 | 11.8<br>±1.0 |
| PLT (10<sup>3</sup>/μl) | 286.3<br>±<br>70.2 | 417.3<br>±<br>59.6 | 467.8<br>±<br>136.1 | 443.0<br>±<br>150.0 |
| MPV (fl) | 6.5<br>±0.4 | 6.5<br>±0.3 | 6.5<br>±0.4 | 6.2<br>±0.1 |

\*These data on 4×4 = 16 mice are a subset of the hematology data in paper #1>table 4. Each “N” represents an analysis of the blood from 1 mouse. We used these mice in the IFC studies (see Table 2 and Table 5) and as the basis for the mathematical simulations. Because we observed no statistically significant differences between sexes in paper #1>table 4, we grouped a males and females for this study. dKO,
double knockout (i.e., *Aqp1*<sup>-/-</sup>, *Rhag*<sup>-/-</sup>). We analyzed data using a one-way ANOVA, followed by the Holm-Bonferroni (Holm, 1979) correction (see Methods). For each parameter (e.g., MCV) we made all possible comparisons between from the four genotypes. \* denotes a significant difference compared with WT. † denotes a significant difference compared with dKO. *P*-values are displayed in parentheses below the mean value if the comparison is significant. See Supporting file 3 for *P*-values for all comparisons.

##### Effect of drugs

IFC shows that under control (no-drug) conditions, nBCDs account for only ∼1.4% of cells among WTs, and ∼2.5% in dKOs (Table 5). These percentages are consistent with our observation of spherical cells or cells with spicules by DIC microscopy/videography are rare (see <u>above</u>^36^ and Figure 4). IFC brightfield images (Figure 8A) reveal that the nBCDs are a mixture of poikilocyte types, mainly echinocytes, acanthocytes and spherocytes.

**Figure 8.**
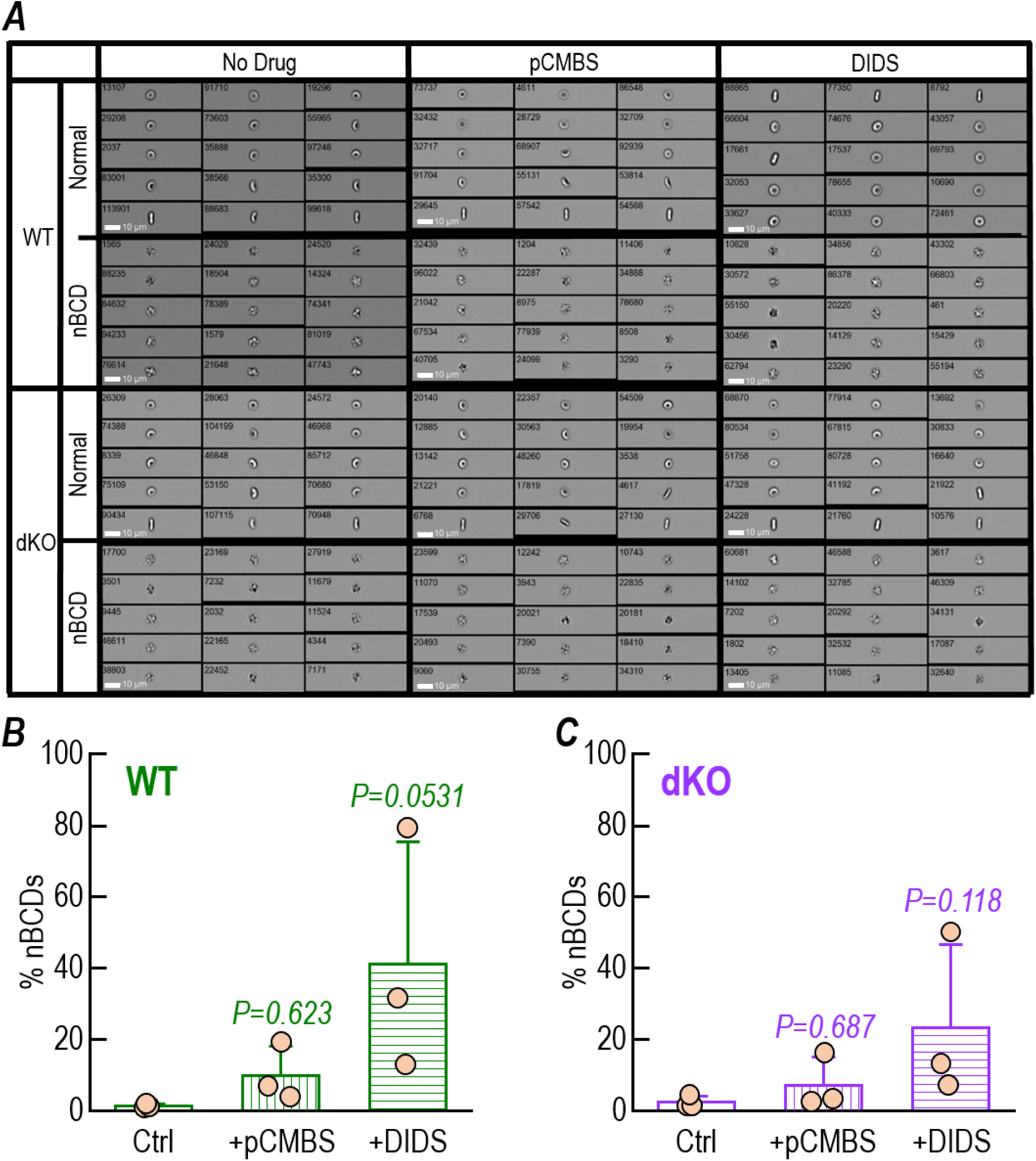
Analysis of bright-field images of WT or dKO RBCs from ImageStream flow cytometry quantify nBCD prevalence following incubation with no drug, pCMBS or DIDS. *A*, We collected blood samples and pre-incubate with no drug (Ctrl), 1 mM pCMBS for 15 min, or 200 μM DIDS for 1 h as described in <u>Methods</u>^53^. We applied gating schemes as described in Figure 7*A*→*E* to discriminate mature RBCs from all other particles. For Ctrl, 1 mM pCMBS and 200 μM DIDS drug-treatment conditions we show 15 examples of normal biconcave disc shaped RBCs or non-biconcave disc (nBCD) shaped from WT and dKO blood. *B,* IFC analysis of the degree of non-BCD formation in experiments on WT RBCs treated with pCMBS or DIDS. *C,* Prevalence of nBCDs among RBCs, treated with pCMBS or DIDS, from dKO mice. In both panels *B* and *C* Bars represent means ± SD and we performed one-way ANOVA, followed by the Holm-Bonferroni correction (see <u>Methods</u>^35^) to test for significant differences between Ctrl and the two drug-treated conditions on WT or dKO RBCs. *P*-values vs. Ctrl are reported above each drug-treated bar. Experiments were performed on blood samples from 3 age-matched pairs of WT vs dKO mice.

**Table 5.** Quantification of Ø_Major_, from IFC, among mature RBCs from WT and dKO mice treated with inhibitors *.

| Mouse | Drug | Cell shape | n total | % | $\emptyset_{\text{Major}}$ ($\mu\text{m}$) | SD | SEM |
| --- | --- | --- | --- | --- | --- | --- | --- |
| WT | No Drug | Biconcave Disk | 224756 | 98.59 | 6.73 | 1.07 | 0.00225 |
|  |  | nBCD | 3223 | 1.41 | 6.47 | 0.89 | 0.0157 |
|  | pCMBS | Biconcave Disk | 111534 | 91.29 | 6.67 | 0.87 | 0.00183 |
|  |  | nBCD | 10646 | 8.71 | 5.93 | 0.57 | 0.0101 |
|  | DIDS | Biconcave Disk | 102170 | 58.97 | 6.43 | 0.79 | 0.0017 |
|  |  | nBCD | 71075 | 41.03 | 5.77 | 0.70 | 0.0123 |
| dKO | No Drug | Biconcave Disk | 183684 | 97.49 | 6.22 | 0.82 | 0.00174 |
|  |  | nBCD | 4726 | 2.51 | 6.11 | 0.57 | 0.00996 |
|  | pCMBS | Biconcave Disk | 151278 | 94.28 | 6.59 | 0.92 | 0.00193 |
|  |  | nBCD | 9184 | 5.72 | 5.90 | 0.59 | 0.0103 |
|  | DIDS | Biconcave Disk | 141579 | 78.73 | 6.47 | 0.87 | 0.00184 |
|  |  | nBCD | 38259 | 21.27 | 5.85 | 0.59 | 0.0104 |

\* We split blood samples from three age-matched pairs of WT and dKO mice into three equal parts and incubated each of these with a different inhibitor (see Methods). We sorted RBCs according to their fluorescence signature by IFC and analyzed the number and $\emptyset_{\text{Major}}$ of normal RBCs and nBCD following each drug treatment (see Figure 8). dKO, double knockout (i.e., *Aqp1*<sup>-/-</sup>, *Rhag*<sup>-/-</sup>).

**Table 6.**
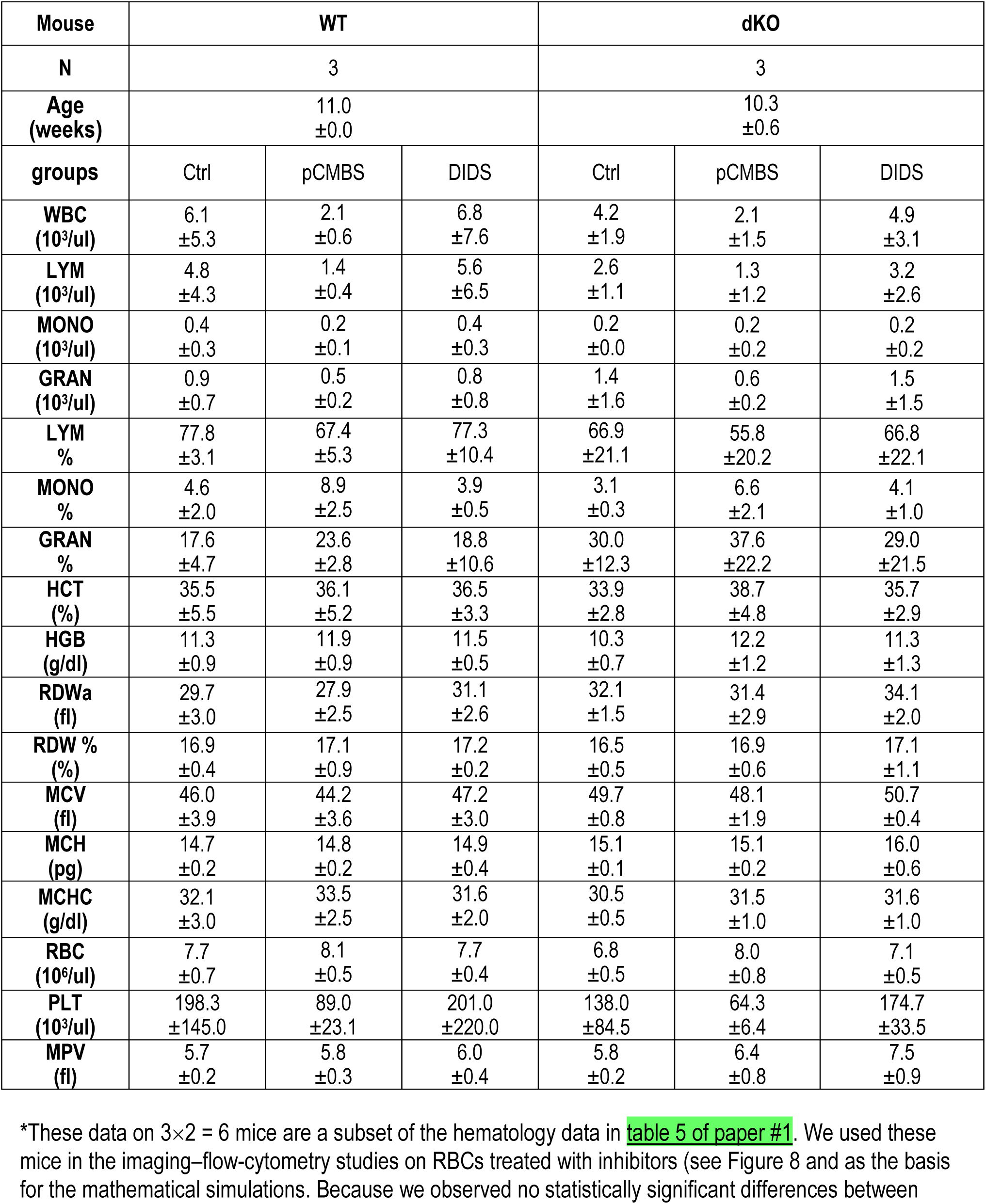
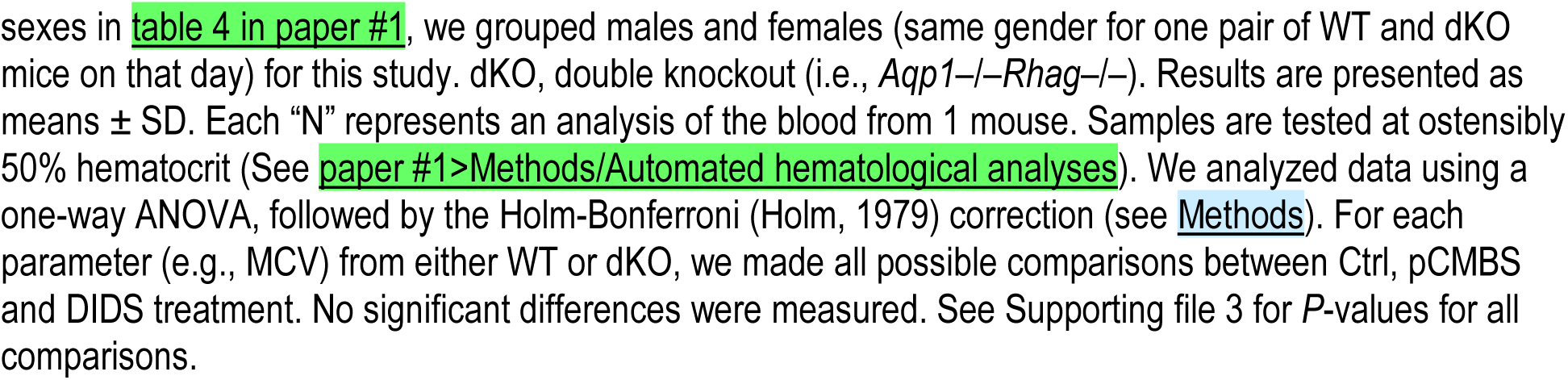
Summary of automated hematological data obtained from the same samples used for IFC and for mathematical simulations*.

| Mouse | WT |  |  | dKO |  |  |
| --- | --- | --- | --- | --- | --- | --- |
| N | 3 |  |  | 3 |  |  |
| Age (weeks) | 11.0<br>±0.0 |  |  | 10.3<br>±0.6 |  |  |
| groups | Ctrl | pCMBS | DIDS | Ctrl | pCMBS | DIDS |
| WBC<br>(10<sup>3</sup>/ul) | 6.1<br>±5.3 | 2.1<br>±0.6 | 6.8<br>±7.6 | 4.2<br>±1.9 | 2.1<br>±1.5 | 4.9<br>±3.1 |
| LYM<br>(10<sup>3</sup>/ul) | 4.8<br>±4.3 | 1.4<br>±0.4 | 5.6<br>±6.5 | 2.6<br>±1.1 | 1.3<br>±1.2 | 3.2<br>±2.6 |
| MONO<br>(10<sup>3</sup>/ul) | 0.4<br>±0.3 | 0.2<br>±0.1 | 0.4<br>±0.3 | 0.2<br>±0.0 | 0.2<br>±0.2 | 0.2<br>±0.2 |
| GRAN<br>(10<sup>3</sup>/ul) | 0.9<br>±0.7 | 0.5<br>±0.2 | 0.8<br>±0.8 | 1.4<br>±1.6 | 0.6<br>±0.2 | 1.5<br>±1.5 |
| LYM<br>% | 77.8<br>±3.1 | 67.4<br>±5.3 | 77.3<br>±10.4 | 66.9<br>±21.1 | 55.8<br>±20.2 | 66.8<br>±22.1 |
| MONO<br>% | 4.6<br>±2.0 | 8.9<br>±2.5 | 3.9<br>±0.5 | 3.1<br>±0.3 | 6.6<br>±2.1 | 4.1<br>±1.0 |
| GRAN<br>% | 17.6<br>±4.7 | 23.6<br>±2.8 | 18.8<br>±10.6 | 30.0<br>±12.3 | 37.6<br>±22.2 | 29.0<br>±21.5 |
| HCT<br>(%) | 35.5<br>±5.5 | 36.1<br>±5.2 | 36.5<br>±3.3 | 33.9<br>±2.8 | 38.7<br>±4.8 | 35.7<br>±2.9 |
| HGB<br>(g/dl) | 11.3<br>±0.9 | 11.9<br>±0.9 | 11.5<br>±0.5 | 10.3<br>±0.7 | 12.2<br>±1.2 | 11.3<br>±1.3 |
| RDW<sub>a</sub><br>(fl) | 29.7<br>±3.0 | 27.9<br>±2.5 | 31.1<br>±2.6 | 32.1<br>±1.5 | 31.4<br>±2.9 | 34.1<br>±2.0 |
| RDW %<br>(%) | 16.9<br>±0.4 | 17.1<br>±0.9 | 17.2<br>±0.2 | 16.5<br>±0.5 | 16.9<br>±0.6 | 17.1<br>±1.1 |
| MCV<br>(fl) | 46.0<br>±3.9 | 44.2<br>±3.6 | 47.2<br>±3.0 | 49.7<br>±0.8 | 48.1<br>±1.9 | 50.7<br>±0.4 |
| MCH<br>(pg) | 14.7<br>±0.2 | 14.8<br>±0.2 | 14.9<br>±0.4 | 15.1<br>±0.1 | 15.1<br>±0.2 | 16.0<br>±0.6 |
| MCHC<br>(g/dl) | 32.1<br>±3.0 | 33.5<br>±2.5 | 31.6<br>±2.0 | 30.5<br>±0.5 | 31.5<br>±1.0 | 31.6<br>±1.0 |
| RBC<br>(10<sup>6</sup>/ul) | 7.7<br>±0.7 | 8.1<br>±0.5 | 7.7<br>±0.4 | 6.8<br>±0.5 | 8.0<br>±0.8 | 7.1<br>±0.5 |
| PLT<br>(10<sup>3</sup>/ul) | 198.3<br>±145.0 | 89.0<br>±23.1 | 201.0<br>±220.0 | 138.0<br>±84.5 | 64.3<br>±6.4 | 174.7<br>±33.5 |
| MPV<br>(fl) | 5.7<br>±0.2 | 5.8<br>±0.3 | 6.0<br>±0.4 | 5.8<br>±0.2 | 6.4<br>±0.8 | 7.5<br>±0.9 |

\*These data on 3×2 = 6 mice are a subset of the hematology data in table 5 of paper #1. We used these mice in the imaging–flow-cytometry studies on RBCs treated with inhibitors (see Figure 8 and as the basis for the mathematical simulations. Because we observed no statistically significant differences between
sexes in table 4 in paper #1, we grouped males and females (same gender for one pair of WT and dKO mice on that day) for this study. dKO, double knockout (i.e., *Aqp1*<sup>-/-</sup>*Rhag*<sup>-/-</sup>). Results are presented as means ± SD. Each “N” represents an analysis of the blood from 1 mouse. Samples are tested at ostensibly 50% hematocrit (See paper #1>Methods/Automated hematological analyses). We analyzed data using a one-way ANOVA, followed by the Holm-Bonferroni (Holm, 1979) correction (see Methods). For each parameter (e.g., MCV) from either WT or dKO, we made all possible comparisons between Ctrl, pCMBS and DIDS treatment. No significant differences were measured. See Supporting file 3 for *P*-values for all comparisons.

In cells from WT mice, nBCDs rise from ∼1.4% of the total count under control conditions to ∼8.7% after 15 min pretreatment with 1 mM pCMBS (Table 5), and to ∼41% after 1 h pretreatment with 200 μM DIDS (see Figure 8B and Table 5).

We already noted—in the context of DIC experiments—that drug-treated cells entered the SF device within 5 min of the completion of the pretreatment, whereas the delay for DIC experiments was typically 30-90 min. Here for IFC experiments, the delay was also typically 60-90 min. Thus, the estimates of nBCDs prevalence likely overestimate what we would have observed in the SF device.

If we examine cells from dKO mice, the prevalences of nBCDs in drug-treated preparations are far lower than for WT mice: only ∼5.7% with pCMBS pretreatment (∼35% lower vs. WT), and only ∼21% with DIDS pretreatment (∼48% lower vs. WT; see Figure 8C vs. *B* and Table 5).

In our mathematical modeling (see <u>paper #3</u>), we used the mean Ø_Major_ values and the percentages of BCDs vs. nBCDs (overestimates though they may be) in each condition to take into account the contribution that the inhibitor-induced changes in RBC morphology have on *k*_HbO_2__.

### Proteomics

Because the genetic deletion of one protein could lead to alterations in the expression of others, we assessed effects of the KOs on RBC membrane-protein levels by generating RBC ghosts, isolating the proteins, and quantitating them by mass spectrometry (label-free LC/MS/MS).

We determined the apparent abundance of 7,188 unique peptides from 1105 total proteins, not all of RBC origin, including 212 plasma-membrane–associated proteins (PMA proteins; Supporting file 1).

#### Validating the deletion of targeted proteins in knockout strains

Our three strains with genetic deletions—namely, *Aqp1–/–*, *Rhag–*/–, and *Aqp1–/–Rhag–*/– (i.e., the dKO)—produced the expected elimination of the cognate proteins from RBC membranes. Thus, AQP1 is absent from the AQP1-KOs and dKOs (Figure 9A), whereas RhAG is absent from the RhAG-KOs and dKOs (Figure 9B). In addition, the deletions of *Rhag* in both the RhAG-KOs and dKOs also eliminate RhD (Figure 9C), which forms heterotrimers with RhAG (Gruswitz *et al*., 2010) and falls pari passu with RhAG. We describe the vastly smaller effects of the knockouts on the inferred abundance of other individual proteins <u>below</u>^37^.

**Figure 9.**
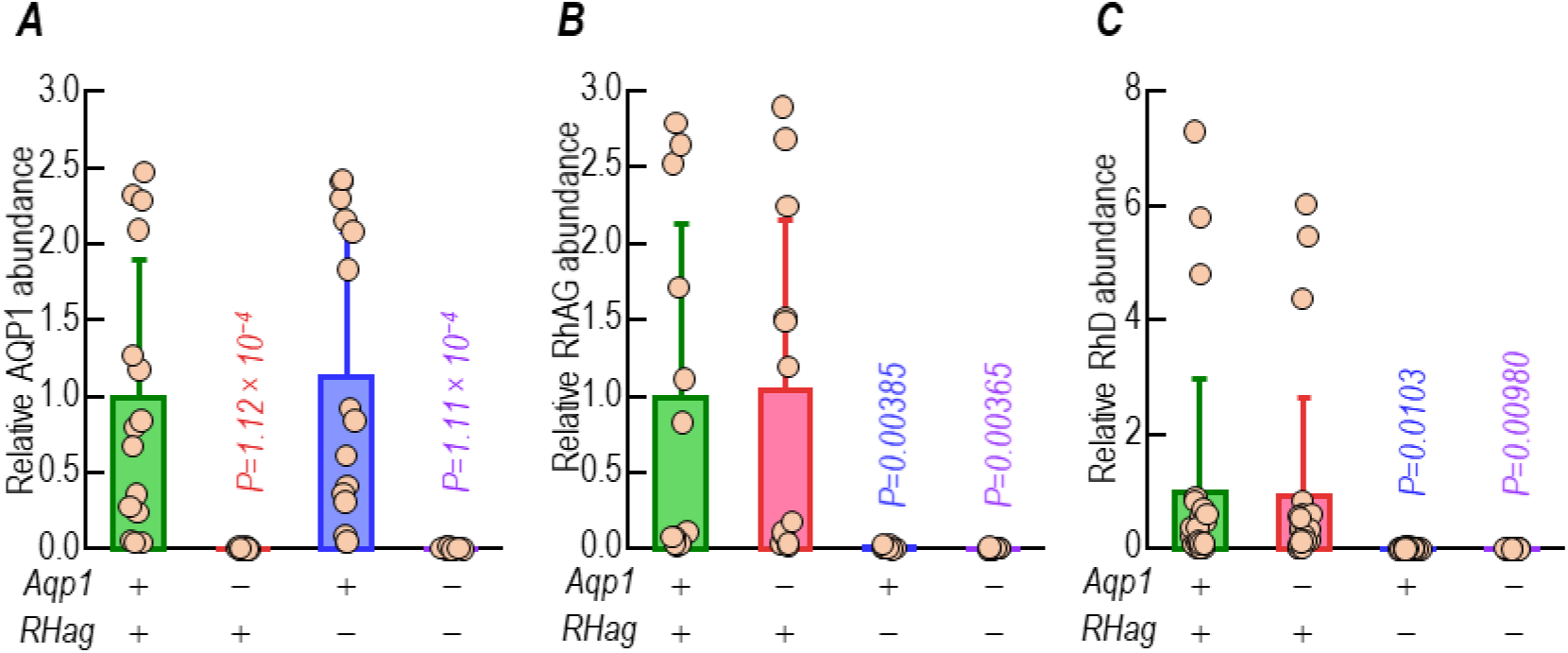
Effect of genetic deletions of *Aqp1*, *Rhag*, or both on AQP1, RhAG and RhD protein expression. Proteomic analysis by LC/MS/MS of all peptides of *A*, AQP1, *B*, RhAG, and *C*, RhD from WT (green bars), AQP1-KO (pink bars), RhAG-KO (blue bars) or dKO (purple bars) RBC ghosts, normalized to 1 for WT. The x-axis annotation reports whether the *Aqp1* or *Rhag* genes are present (+) or not (−). Bars represent normalized mean peak intensity (AUC) ± SD. Each individual data point (beige circles) reports the normalized abundance of one unique peptide detected from one ghost sample purified from one mouse (e.g., in panel *A*, for AQP1 protein, we plot 15 individual data points overlaying each bar, representing 5 unique peptides, from 3 RBC ghost samples from each of four strains). We performed one-way ANOVA with the Holm-Bonferroni correction to test for statistical significance, as described in the <u>Methods</u>^35^, *P*-values vs. WT for the effect of genetic deletions of *Aqp1*, *Rhag*, or both on AQP1, RhAG and RhD protein expression are displayed above bars for the knocked-out gene(s) in each panel. See Supporting file 4 for *P*-values for all comparisons.

#### PMA proteins of highest inferred abundance in WT ghosts

Among the 50 total proteins with the greatest inferred abundance, 22 are PMA proteins (Table 7). Figure 10 displays the abundance of the unique peptides from the top 22 PMA proteins in WT RBC ghosts. These 22 proteins span a ∼40-fold range in inferred abundance. As can be seen by totaling the numbers Table 7, the 22 top PMA proteins comprise ∼87% of PMA protein abundance in WT ghosts (column 5: from AE1 down to SLC16A10), and ∼42% of total protein abundance (column 4)^38^. The 25 PMA proteins with the next-highest inferred abundance—also listed in Table 7 (start at KEL down to DNAJC5/Hsp40)—collectively comprise merely an additional ∼5.7% of PMA protein abundance in WT ghosts, and ∼2.8% of total protein abundance. Thus, a relatively small number of PMA proteins represent an overwhelming fraction of the summated abundance of proteins associated with the RBC membrane of WT mice.

**Figure 10.**
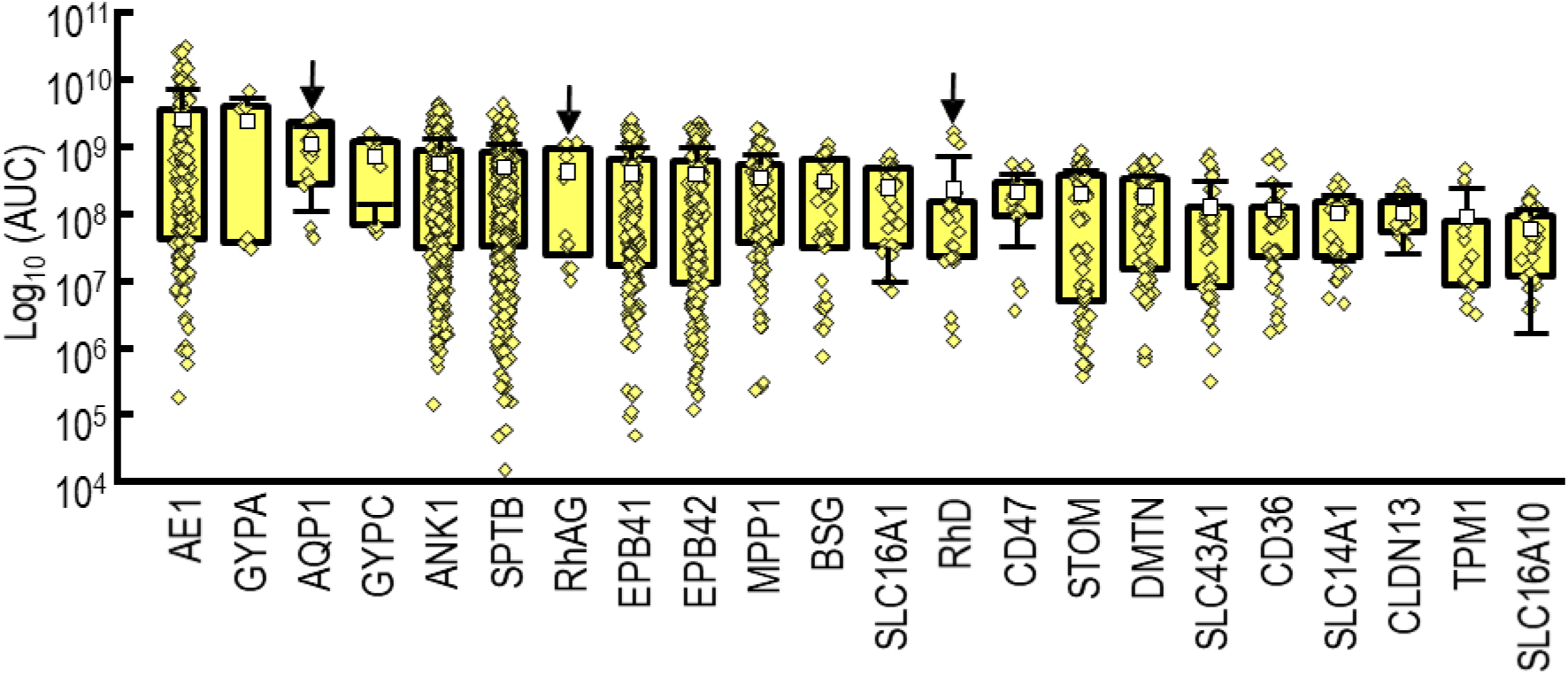
Proteomic analysis by LC/MS/MS of proteins in samples of RBC ghosts from WT mice. *A,* Plasma-membrane–associated proteins, ranked by abundance, as inferred from mass-spectrometry. AUC is area under curve. We detected a total of <u>7,188</u> unique peptides from <u>1,104</u> unique proteins, not all of RBC origin (Supporting file 1). Of these, <u>1,902</u> peptides and <u>212</u> proteins are “plasma-membrane– associated” (integral-membrane proteins + others) as defined by the data-analysis software Peaks (see <u>Methods</u>^35^). Each individual data point (yellow diamonds) overlaying a box plot reports the abundance of one unique peptide detected from one ghost sample purified from one mouse (e.g., for AE1 we plot 171 individual data points overlaying each bar, representing 57 unique peptides, from 3 RBC ghost samples from the WT mice). None of the 47 PMA proteins of greatest inferred abundance exhibited a significant change in response to any of the deletions. Of these 47, panel *A* shows the 22 PMA proteins, with greatest inferred abundance (Table 7 provides protein glossary and rank order of abundance of all 47 PMA proteins). Arrows AQP1, RhAG, and RhD which are the targets of the genetic KO mice. Of these 22 most abundant PMA protein species, only the intended target proteins in each KO strain display a significant change in abundance. See Figure 10*A* and *B* for effects of gene deletions on abundance of AQP1, RhAG, and RhD. See Figure 11, for effects of gene deletions on abundance of each of these 22 proteins. Others have shown that RhD expression requires *Rhag* in mice (Goossens *et al*., 2010). Boxes represent the interquartile range. Open white squares represent the mean abundance. Whiskers represent one standard deviation. We performed one-way ANOVA with the Holm-Bonferroni correction to test for statistical significance, as described in <u>Methods</u>^35^. See Supporting file 4 for *P*-values for all comparisons.

**Table 7:**
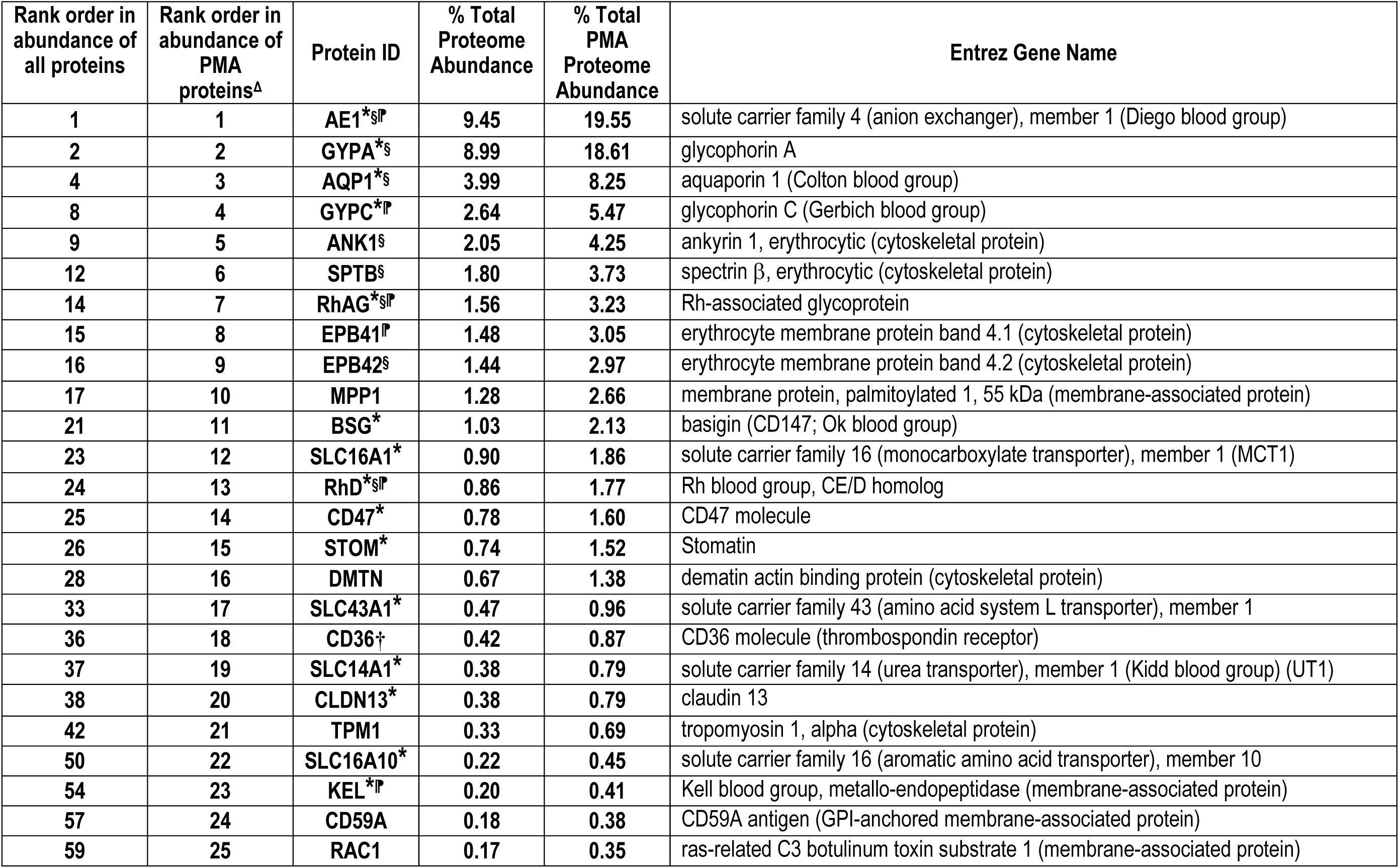
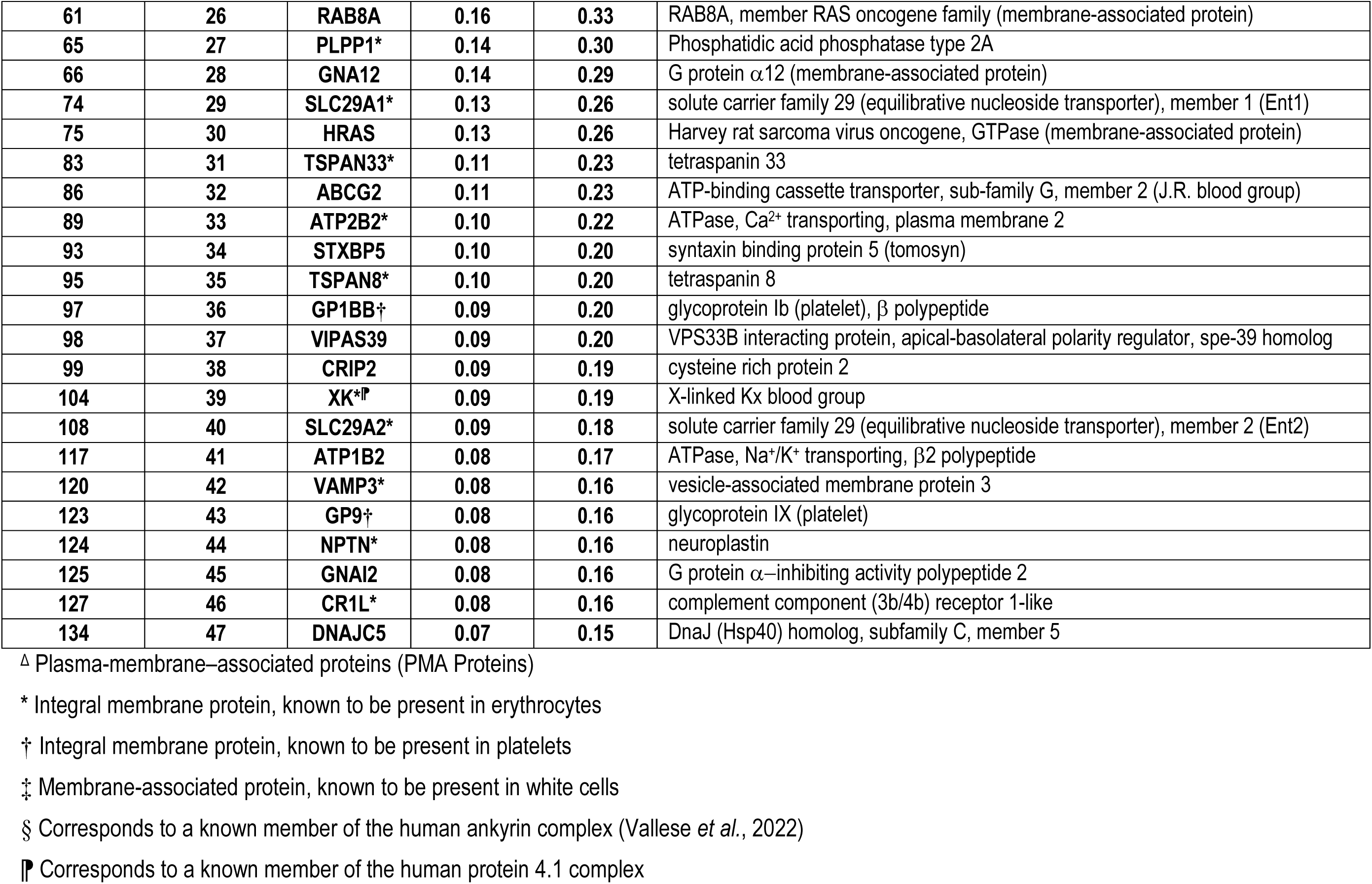
The 47 “Plasma-membrane–associated” (PMA) proteins with the largest inferred abundance in samples of RBC ghosts from WT mice.

The most dominant protein is AE1 (represented by 57 unique peptides in each of three analyzed WT mice, for a total of 3 × 57 = 171 peptides), which, by itself, comprises ∼9.5% of the total WT RBC ghost proteome in our dataset, and ∼20% of the identified PMA proteome (Table 7).

The second most abundant PMA protein is glycophorin A (GYPA), which facilitates AE1 cell-surface expression (Groves & Tanner, 1992, 1994; Pang & Reithmeier, 2009; Hsu *et al*., 2022) and comprises ∼9% of the total WT RBC ghost proteome, and ∼19% of all PMA protein content (Table 7).

Collectively AQP1—the only AQP species detected in the murine ghosts in these analyses—and the Rh_Cx_, the proteins targeted for knockout in the present paper (indirectly in the case of RhD), comprise ∼6.4% of the total WT RBC ghost proteome, and ∼13% of the WT RBC ghost PMA proteins. All of them are among the top 13 most abundant PMA proteins (see Figure 10) and top 24 in overall abundance (see Supporting file 1).

#### Inferred abundance of SLC and other transport proteins in WT ghosts

The data in paper #1 indicate that, although AQP1 and Rh_Cx_ account for ∼55% of *P*_M,O_2__, other membrane protein(s) sensitive to pCMBS account for an additional 35%. In principle, any combination of other membrane proteins—especially multitopic proteins that form transporters and channels—could account for this missing 35%.

##### SLC proteins

Besides AE1, we detected 23 other SLC proteins (Table 8). These represent only ∼3% of the total RBC proteome, and ∼5% of the PMA protein content. The most abundant of these 23 other SLC proteins is the monocarboxylate transporter 1 (MCT1; SLC16A1), representing ∼2% of PMA proteins. In the 4^th^ row of Table 8, ranked #19 in PMA protein abundance, is the UT-B urea transporter (SLC14A1), representing only ∼0.38%; it is also demonstrated to act as an NH_3_ channel (Geyer *et al*., 2013*a*). In row 9 of Table 8, ranked #65 in PMA protein abundance with just ∼0.12% of the PMA proteome is NBCe1 (SLC4A1), that in addition to performing Na^+^/CO_3_^=^ cotransport (Lee *et al*., 2023), is hypothesized to mediate transmembrane CO_2_ fluxes via non-canonical pathways within the transporter (Moss *et al*., 2019).

**Table 8:** SLC proteins in WT RBC ghosts, ranked by abundance among plasma-membrane–associated proteins.

| Rank order Proteins |  | Primary accession | Protein ID | Alt. Protein ID | RBC Proteome |  | Full Protein name | Mean Intensity (AUC) | St. Dev | n |
| --- | --- | --- | --- | --- | --- | --- | --- | --- | --- | --- |
| All | PMA<sup>Δ</sup> |  |  |  | % Total | %PMA |  |  |  |  |
| 1 | 1 | P04919 | SLC4A1 | AE1 | 9.45 | 19.6 | Anion exchange protein 1 | 2.61×10<sup>9</sup> | 4.91×10<sup>9</sup> | 171 |
| 14 | 7 | Q9QUT0 | SLC42A1 | RhAG | 1.56 | 3.23 | Rh-associated glycoprotein | 5.00×10<sup>8</sup> | 4.88×10<sup>8</sup> | 12 |
| 23 | 12 | P53986 | SLC16A1 | MCT1 | 0.898 | 1.86 | Monocarboxylate transporter 1 | 2.48×10<sup>8</sup> | 2.38×10<sup>8</sup> | 27 |
| 24 | 13 | Q9QUT0 | SLC42A5 | RhD | 0.857 | 1.77 | Rh blood group, CE/D homolog | 2.75×10<sup>8</sup> | 4.67×10<sup>8</sup> | 24 |
| 33 | 17 | Q8BSM7 | SLC43A1 | LAT3 | 0.466 | 0.964 | Large neutral amino acids transporter small subunit 3 | 1.29×10<sup>8</sup> | 1.90×10<sup>8</sup> | 42 |
| 37 | 19 | Q8VHL0 | SLC14A1 | UT-B | 0.384 | 0.795 | Urea transporter 1 | 1.06×10<sup>8</sup> | 8.62×10<sup>7</sup> | 36 |
| 50 | 22 | Q3U9N9 | SLC16A10 | MCT10 | 0.218 | 0.451 | Monocarboxylate transporter 10 | 6.02×10<sup>7</sup> | 5.86×10<sup>7</sup> | 30 |
| 74 | 29 | Q9JIM1 | SLC29A1 | ENT1 | 0.127 | 0.263 | Equilibrative nucleoside transporter 1 | 3.50×10<sup>7</sup> | 4.26×10<sup>7</sup> | 33 |
| 108 | 40 | Q61672 | SLC29A2 | ENT2 | 0.0892 | 0.185 | Equilibrative nucleoside transporter 2 | 2.46×10<sup>7</sup> | 4.57×10<sup>5</sup> | 3 |
| 142 | 51 | Q9JHI9 | SLC40A1 | MTP1 | 0.0707 | 0.146 | Ferroportin | 1.95×10<sup>7</sup> | 1.10×10<sup>7</sup> | 24 |
| 180 | 65 | O88343 | SLC4A4 | NBCe1 | 0.0573 | 0.119 | Electrogenic Na<sup>+</sup> HCO<sub>3</sub><sup>-</sup> cotransporter 1 | 1.58×10<sup>7</sup> | 3.39×10<sup>6</sup> | 3 |
| 215 | 74 | P32037 | SLC2A3 | GLUT3 | 0.0432 | 0.0893 | Facilitated glucose transporter member 3 | 1.19×10<sup>7</sup> | 1.09×10<sup>7</sup> | 21 |
| 349 | 93 | Q61165 | SLC9A1 | NHE1 | 0.0268 | 0.0555 | Na<sup>+</sup>/H<sup>+</sup> exchanger 1 | 7.41×10<sup>6</sup> | 4.18×10<sup>6</sup> | 24 |
| 350 | 94 | P58735 | SLC26A1 | SAT1 | 0.0268 | 0.0554 | Sulfate anion transporter 1 | 7.40×10<sup>6</sup> | 3.41×10<sup>6</sup> | 6 |
| 490 | 123 | P10852 | SLC3A2* | MDU1 | 0.0171 | 0.0355 | Amino acid transporter (CD98 Antigen) | 4.73×10<sup>6</sup> | 2.28×10<sup>6</sup> | 30 |
| 533 | 132 | P49282 | SLC11A2 | DMT1 | 0.0154 | 0.0318 | Natural resistance-associated macrophage protein 2 | 4.24×10<sup>6</sup> | 2.99×10<sup>6</sup> | 12 |
| 555 | 134 | Q09143 | SLC7A1 | CAT1 | 0.0146 | 0.0302 | High affinity cationic amino acid transporter 1 | 4.03×10<sup>6</sup> | 4.92×10<sup>5</sup> | 3 |
| 594 | 140 | Q9Z127 | SLC7A5 | LAT1 | 0.0133 | 0.0276 | Large neutral amino acids transporter small subunit | 3.68×10<sup>6</sup> | 2.76×10<sup>6</sup> | 12 |
| 669 | 151 | Q60857 | SLC6A4 | SERT | 0.0116 | 0.0240 | Na<sup>+</sup>-dependent serotonin transporter | 3.20×10<sup>6</sup> | 2.98×10<sup>6</sup> | 3 |
| 729 | 161 | P28571 | SLC6A9 | GLYT1 | 0.0101 | 0.0208 | Na<sup>+</sup> Cl<sup>-</sup>-dependent glycine transporter 1 | 2.78×10<sup>6</sup> | 1.91×10<sup>6</sup> | 6 |
| 737 | 164 | Q6X893 | SLC44A1 | CD92 | 0.00987 | 0.0204 | Choline transporter-like protein 1 | 2.73×10<sup>6</sup> | 1.01×10<sup>6</sup> | 9 |
| 793 | 173 | Q60738 | SLC30A1* | ZnT1 | 0.00875 | 0.0181 | H<sup>+</sup>-coupled Zn<sup>2+</sup> antiporter | 2.42×10<sup>6</sup> | 7.40×10<sup>5</sup> | 12 |
| 850 | 181 | Q9WVC8 | SLC26A3 | DRA | 0.00741 | 0.0153 | Cl<sup>-</sup> anion exchanger | 2.05×10<sup>6</sup> | 4.61×10<sup>5</sup> | 3 |
| 917 | 191 | Q3UHH2 | SLC22A23 | OCT2 | 0.00605 | 0.0125 | Organic cation transporter 2 | 1.67×10<sup>6</sup> | 1.66×10<sup>5</sup> | 3 |
| 940 | 193 | P41438 | SLC19A1 | RFC1 | 0.00558 | 0.01154 | Reduced folate transporter | 1.54×10<sup>6</sup> | 2.00×10<sup>5</sup> | 3 |
| 1092 | 210 | Q8BRU6 | SLC18A2 | VAT2 | 0.00149 | 0.00307 | Synaptic vesicular amine transporter | 4.10×10<sup>5</sup> | 2.15×10<sup>5</sup> | 3 |

##### ATP-driven pumps

Combined, 0.28% of the PMA protein of WT RBC ghosts comes from the ATP1A1 (0.096%; Supporting file 1), ATP1B2 (0.167%; Table 7) and ATP1B3 (0.013%; Supporting file 1) Na^+^/K^+^ ATPase subunits. The plasma membrane Ca^2+^-transporting ATPase 2 (ATPB2) constitutes ∼0.2% of the PMA protein (Table 7). The phospholipid-transporting ATPase 11C (ATP11C)—the catalytic component of a P4-ATPase flippase complex in the RBC plasma membrane that is critical for retaining phosphatidylserines (PS) within the inner leaflet and preventing surface exposure (Yabas *et al*., 2014)—constitutes ∼0.04% of the PMA protein (Supporting file 1).

##### Ion channels

We detect only two other PMA proteins with permanent pores, both of which are ion channels low in abundance: (1) The piezo-type mechanosensitive ion channel component 1 (PIEZO1; 0.074% of PMA proteins, Supporting file 1) defines the Er-blood group (Coste *et al*., 2010; Fang *et al*., 2021; Karamatic Crew *et al*., 2023). PIEZO1 ranks only #305 in abundance among all WT RBC proteins, and #79 among PMA proteins (2) The transient receptor potential cation channel subfamily V member 2 (TRPV2; 0.060%), which medicates Ca^2+^ influx into RBCs (Belkacemi *et al*., 2021), is #329 in overall abundance, and #90 among PMA proteins (Supporting file 1).

#### Effects of knockouts on PMA proteins of greatest inferred abundance

Among the 22 PMA proteins with greatest inferred abundance in WT ghosts (see Figure 10 and list in Table 7), none undergo significant changes in response to the deletions of *Aqp1* and/or *Rhag*. Supporting this conclusion is Figure 11*A–V*, which summarizes the fold changes (normalized to WT) for each of these top-22 PMA proteins, and for each of the three KO genotypes. Similarly, none of the next 25 most abundant PMA proteins (see Table 7 for listing) exhibit a significant change in inferred abundance in response to single or double gene deletions (see Supporting file 4).

**Figure 11.**
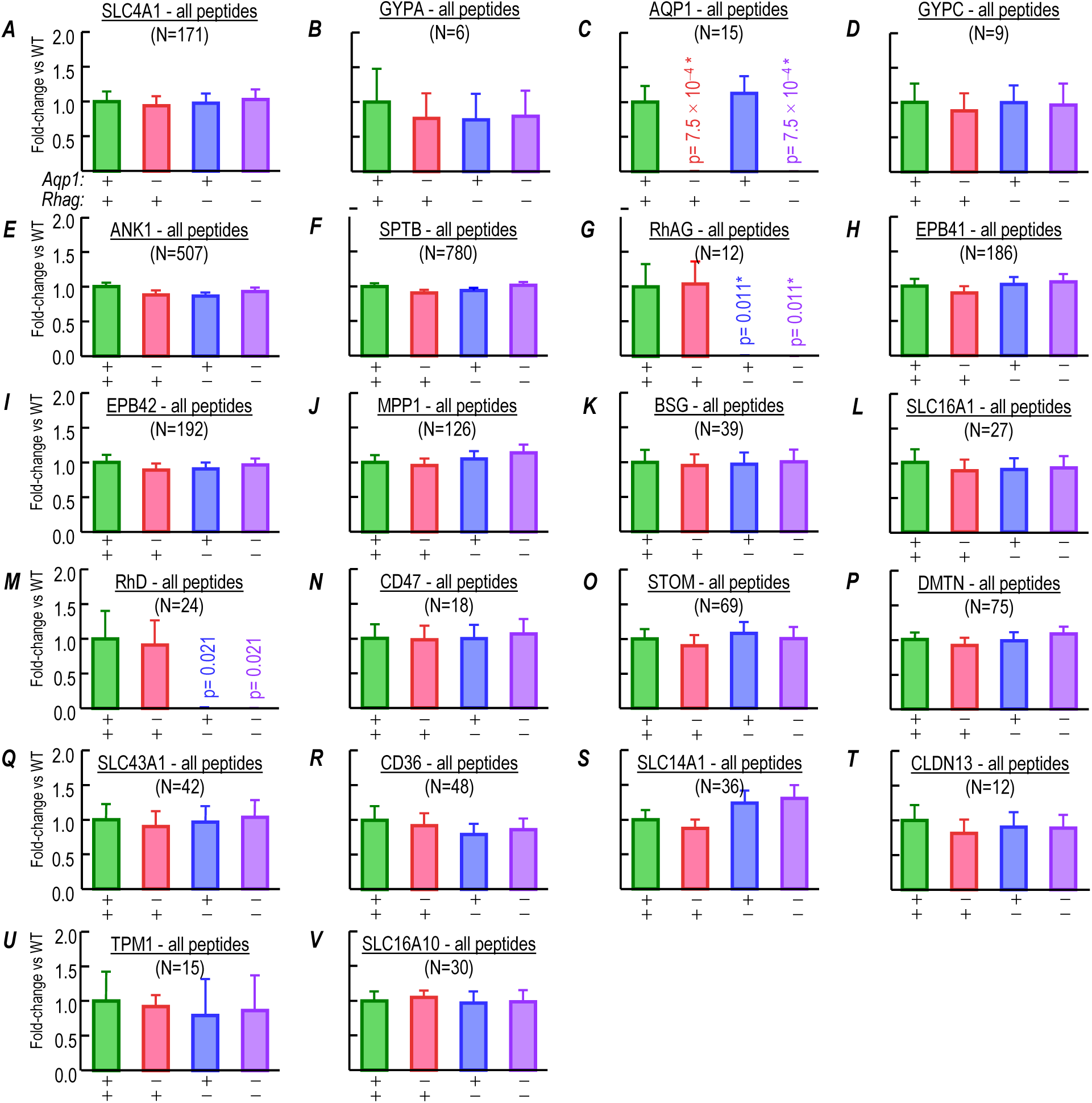
Summary of fold-changes in expression of 22 “plasma membrane– associated” proteins comparing samples from RBC ghosts of WT vs. AQP1-KO, RhAG-KO and double-KO (dKO) mice. These 22 panels represent the 22 plasma-membrane–associated proteins, from among the 50 proteins with the greatest inferred abundance in RBC ghosts from WT mice. The panels are in rank order, based on the inferred abundance of the protein in cells from WT mice (see Figure 10*A*). In each panel, data from WT mice (green bar) is the reference value for fold-changes in expression for the each mouse knockout strains (AQP1*-*KO, red bars; RhAG*-*KO, blue bars; dKO, purple bars), as determined by a mass-spectrometry based investigation of peptide peak intensity (AUC) obtained from RBC ghosts. We purify proteins from RBC ghosts of 3 mice/genotype (same samples as Figure 13) and the numbers in parentheses above the plot in each panel are the total number of peptides analyzed for each protein from blood samples acquired the 3 mice per genotype (e.g. for AE1, peptide peak intensities are measured for 57 unique peptides from 3 mice for a total of n=171 replicates). *A,* AE1 (SLC4A1). *B,* glycophorin A (GYPA). *C,* aquaporin 1 (AQP1). *D,* glycophorin C (GYPC). *E,* ankyrin 1 (ANK1). *F,* erythrocytic spectrin beta (SPTB). *G,* Rhesus blood group associated Glycoprotein A, RhAG (SLC42A1). *H* erythrocyte membrane protein band 4.1 (EPB41). *I,* erythrocyte membrane protein band 4.2 (EPB42). *J,* membrane protein, palmitoylated 1 (MPP1). *K,* basigin (BSG). *L,* monocarboxylic acid transporter member 1, MCT-1 (SLC16A1). *M,* mouse Rhesus blood group, D antigen (RhD). *N,* CD47 molecule (CD47). *O,* stomatin (STOM). *P,* dematin actin binding protein (DMTN). *Q,* solute carrier family 43 (amino acid system L transporter), member 1(SLC43A1). *R,* CD36 antigen (CD36). *S,* urea transporter member 1, UT-1 (SLC14A1). *T,* claudin 13 (CLDN13). *U,* tropomyosin 1 (TPM1). *V,* monocarboxylate transporter member 10, MCT-10 (SLC16A10). Bars represent the mean ± SD. peak intensity (AUC) for all peptides from each protein, normalized to the abundance in WT. Number of detected peptides per protein is displayed in parentheses. Not all of these proteins are necessarily RBC integral membrane proteins. We performed a one-way ANOVA, followed by the Holm-Bonferroni correction (Holm (1979); see <u>Methods</u>^35^). *P*-values vs. WT for the effect of genetic deletions of *Aqp1*, *Rhag*, or both on AQP1, RhAG and RhD protein expression are displayed above bars in panels *C* and *G*. The analyses show that expression of the target protein in RBCs of each knockout strain is essentially abolished. However, the difference in expression of other the RBC proteins in each knockout strain is not significant compared to WT. See Table 7 for glossary and rank order of abundance. See Supporting file 4 for *P*-values for all comparisons.

In fact, we find that only 27 out of 1105 proteins—16 non-PMA proteins and 11 PMA proteins—in the RBC-ghost preparation exhibit a significant change with at least 1 genetic deletion (Figure 12). Of these, the protein with the greatest inferred abundance in WT ghosts is a non-PMA protein, proteasome subunit α type-4 (PSMA4), a subunit of the 20S core proteasome complex that is present in RBCs (Kakhniashvili *et al*., 2004; Majetschak & Sorell, 2008; Neelam *et al*., 2011; Tzounakas *et al*., 2022). This protein, which ranks #101 in the WT-ghost proteome, exhibits a ∼38% lower abundance in the AQP1-KO (Figure 13); the difference is significant. The #133 ranked protein, proteasome subunit α type-1 (PSMA1), likewise exhibits a decreased abundance in AQP1-KO ghosts (Figure 13); the difference is significant. Note that the inferred abundances of PSMA1 and PSMA4 (Table 9), each, are <1% that of the #1 protein, AE1 (see Table 7).

**Figure 12.**
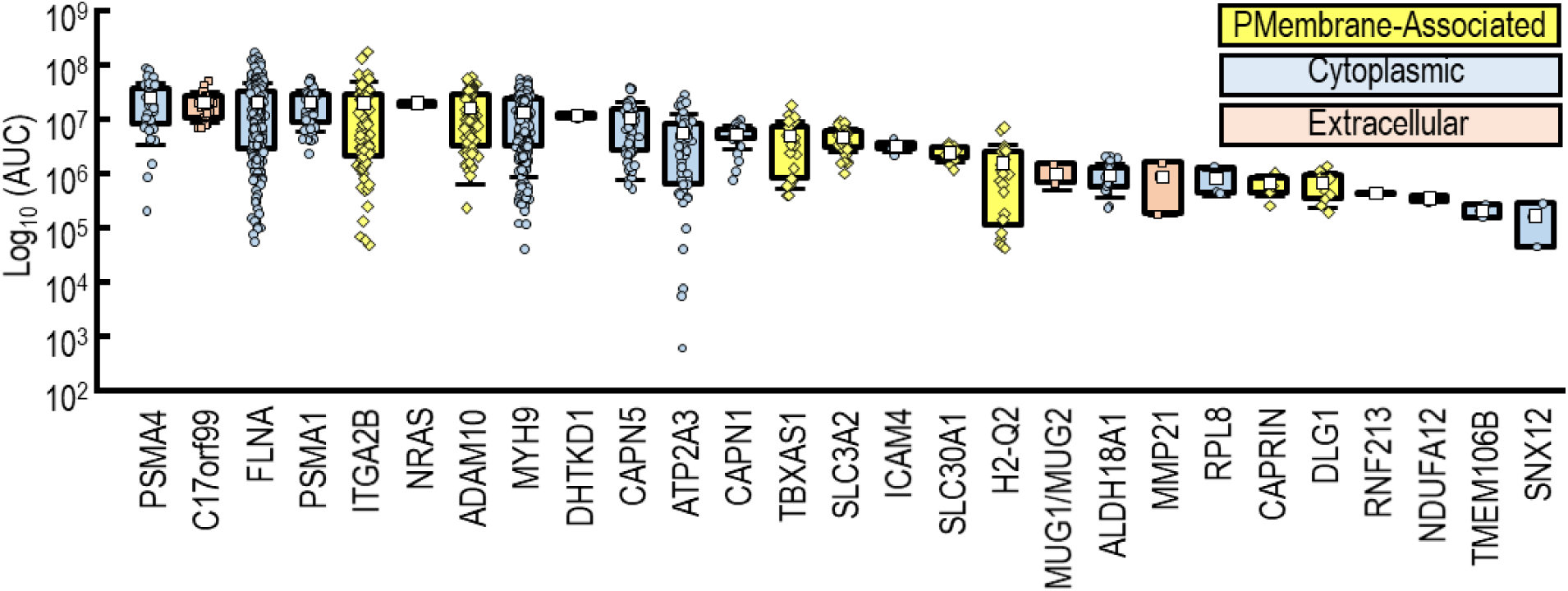
Proteins from RBC ghosts (ranked by inferred abundance in WTs) that exhibit significant differences from WT in one or more of the KO mouse strains in proteomic analysis by LC/MS/MS. Ranking of the inferred abundance of the unique peptides from the 27 proteins (out of all <u>1,104</u> detected by mass spectrometry) from RBC ghosts that demonstrated a significant difference from WT in one or more of the KO mouse strains. Each individual data point (yellow diamonds, PMA-proteins; blue circles, cytoplasmic proteins; apricot squares, extracellular proteins) overlaying a box plot reports the abundance of one unique peptide detected from one ghost sample purified from one mouse (e.g., for Integrin α2b, ITGA2B, we plot 81 individual data points overlaying each bar, representing 27 unique peptides, from 3 RBC ghost samples from the WT mice). For protein glossary and rank order by inferred abundance, see Table 9 for non PMA proteins and Table 10 for PMA. Figure 13 shows the effects of gene deletions on abundance of each of these 27 proteins. The colors of the panel backgrounds indicate the expected major location of each the proteins, as assigned by the data-analysis software Peaks (see Supporting file 1): yellow (plasma-membrane– associated), blue (cytoplasmic), and apricot (extracellular). For both panels *A* and *B*, we purified proteins from RBC ghosts of 3 mice/genotype. Each individual data point reports the abundance of one unique peptide detected from one ghost sample purified from one mouse (e.g., in Panel *A*, for RhAG protein, there are 12 individual data points, from 4 unique peptides, from 3 WT RBC ghost samples). Boxes represent the interquartile range. Open white squares represent the mean abundance. Whiskers represent one standard deviation. We performed one-way ANOVA with the Holm-Bonferroni correction to test for statistical significance, as described in <u>Methods</u>^35^.

**Figure 13.**
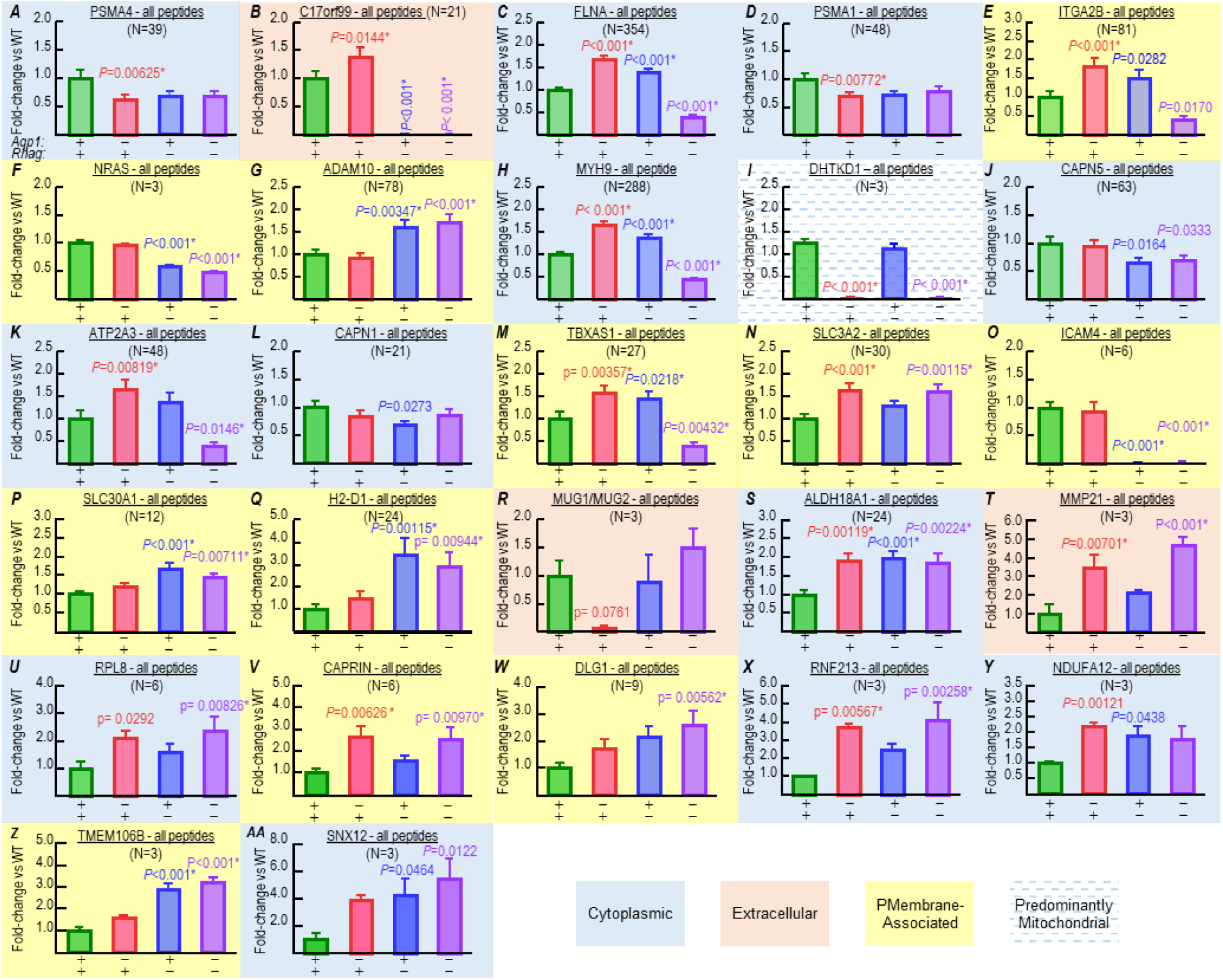
Summary of fold-changes in expression of all 27 proteins in which abundance changed in at least one KO genotype, comparing samples from RBC ghosts of WT vs. AQP1-KO, RhAG-KO, and double-KO (dKO) mice. These 27 panels represent the 27 proteins—from among all 1105 detected proteins—that exhibited a significant change in at least one knockout strain vs. WT mice. The panels are in rank order, based on the inferred abundance of the protein in cells from WT mice (see Figure 10*B*). In each panel, data from WT mice (green bar) is the reference value for fold-changes in expression for the mouse knockout strains (AQP1-KO, red bars; RhAG-KO, blue bars; dKO, purple bars), as determined by a mass-spectrometry based investigation of peptide peak intensity (AUC) obtained from RBC ghosts. *A,* proteasome (prosome, macropain) subunit, alpha type, 4 (PMSA4). *B,* Interleukin 40 (IL-40) from chromosome 17 open reading frame 99 (C17orf99). *C,* filamin A, alpha (FLNA). *D,* proteasome (prosome, macropain) subunit, alpha type, 1 (PSMA1). *E,* integrin, alpha 2b (ITGA2B). *F,* neuroblastoma RAS viral (v-ras) oncogene homolog (NRAS). *G,* ADAM metallopeptidase domain 10 (ADAM10). *H,* myosin, heavy chain 9, non-muscle (MYH9). *I,* dehydrogenase E1 and transketolase domain containing 1 (DHTKD1). *J,* calpain 5 (CAPN5). *K,* ATPase, Ca^++^-transporting, ubiquitous (ATP2A3). *L,* calpain 1 (CAPN1). *M,* thromboxane A synthase 1 (platelet) (TBXAS1). *N,* solute carrier family 3 (amino acid transporter heavy chain), member 2 (SLC3A2). *O,* intercellular adhesion molecule 4 (ICAM4). *P,* solute carrier family 30 (zinc transporter), member 1, Zn-T1 (SLC30A1). *Q,* major histocompatibility complex (H2-Q2). *R,* Murinoglobulin 1 (MUG1/MUG2). *S,* aldehyde dehydrogenase 18 family, member A1 (ALDH18A1). *T,* matrix metallopeptidase 21 (MMP21). *U,* ribosomal protein L8 (RPL8). *V,* cell cycle associated protein 1 (CAPRIN1). *W,* discs, large homolog 1 (Drosophila) (DLG1). *X,* ring finger protein 213 (RNF213). *Y,* NADH dehydrogenase (ubiquinone) 1 alpha subcomplex 12 (NDUFA12), *Z,* transmembrane protein 106B (TMEM106B). *AA,* sorting nexin 12 (SNX12). Bars represent the mean ± SD peak intensity (AUC) for all peptides from each protein normalized to the abundance in WT. Number of peptides per protein is displayed in parentheses. Not all of these proteins are from RBCs, let alone RBC integral membrane proteins. The colors of the bars indicate the location of the proteins, as assigned by the Peaks software: yellow (plasma-membrane associated), blue (cytoplasmic), and apricot (extracellular). We purified proteins from RBC ghosts of 3 mice/genotype (same samples as Figure 11). We performed a one-way ANOVA, followed by the Holm-Bonferroni correction (Holm (1979); see <u>Methods</u>^35^). *P*-values <0.05 for comparisons vs. WT are displayed above the appropriate KO bars.* denotes statistical significance. See Supporting file 6 for all *P*-values for all comparisons in each panel. For glossaries, see Table 9 for non-PMA proteins and Table 10 for PMA proteins that include the inferred abundance of each protein WT ghosts.

**Table 9:** All non-PMA with significant changes in inferred abundance in RBC ghosts in one or more KO or dKO mouse strains, ranked by abundance in WT ghosts (see Figure 13)

| Rank order in abundance of all proteins | Protein ID | % Total proteome abundance | Entrez Gene Name |
| --- | --- | --- | --- |
| 101 | PSMA4 | 0.0917 | proteasome (prosome, macropain) subunit, alpha type, 4 |
| 130 | C17ORF99 | 0.0755 | chromosome 17 open reading frame 99 (Protein IL-40) |
| 132 | FLNA | 0.0753 | filamin A, alpha (cytoskeletal protein) |
| 133 | PSMA1 | 0.0745 | proteasome (prosome, macropain) subunit, alpha type, 1 |
| 195 | MYH9 | 0.0501 | myosin, heavy chain 9, non-muscle (cytoskeletal protein) |
| 214 | DHTKD1* | 0.0433 | dehydrogenase E1 and transketolase domain containing 1 (mitochondrion) |
| 236 | CAPN5 | 0.0395 | calpain 5 (Protease) |
| 433 | ATP2A3† | 0.0203 | ATPase, Ca<sup>++</sup>-transporting, ubiquitous (a SERCA) |
| 448 | CAPN1 | 0.0194 | calpain 1 (protease) |
| 1045 | MUG1 | 0.00353 | murinoglobulin 1 (proteinase) |
| 1051 | ALDH18A1 | 0.00337 | aldehyde dehydrogenase 18 family, member A1 |
| 1052 | MMP21‡ | 0.00322 | matrix metalloproteinase 21 |
| 1059 | RPL8♥ | 0.00291 | ribosomal protein L18 |
| 1087 | RNF213 | 0.00158 | ring finger protein 213 |
| 1095 | NDUFA12 | 0.00128 | NADH dehydrogenase (ubiquinone) 1 alpha subcomplex, 12 |
| 1102 | SNX12 | 0.000604 | sorting nexin 12 |

\*Mitochondrial
†Endoplasmic reticulum
‡Extracellular
♥Ribosomal

**Table 10:**
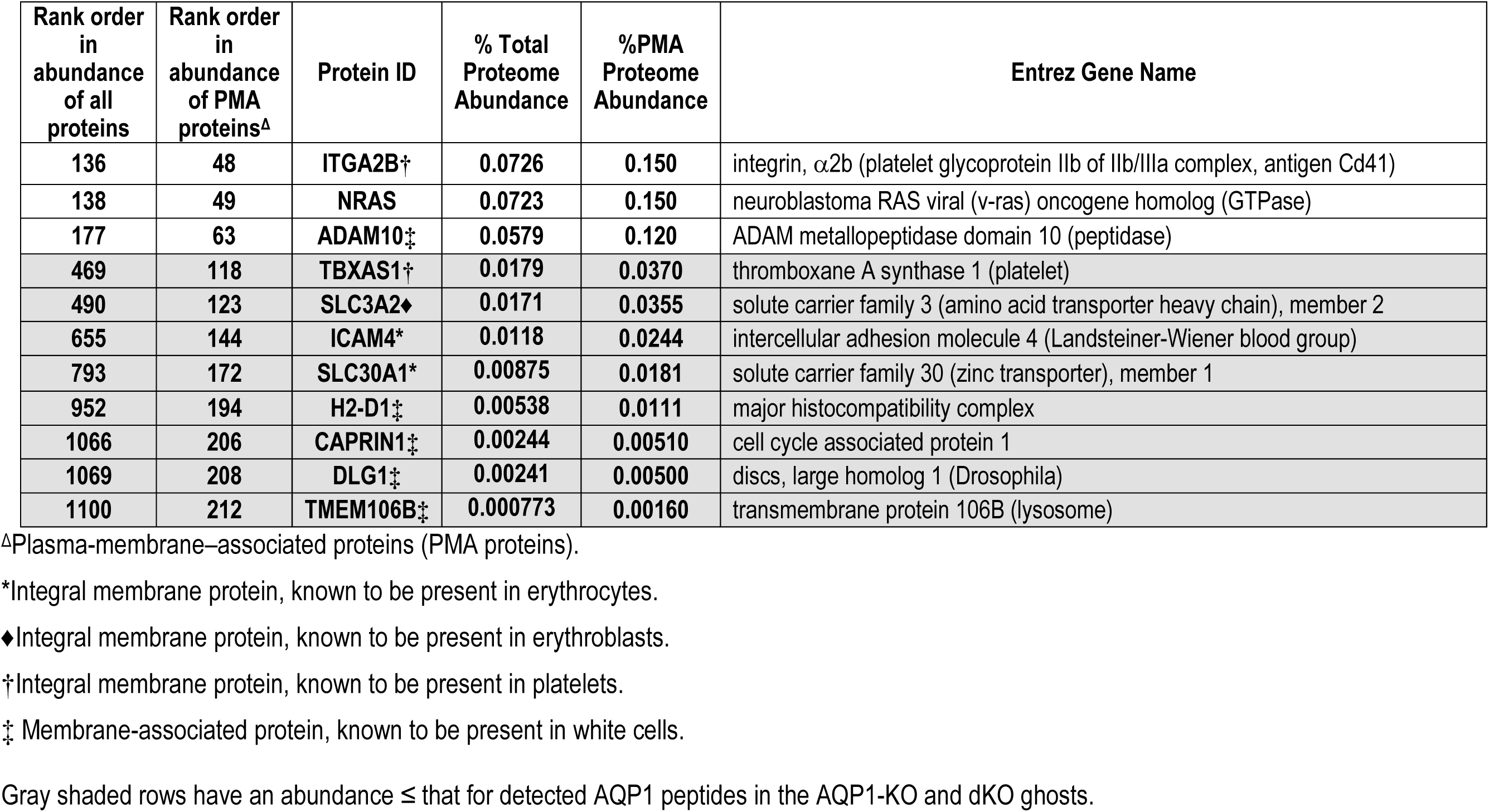
All PMA proteins with significant changes in inferred abundance in RBC ghosts in one or more KO or dKO mouse strains, ranked by abundance in WT ghosts (see Figure 13).

Among PMA proteins exhibiting significant changes, the one with the highest inferred abundance was the integrin ITGA2B (Table 10), which ranks #136 overall (0.07% of all proteins) and #48 (0.15%) among PMA proteins. It is prevalent in platelets, rather than mature RBCs (Nurden, 2021). ITGA2B peptide abundance significantly increases in AQP1-KO and RhAG-KO samples, but decreases in the dKO samples (Figure 11*E*), most likely reflecting small differences in platelet contamination in our ghost samples rather than true changes in ITGA2B expression in erythrocyte membranes. Because the automated hematology in <u>paper #1</u>^39^ reveals no significant differences in platelet numbers among genotypes, the differences observed here could reflect platelet adherence to RBCs.

## Discussion

In this series of three interdependent papers, the present study (<u>paper #2</u>) characterizes the morphologies and proteomes of murine RBCs for our lab-standard WT strain, C57BL/6_Case_, as well as AQP1-KO, RhAG-KO, and dKO mice, all on the same genetic background. We conclude that neither RBC morphology nor PMA protein abundance—aside from the proteins specifically targeted by the KO models—exhibit significant inferred-abundance differences in the KO animals. Thus, we can exclude these parameters as meaningful contributors to the *k*_HbO_2__ differences noted in <u>paper #1</u>. Moreover, the present work provides the necessary experimentally determined parameters for the modeling in <u>paper #3</u>.

In the present work, we use murine RBCs as surrogates for human RBCs to facilitate the genetic manipulation candidate gas-channel proteins. It is important to note that mouse RBCs differ from their human counterparts in several key aspects. Mice have relatively more circulating reticulocytes and platelets than humans (O’Connell *et al*., 2015), and their RBCs have a decidedly smaller MCV (∼47 fl vs. ∼90 fl), an MCHC that is on the lower end of the human range (∼32 g/dl vs. ∼34 g/dl), a smaller Ø_Major_ (∼6.8 μm vs. ∼9.1 μm^40^), and a thickness that is on the lower end of the range for humans (∼2 μm vs. 2.0–2.5 μm). Note, however, that the two critical parameters for gas-exchange by a BCD, and for *k*_HbO_2__ computed in our mathematical simulations—MCHC and thickness—are rather similar between murine and human RBCs. Murine RBCs have a shorter lifespan than human RBCs (55 days vs. 120 days). Finally, the shapes of the RBCs (consistency of biconcavity) vary more in mice than humans (Gilson *et al*., 2009; O’Connell *et al*., 2015). Of course, because mouse RBCs have about half the MCV and only a slightly smaller MCHC compared to humans, the hemoglobin content and oxygen-carrying capacity is much lower in mice than humans (Russell *et al*., 1951; Webb *et al*., 2021).

### Morphometry

#### Morphology

Our work with blood smears (Figure 2) and imaging of living, tumbling RBCs (Figure 4) indicate that the RBCs from mice with genetic deletions in *Aqp1* and or *Rhag* are morphologically indistinguishable from those from WT mice. Our data align with previous reports that erythrocyte morphology is unaltered in AQP1-KO and AQP1/UT-B dKO mice (Yang & Verkman, 2002; Al-Samir *et al*., 2025), and that AQP1-deficient (Colton-null) human RBCs are morphologically identical to *AQP1*+/+ human RBCs (Mathai *et al*., 1996). Similarly, our findings on mice with genetic deletions of *Rhag* align with other microscopic analyses of murine blood that revealed no major morphological differences between of RhAG-KO and WT RBCs (Goossens *et al*., 2010). To our knowledge, we are the first to study the morphology of AQP1/RhAG-dKO RBCs. The prevalence of nBCDs in each genotype, as well as in RBCs treated with pCMBS or DIDS, is addressed below in our discussion of IFC.

#### Imaging flow cytometry for qualitative and quantitative analyses of blood cell morphology

##### Historical background

IFC is highly effective for analyzing the morphology of both living and fixed blood cells. Parasitologists have employed IFC for size-classification of erythrocyte-derived microvesicles in human malaria infections (Mantel *et al*., 2013), screening for compounds that destabilize the *P. falciparum* digestive vacuole (Lee *et al*., 2014), and demonstrating that RBCs are the blood cell type predominantly infected by *Brucella melitensis* (Vitry *et al*., 2014). IFC is also useful for classifying and counting erythroid progenitors to delineate the stages of erythropoiesis (McGrath *et al*., 2017; Kalfa & McGrath, 2018), for characterizing the magnetic susceptibility of human monocytes (Kim *et al*., 2019), or to ascertain whether the substantially elevated platelet counts observed during *ex vivo* lung and liver xenografts originate from RBC fragmentation into platelet-sized particles (Habibabady *et al*., 2022).

Some studies specifically examined the dimensions and morphology of fixed human RBCs. Safeukui *et al*.(2012, 2013) assessed compactness, circularity, aspect ratio, shape ratio and dimensions (surface area and perimeter) under control conditions and after *P. falciparum*-infection or after lysophosphatidylcholine treatment, which induces echinocytosis and, eventually, spherocyte formation. In the above investigations, the distribution of Ø_Major_ values under control conditions centered on 9 to 10 μm. Another group, in a study of patients with severe malaria, used IFC to demonstrate that, after treatment with artesunate induces ejection of the parasite, the RBCs have a projected surface area that 8.9% lower than without treatment (Jauréguiberry et al., 2014).

Some studies use IFC to quantify the proportion of echinocytes, sphero-echinocytes, and spherocytes in stored non-fixed human RBC samples (Pinto *et al*., 2019; Marin *et al*., 2022).

In the present study, we focus on live murine RBCs (not fixed) and—among control cells— find no remarkable differences in RBC morphology, particularly Ø_Major_, between the WT and the three KO mouse strains, by either light-scattering flow-cytometry or IFC. We describe the impact of drug treatments below.

##### nBCD (poikilocyte) abundance

Whereas our major goal in the present IFC study was to determine the mean Ø_Major_ of BCDs from different mouse strains—a critical step in computing RBC thickness—we also used IFC to ascertain nBCD abundance, which we found to be extremely small, both among RBCs from WT mice (∼1.4%) and dKOs (∼2.5%). However, we found that drug pretreatment—followed by long, continuous drug exposure—significantly increases nBCD prevalence, to 8.7% in the case of WT/pCMBS and 41% for WT/DIDS (Table 5). A recent study reported a pre-incubation of murine RBCs with 100 µM DIDS for 10 min at 25 °C (Al-Samir *et al*., 2025), as opposed to 200 µM for ≥90 min in the present investigation. Indeed, in qualitative light-microscopy analyses of samples fixed RBC samples, post-DIDS incubation, they demonstrated substantial DIDS-induced poikilocytosis (mouse > human). However, they neither quantified nBCD prevalence nor accommodated their *k*_HbO_2__ values for the presence of nBCDs. Although these investigators, like us, presumably had a small percentages of nBCDs in their “control” samples, it is unlikely that this had a major impact on the conclusions.

Interestingly, in the present study nBCD prevalence is substantially less in drug-treated RBCs from dKO mice, ∼5.7% for WT/pCMBS, and ∼21% for WT/DIDS.

###### Mechanism of drug action

It has long been supposed that both pCMBS and DIDS are confined to the outside of cells. To our knowledge, <u>paper #1</u>^41^ provides the first experimental evidence of exclusion of both drugs from RBC cytosol. We are unaware of any data implicating either pCMBS or DIDS in interactions with membrane lipids (Rothstein, 1971; Schnell *et al*., 1992), only with proteins (Vansteveninck *et al*., 1965; Rothstein, 1971; Rothstein *et al*., 1973; Cabantchik & Rothstein, 1974*a*). Nevertheless, both drugs are notoriously nonspecific. Vansteveninck *et al*. (1965) concluded that pCMBS interacts only with extracellularly accessible cysteine residues in PMA proteins, which is in stark contrast to the situation for the highly toxic Hg^2+^, which penetrates the membrane and thereby reacts with both extra- and intracellular thiols to drastically alter BCD shape (Notariale *et al*., 2022). Among targets of pCMBS is AQP1 specifically Cys-189 in the extracellular mouth of the monomeric pore, the result of which is inhibition of both H_2_O permeability (Preston *et al*., 1993; Echevarría *et al*., 1993) and CO_2_ permeability (Cooper & Boron, 1998; Musa-Aziz *et al*., 2025).

DIDS, via its two sulfonate groups, can interact ionically with positively-charged regions. Moreover, via the isothiocyano groups on opposite ends of the molecule, DIDS can covalently react with (or even cross-link) extracellular lysine residues on PMA proteins (Rothstein *et al*., 1973; Cabantchik & Rothstein, 1974*a*, 1974*b*; Hsu & Morrison, 1983). A prominent target of DIDS are extracellular-facing lysines on AE1, with the interactions inhibiting conformational transitions required substrate transport (Wong, 1994, 2004; Arakawa *et al*., 2015; Capper *et al*., 2023). Another DIDS target is AQP1; DIDS not only reduces CO_2_ permeability (Boron & Cooper, 1998; Endeward *et al*., 2006; Musa-Aziz *et al*., 2025), but also crosslinks monomers within tetramers (Musa-Aziz *et al*., 2025). In the case of the Rh_Cx_, DIDS actions appear to reduces CO_2_ permeability (Endeward *et al*., 2008).

The substantial decrease in drug-induced nBCD prevalence in dKOs—both for pCMBS and for DIDS treatments—supports is consistent with the hypothesis that interactions of both drugs with AQP1 and/or Rh_Cx_ contribute in a major way to poikilocytosis. It is noteworthy that both AQP1 and Rh_Cx_, as well as AE1, are central members of the RBC ankyrin complex (Vallese *et al*., 2022). Thus, it is possible that pCMBS and DIDS interactions with these and other members of complexes contribute to nBCD formation. Finally, it is also possible that the drugs promote poikilocytosis by inhibiting solute transport.

Inasmuch as all known ankyrin-complex subtypes from human RBCs include the Rh_Cx_, an interesting question is how the RBCs from RhAG-KOs and dKOs are able to maintain their normal biconcave shape.

###### Time dependence

As noted in <u>Results</u> in the present paper, whereas pretreatments with pCMBS or DIDS were complete only ∼5 min before SF experiments, much longer periods elapsed (∼30-90 min) before we were able to visualize RBCs by either microvideography or imaging flow cytometry. Thus, our measures of nBCD prevalence reflect worst-case scenarios for poikilocytosis, and could very well overestimate nBCD prevalence.

###### Accommodation for nBCDs

As described in <u>paper #3</u>^42^, we developed a novel method— based in part on the prevalence of nBCDs, as reported in the present paper—to correct for the presence of nBCDs in our RBC samples. Only in the case of DIDS-treated RBCs are these corrections substantial; for pCMBS-treated RBCs, the corrections are modest; and for control RBCs, the corrections are minimal.

##### Impact of IFC data

To the best of our knowledge, we are the first to employ IFC to compare RBC morphology between WT and different strains of KO mice. The minor degrees of poikilocytosis that we observed in control RBCs, and the larger degrees in drug-treated cells, were presumably present in previous RBC studies. In future investigations, it would be valuable to determine the interrelationships among (1) drug concentration and exposure time, (2) decrease in *P*_M,O_2__, and (3) nBCDs abundance, with the goal of determining protocols that maximize inhibition while minimizing nBCDs formation.

##### RBC geometry cytometry vs in blood vessels

Quantifying RBC morphology “in flow” by IFC is an accurate method to determine Ø_Major_ and nBCD abundance. *In vivo*, RBCs entering capillaries (human diameter, ∼6 ± 2 μm) reversibly adopt a narrower and more elongated bullet-like shape, the more so in narrower capillaries (Secomb, 2017). In 4-μm human capillaries, the “bullet” diameter is estimated to be just under 4 μm. In the smaller murine capillaries (∼4 ± 1 μm; Nicolas & Roux, 2021), it may be reasonable to suggest that the “bullet” diameter can be as small as ∼3 μm, which is ∼50% greater than the diameter of the ∼2-μm diameter equivalent spheres that we use in our mathematical model of the outer rim of a biconcave disk (<u>paper #3</u>^43^).

Thus, not accounting for the lower temperature (i.e., 10°C) of our SF studies vs. body temperature, our mathematical simulations would predict somewhat faster O_2_ offloading in SF experiments than is realistic in small murine capillaries.

A study using high-resolution adaptive optics imaging for non-invasive monitoring of RBCs in the living human retina reports that the computed volumes and surface areas of RBCs fall linearly with capillary diameter (Bedggood *et al*., 2024). Although, in principle, these relationships could reflect pre-existing RBCs morphology, the authors conclude that the RBCs lose considerable volume (presumably H_2_O) as they enter a capillary—volume that they would have to regain upon exiting the capillary. Others suggest that the H_2_O exits in large part via AQP1 (Sugie *et al*., 2018). If this is true, it is not clear why our AQP1-KO and dKO mice appear normal, and how (in preliminary studies^44^) they appear to run well on a treadmill.

##### Non RBCs

We determined that nucleated cells represent ∼1.5% of all cells counted by IFC in WT blood samples, ∼1.7% in AQP1-KO, ∼2.5% in RhAG-KO and ∼2.5% in dKO. These nucleated cells are a mixture of erythroblasts and residual white blood cells (WBCs). In humans, nucleated RBC prevalence is 0–0.5% of WBCs.

Because we preferentially remove WBCs during the early centrifugation steps of our RBC sample preparation, the prevalence of nucleated RBC precursors in our IFC samples may be higher than in whole blood. Note that our gating schemes allow us to exclude the nucleated cells from our calculation of mean Ø_Major_. Note also that because the vast majority of nucleated cells are WBCs—which, unlike reticulocytes and mature RBCs, lack hemoglobin—they do not contribute to our calculations of *k*_HbO_2__ in <u>paper #1</u>^45^ and *P*_M,O_2__ in <u>paper #3</u>^46^.

### Proteomics

#### Historical background

Various proteomic analyses of both human and murine RBC ghosts reveal numerous plasma membrane-associated proteins, some of which likely affect transmembrane O_2_ transport (Kakhniashvili *et al*., 2004; Pasini *et al*., 2006, 2008; Goodman *et al*., 2007; Pesciotta *et al*., 2012, 2012; Bryk & Wiśniewski, 2017; Gautier *et al*., 2018; Vallese *et al*., 2022). These previous studies consistently indicate that AE1 (SLC4A1; Band 3) is the predominant PMA transport protein, followed by the Rh_Cx_ proteins and AQP1 among the next most abundant PMA species.

#### The murine RBC ghost proteome

In our proteomic analyses of the RBC ghosts obtained from the WT and the three KO mouse strains used in the present study, “inferred abundance” from LC/MS/MS data refers to the estimated quantity of specific charged protein fragments in a sample, based on detected peak areas in the mass spectra. The main objective of conducting the proteomics was to confirm that the expression of the target proteins in each KO strain was effectively abolished, and to determine if the gene deletions led to unexpected changes in the RBC proteomes.

##### PMA proteins of low abundance

In our analyses, we focus particularly on the PMA proteins, defined as proteins that are either embedded within the membrane (i.e. integral), or attached to it directly (e.g., via a glycosylphosphatidylinositol anchor) or indirectly (e.g. a cytoskeletal protein such as ankyrin interacting with an integral membrane protein like AE1). For PMA proteins with indirect associations, the amount of protein detected in the ghost samples depends not only on genotype-specific differences in expression level but also on minor variations in wash-step stringency. For high-abundance cytoskeletal proteins like ankyrin (#9 overall rank, #5 among PMA proteins; Table 7) inter-sample variations resulting from minor differences in wash stringency have minimal impact. For low-abundance PMA species, and especially those contributed by contaminating non-RBC cells, minor differences in sample preparation are more likely to have contributed to systematic variability in measured abundance of detected peptides, and thus may have contributed to statistically significant fold-changes. An example is the disks large homolog 1 (DLG1) scaffolding protein (#1069 overall rank, #208 among PMA proteins; Table 10), which is 850-fold less abundant than ANK1, which originates in WBCs.

##### Statistically significance differences in abundance

In all, we detected statistically significant fold-changes in a total of 27 proteins (see Figure 13), 16 non-PMA proteins (Table 9), and 11 PMA associated (Table 10). As a quantitative threshold for considering which of these changes are potentially meaningful, we suggest using the residual AQP1 signal in AQP1-KO and dKO ghosts. For example, the mean AQP1 intensity in WT mice was ∼1.1 × 10^9^ (see Supporting file 1 Tab 7, Cell D6), but only ∼5.2 × 10^6^ in the AQP1-KO (see Supporting file 1 Tab 7, Cell N552).

##### PMA proteins of statistical significance

Only three PMA species (1) had baseline levels in WT ghosts that were above this threshold and (2) exhibited significant genotype-related differences in expression in ≥1 KO sample. Listed in the top three lines of Table 10 and also in Figure 13*E–G* (yellow highlighted panels), these are ITGA2B, NRAS, and ADAM10. However, even these three mostly likely arose from residual platelet, erythroblast, or WBC contamination in the samples, rather than from unusual RBC expression.

Aside from these three proteins, we detected significant genotype-specific changes in eight other PMA proteins—listed in the bottom/gray lines of Table 10 and also in the eight other yellow panels in Figure 13 that where nevertheless below our quantitative threshold. Note that six of these are non-RBC proteins: TBXAS1 is in platelets (see Figure 13*M*), SLC3A2 (see Figure 13*N*) is an acknowledged erythroblast protein, and four others are recognized WBC-specific PMA proteins (see Figure 13*Q,V,W,Z*).

The two remaining PMA proteins of statistical significance are known to express in RBCs (see Table 10): intercellular adhesion molecule 4 (ICAM4; a glycoprotein and the Landsteiner-Wiener antigen, LW; see Figure 13*O*) and SLC30A1, the H^+^-coupled Zn^2+^ exchanger SLC30A1 (ZnT1; see Figure 13*P*). Although ICAM4 is below our abundance threshold, the near-total loss of ICAM4 coincides with the near-total losses of RhAG (Figure 11*G*) and RhD (Figure 11*M*) in both the RhAG-KOs and dKOs. Indeed, ICAM4 is absent from Rh_null_ human RBCs (Hermand *et al*., 1996; Rosa *et al*., 2005). ZnT1 expression is restricted to reticulocytes in mice (Pasini *et al*., 2008; Ryu *et al*., 2008). The 69% increase in ZnT1 abundance in RhAG-KO and the 46% increase in dKO compared to WT ghosts closely corresponds with the minor increases in reticulocyte prevalence observed in the RhAG-KO and dKO IFC samples (Table 2) Overall, we confirm that AQP1 and/or components of the Rh_Cx_ are effectively and appropriately absent in AQP1-KO, RhAG-KO, or dKO RBCs, and that—among the 22 proteins that comprise ∼87% of the PMA protein in WT RBCs—that there are no other significant alterations in abundance (Figure 11). Thus, changes in abundance of membrane proteins other than AQP1 and RhAG/RhD are unlikely to contribute to the observed KO-induced decreases in *k*_HbO_2__ in <u>paper #1</u>.

##### Non-PMA proteins of statistical significance

Determining the relevance of statistically significant differences in non-PMA proteins identified by LC/MS/MS is challenging, inasmuch as we attempted to wash away the cytoplasmic contents from the RBC ghost samples. The 16 non-PMA proteins exhibiting statistically significant genotype-specific alterations constitute < 0.1% of the overall ghost proteome (see Table 9, column 3). The observed differences presumably reflect some combination of sample-to-sample variations in sample preparation and fluctuations in protein expression.

We suggest that soluble proteins could influence channel-mediated gas permeation by at least two mechanisms: (1) protein-protein interactions that either establish one kind of binding or prevent another, and (2) enzymatic activity that—ultimately—could influence channel activity. Given the extremely low abundance of the 16 statistically significant non-PMA proteins, we think that mechanism #1 is highly unlikely. Several of the 16 are enzymes that, if present in RBCs, could directly or indirectly influence *P*_M,O_2__.

#### RBC membrane-protein complexes

##### The ankyrin complex

Vallese *et al*. (2022) determined the structures of several subtypes of the human erythrocyte ankyrin complex, the major RBC PMA protein complex that contributes to the stability and shape of the RBC membrane by tethering the spectrin-actin cytoskeleton to the lipid bilayer. In the present study, we determine that, collectively, murine RBC proteins that correspond to components of the human ankyrin-1 “complex a” comprise ∼62% of all PMA proteins that we detect (see Table 7): AE1 (∼20% of PMA proteins), GYPA (∼19%%), AQP1 (∼8%), ankyrin-1 (ANK1, ∼4%), spectrin β (SPTB, ∼4%), RhAG (∼3%), erythrocyte membrane protein 4.2 (EPB42, ∼3%), and RhD (∼2%).

The human “Class 1a” ankyrin complex (Vallese *et al*., 2022) contains three AE1 homodimers (designated I, II, III), one AQP1 tetramer, one Rh_Cx_ heterotrimer (2×RhAG/1×RhCE), GYPA, GYPB, and protein 4.2 (all of which are integral membrane proteins) as well as spectrin and ankyrin-1, which provide the anchoring to the cytoskeleton. In the Class 1a complex, “AE1-I” interacts via protein 4.2 with ANK1, whereas “AE1-II” and “AE1-II” directly bind ANK1 via their N-termini. The charged AQP1 N-terminal residues K7, K8 and R12 mediate the interactions of AQP1 with ANK1. It is the RhCE that anchors the Rh_Cx_ to ANK1 in the complex via both its N- and C-termini. In murine RBCs, which do not express RhCE, this role may be played by RhD.

Vallese *et al*. (2022) identified six other lower-order ankyrin complex subclasses (Class 1-6), each of which contains one to three AE1 homodimers and one Rh_Cx._ The inclusion of one AQP1 is variable. Classes 1-6 are predicted to occur *in vivo* but they may be less prominent than under the conditions designed to stabilize the interactions for the Cryo-EM study. Vallese *et al*. (2022) suggested that, in mature Class 1a ankyrin complexes, the common spectrin-actin cytoskeletal anchor can transduce mechanical forces among the three AE1 components (I, II, III), AQP1, and Rh_Cx_. It is intriguing to speculate that these interdependent mechanical forces could impact the activities (including *P*_M,O_2__) of these proteins. For example, the cycle of conformational changes of AE1 could impact the activities of AQP1 and Rh_Cx_. Similarly, it is possible that a pharmacological effect on one member of the Class 1a ankyrin complex could indirectly affect other members. However, the nature of both conformational and pharmacological effects could differ among the seven ankyrin-complex subtypes depending on the arrangement of complex members.

If the ankyrin complex(es) of murine RBCs are similar to those of human RBCs, one would expect that the absence of AQP1 in our AQP1-KOs would eliminate all classes containing AQP1: Class 1a (AE1-I/II/III +AQP1 + Rh_Cx_), Class 2 (AE1-I +AQP1 + Rh_Cx_), and Class 5 (AE1-I/III, RhCx, AQP1). The KO of AQP1 would presumably alter the dispositions of the constituent proteins (e.g., isolated Rh_Cx_), including prevalences of the remaining classes—Class 1 (AE1-I/II/III + Rh_Cx_), Class 2 (AE1-I +AQP1 + Rh_Cx_), Class 3 (AE1 +Rh_Cx_), Class 4 (AE1-I/II/III + Rh_Cx_ + an unknown protein), Class 5 (AE1-I/III, RhCx, AQP1), and Class 6 (AE1-I/III Rh_Cx_). Rh_Cx_ is common to all of the seven ankyrin complex classes described by Vallese *et al*., (2022). Thus, in RhAG-KOs, it is unclear how (or if) AQP1, AE1, and the other components of the ankyrin complexes would assemble. The same question pertains to RBCs from dKOs. Presumably some ankyrin complex remain even in dKOs inasmuch as RBCs from these mice have a normal shape (see Figure 4). We suggest that these questions call for a study of ankyrin-complex structures in mice, which would allow the exploitation not only of knockouts, but also of the knockin of designed mutations.

It is probable that the cryo-EM structures did not capture proteins some weaker or more transient interactions with ANK1. For example, CD-47 and ICAM4 are previously recognized constituents of the ankyrin complex (Mankelow *et al*., 2012) that were not detected in the human structures of Vallese *et al*., (2022), who raised the possibilities of loose associations, loss during digitonin extraction, or affiliation with an ankyrin-complex subtype (or other complex type) that has yet to be isolated in structural studies.

In the present study, we detect five unique peptides for CD47, the abundance of which ranks #25 overall and #14 among PMA proteins in our WT mouse RBC proteome (Table 7 and Tab 2 of Supporting file 1). However, we only detect two unique ICAM4 peptides with extremely low abundance (#144 among PMA proteins, Table 10 and Tab 2 of Supporting file 1). The two detected ICAM4 peptides reside within the C-terminal 60% of the 262-residue type-1 protein (residues 105-114 and 191-215), indicating that some of the signal loss may be due to cleavage of a portion of its large extracellular domain during ghost sample washing.

##### The protein 4.1 complex

AE1 and RhAG also link the plasma membrane with the RBC cytoskeleton via interaction with erythrocyte membrane protein 4.1 (EPB41) in a second macromolecular protein complex that includes glycophorin C (GYPC), atypical chemokine receptor 1 (ACKR1; Duffy blood group antigen), the endoplasmic reticulum membrane adapter protein XK, and the zinc endopeptidase KEL, a type-2 glycoprotein that is part of the Kell blood group (Bennett, 1983; Reid *et al*., 1990; Marfatia *et al*., 1994; Nicolas *et al*., 2003; Salomao *et al*., 2008; Suzuki *et al*., 2014). In our murine RBC ghosts, EPB41 is present in almost equal abundance as EPB42 (Table 7)—a component of the ankyrin complex—but is absent from the structures of Vallese *et al*., (2022).

The contributions of each component of the murine “protein 4.1 complex” to the RBC ghost proteome are displayed in Table 7. Collectively, AE1, Rh_Cx_, EPB41, GYPC, XK and KEL comprise 33.7% of the PMA proteins.

The glucose transporter type 1 (GLUT1; SLC2A1), a constituent of the human “protein 4.1 complex” is not detected in mouse RBC ghosts (Rungaldier *et al*., 2013) because mice synthesize their own vitamin C (Chen *et al*., 2012). However, GLUT3 (SLC2A3) is present, constituting approximately 0.1% of the PMA protein.

Besides GYPA, which is an important part of the ankyrin complex, glycophorin C (GYPC)—important, as part of the protein 4.1 complex, in maintaining RBC shape and deformability (Reid *et al*., 1987)—is the only other murine glycophorin detected in the present study. GYPC comprises an additional ∼5% of total PMA protein content (Table 7).

##### Stomatin-associated complexes

Stomatin (STOM; also known as lipid raft protein band 7) is a scaffolding protein that oligomerizes in RBC cholesterol-rich lipid rafts. STOM ranked #15 in murine ghost PMA protein abundance but constitutes only ∼1.5% of the RBC PMA protein proteome (Table 7). STOM (1) upregulates AE1 transporter activity (Genetet *et al*., 2017), (2) indirectly interacts with AQP1 in a cholesterol-dependent manor, (3) interacts with EPB42 and RhD found in ankyrin or protein 4.1 complexes (Rungaldier *et al*., 2013), and (4) interacts with the scaffolding proteins flotillin-1 (FLOT1) and flotillin-2 (FLOT2) (collectively ∼0.016% of murine RBC PMA protein). STOM also could modulate *P*_M,NH3_ via its interactions with both AQP1 and the urea transporter 1 (UT-B; SLC14A1; Rungaldier *et al*., 2013; Geyer *et al*., 2013; Musa-Aziz *et al*., 2025), which constitutes ∼0.8% of RBC PMA protein (Table 7).

#### Other potential gas channels in the RBC membrane

Because the double knockout (dKO) reduces *P*_M,O_2__ by ∼55%, and pCMBS+dKO, reduces *P*_M,O_2__ by ∼91% (<u>paper #1</u>^47^), we consider which proteins or protein combinations in the murine RBC proteome could underlie the unassigned 36% of *P*_M,O_2__. Below we consider membrane proteins with (1) permanent pores or (2) substrate-translocation pathways. We consider such transporters because it is possible that, even if O_2_ cannot permeate via a canonical transmembrane pathway, it might move via transient O_2_-permable “cracks” in the membrane as they cycle through their conformational changes.

##### Other aquaporins

AQP1 is the only AQP detected in mouse RBCs in the present study. Although human RBCs express the aquaglyceroporin AQP3 at ∼30-fold lower levels than AQP1 (Roudier *et al*., 1998), our findings align with previous reports that AQP3 is not functionally important in the water or glycerol permeabilities of mouse RBCs (Yang *et al*., 2001). Thus, together with the observation that AQP3 appears to lack CO_2_ permeability when heterologously expressed in oocytes (Geyer *et al*., 2013*b*), we conclude that it is highly unlikely that another AQP besides AQP1 contributes in a meaningful way to murine RBC *P*_M,O_2_._

##### Solute transporters

Table 8 lists the rank order of the SLC transporters detected in the RBC ghosts. AE1 is by far the most abundant species comprising ∼20% of the PMA protein. AE1 mediates the “chloride shift” (Hamasaki, 1999; Westen & Prange, 2003). In systemic capillaries, following the influx of CO_2_ (mainly via AQP1 and Rh_Cx_ and one or more unknown pathways) into RBCs, nearly 25% of the incoming CO_2_ forms carbamino hemoglobin, shifting the HbO_2_ dissociation curve to the right, and promoting O_2_ offloading and delivery, especially to metabolically active tissues. This is the so-called CO_2_-Bohr effect (for review, see Boron, 2017). Some 70% of the incoming CO_2_ forms 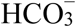 and H^+^, catalyzed by carbonic anhydrases (CAs). This H^+^ binds to HbO_2_ and thereby promotes O_2_ offloading via the much-stronger pH-Bohr effect. Promoting this H^+^ formation are the CAs (by catalyzing the net reaction CO_2_ + H_2_O → 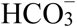 + H^+^) and AE1 (by exporting the newly formed 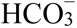 in exchange for Cl^−^). Together, the CAs and AE1 dispose of the newly arriving CO_2_, maximizing the out-to-in CO_2_ gradient, and accelerating CO_2_ influx. The conformation changes (i.e., “elevator mechanism”) that allow the monomers of AE1 and other SLC4 members to alternate between open-to-out and open-to-in conformations (Huynh *et al*., 2018; Zhekova *et al*., 2022) may also provide non-canonical hydrophobic pathways in the transport and scaffold domains of each AE1 monomer similar as hypothesized for CO_2_ via NBCe1 (Moss *et al*., 2019). Thus, it is intriguing to speculate that AE1 could mediate part of the missing ∼35% of *P*_M,O_2__ in murine RBCs. Preliminary results^48^ on RBCs from *Ae1^+/–^*mice are consistent with the idea that AE1 makes such a contribution. Future physiology experiments and molecular dynamic simulations will be required to determine whether AE1 mediates some or all of the remaining *P*_M,O_2__ present in dKO RBCs.

Altogether, the remaining 22 SLC solute transporters in Table 8—excluding AE1, RhAG, and RhD—comprise only ∼5% of all PMA RBC proteins equivalent summed abundance of RhAG and RhD. Like AE1, these species will transport their canonical substrates via elevator or “alternating access” mechanisms, and during this activity, transient O_2_-permable pathways may open between the helices of the core and gate, or transport and scaffold domains of the SLC.

<u>Paper #1</u>^49^ shows that, together, AQP1 and Rh_Cx_ account for ∼55% of *P*_M,O_2__. The present paper shows that, together, AQP1 and Rh_Cx_ account for ∼13% of the inferred abundance of PMA proteins (see Table 7). If we assume that the PMA protein(s) responsible for the missing ∼36% of *P*_M,O_2__ have the same unitary O_2_ conductance (*g*_O_2__) as AQP1 and Rh_Cx_, and that inferred abundance reflect(s) actual protein number, then (36% *P*_M,O_2__/55% *P*_M,O_2__) × (13% PMA proteome) = ∼8.5% of the PMA proteome would mediate the missing ∼36% of *P*_M,O_2__. However, excluding AE1, and Rh_Cx_, only 5% of SLC proteins remain unaccounted for. Thus, either the unaccounted-for SLC proteins have an exceptionally high *g*_O_2__ or we must look somewhere else for the missing ∼36% of *P*_M,O_2__.

##### Ion channels

The PIEZO1 stretch-activated cation channel (Coste *et al*., 2010; Gnanasambandam *et al*., 2015; Fang *et al*., 2021) is not tethered to the cytoskeleton and its distribution in the RBC surface is governed by membrane curvature and tension. PIEZO1 tends to cluster within the concave dimple at the center of RBCs (Vaisey *et al*., 2022). and it is involved in RBC sickling (Chow *et al*., 2021). Missense mutations in *Piezo1* define the Er-blood group (Karamatic Crew *et al*., 2023).

Because PIEZO1 is adapted to gate cations, and is expressed in low abundance (ranking #79 and 0.074% of PMA proteome) relative to other transporters at the RBC plasma membrane, it is an unlikely candidate to directly mediate the missing component of *P*_M,O_2__. However its ion-channel activity and surface distribution is of possible significance in modulating other membrane proteins. The Ca²⁺ fluxes mediated by PIEZO1 in RBCs predominantly initiate various second-messenger signaling pathways. One of the principal pathways entails the activation of calmodulin to regulate cytoskeletal protein interactions in complexes containing AE1, the Rh_Cx_ or AQP1, and facilitate the activation of numerous other signaling pathways (Takakuwa & Mohandas, 1988; Nunomura & Takakuwa, 2006; Nunomura *et al*., 2011). Ca²⁺-influx through PIEZO1 also modulates adenylyl cyclase activity. The resultant modulation in cAMP levels impacts protein kinase A (PKA) activity, resulting in additional downstream effects including but not limited to regulation of RBC volume, deformability, and aggregation (Muravyov & Tikhomirova, 2012).

### Conclusions

In the morphometry part of the present paper, we show that the genetic deficiencies of *Aqp1* and/or *Rhag* do not noticeably alter RBC morphology. Only a very small fraction (∼2%) of control RBCs appear as poikilocytes in imaging flow cytometry. This fraction increases with treatment of WT RBCs with pCMBS and especially DIDS, but much less so in dKOs—shape changes accommodated by novel procedures introduced in <u>paper #3</u>.

In the proteomics part of the present paper, we find that the genetic deficiency of *Aqp1* and/or *Rhag* effectively eliminate the targeted proteins. Specifically, we find no other changes in expression among the 100 proteins with the greatest inferred abundance proteins in expressed in murine RBCs. The most abundant protein to change in at least one knockout strain had an inferred abundance <0.1% of the total proteome, and was not a plasma-membrane–associated protein. The most abundant PMA protein to change in at least one genetic manipulation represents only 0.15% of PMA proteins, and is not even an RBC protein.

We conclude that alterations in RBC morphology cannot account for changes in O_2_ permeability due to genetic manipulations, as reported in <u>paper #1</u>^49^, and at most make a small contribution to computed *P*_M,O_2__ values in pCMBS experiments. The only proteins that could contribute to genetically induced changes in *P*_M,O_2__—a total of 55%—are AQP1 and Rh_Cx_. Although the identity of the protein(s) responsible for the missing ∼36% of *P*_M,O_2__ in <u>paper #1</u>^50^ remains unknown, only a few RBC membrane proteins are sufficiently abundant to be realistic candidates.

## Acknowledgements

We thank Jean-Pierre Cartron for the gift of *Rhag*–/– mice, and for most helpful discussions. Seong-Ki Lee for developing an improved approach for genotyping the *Rhag*–/– mice. We also thank Thomas Radford for organizing the husbandry of the mouse colonies; Gerald Babcock for his role as laboratory manager; Philip G. Woost of the CWRU Flow Cytometry and Imaging Microscopy Core (FCIMC) for his assistance with the flow cytometry data collection and analysis; Daniela Schlatzer of the CWRU Center for Proteomics and Bioinformatics for their assistance with mass spectrometry. This work was supported by Office of Naval Research (ONR) grant N00014-11-1-0889, N00014-14-1-0716, and N00014-15-1-2060; a Multidisciplinary University Research Initiative (MURI) grant N00014-16-1-2535 from the DoD, NIH grant multi-scale modeling grant 5U01GM111251 (to WFB), and NIH grant R01 HL160857 (to W.F.B.). W.F.B. gratefully acknowledges the support of the Myers/Scarpa endowed chair.

## Statistics Tables

**Statistics Tables 1 through 3** Tables of *P*-values for one-way ANOVA with Holm-Bonferroni post-hoc means comparison for comparisons of differences in prevalence of RBCs in Figure 6*D,* reticulocytes in Figure 6*E,* and nucleated cells in Figure 6*F* determined by light-scattering flow cytometry. The tables are split in two halves, with FWER (α) set at 0.05, the upper half shows the adjusted α-value for each comparison and the lower half the *P*-value. If significant, *P*-values are highlighted bold

**Statistics Table 1.** Comparison of RBC prevalence analyzed by determined by light-scattering flow cytometry in Figure 6*D*:

| Mouse |  | WT | AQP1-KO | RhAG-KO | dKO |
| --- | --- | --- | --- | --- | --- |
| | $P$ $\alpha$ | | $\alpha$ | $\alpha$ | $\alpha$ |
| WT |  |  | 0.0500 | 0.00833 | 0.0125 |
| AQP1-KO | *P* | 0.826 |  | 0.0100 | 0.0167 |
| RhAG-KO | *P* | 0.0938 | 0.137 |  | 0.0250 |
| dKO | *P* | 0.299 | 0.407 | 0.476 |  |

**Statistics Table 2.** Comparison of reticulocyte prevalence analyzed by determined by light-scattering flow cytometry in Figure 6*E*.

| Mouse |  | WT | AQP1-KO | RhAG-KO | dKO |
| --- | --- | --- | --- | --- | --- |
| | $P$ $\alpha$ | | $\alpha$ | $\alpha$ | $\alpha$ |
| WT |  |  | 0.0500 | 0.00833 | 0.0125 |
| AQP1-KO | *P* | 0.815 |  | 0.0100 | 0.0167 |
| RhAG-KO | *P* | 0.0909 | 0.136 |  | 0.0250 |
| dKO | *P* | 0.286 | 0.397 | 0.485 |  |

**Statistics Table 3.** Comparison of nucleated cell prevalence analyzed by determined by light-scattering flow cytometry in Figure 6*F*.

| Mouse |  | WT | AQP1-KO | RhAG-KO | dKO |
| --- | --- | --- | --- | --- | --- |
| | $P$ $\alpha$ | | $\alpha$ | $\alpha$ | $\alpha$ |
| WT |  |  | 0.0125 | 0.0500 | 0.0100 |
| AQP1-KO | *P* | 0.716 |  | 0.0167 | 0.0250 |
| RhAG-KO | *P* | 1.00 | 0.716 |  | 0.00833 |
| dKO | *P* | 0.527 | 0.785 | 0.527 |  |

**Statistics Table 4 and Statistics Table 5** Tables of *P*-values for one-way ANOVA with Holm-Bonferroni post-hoc means comparison for comparisons of differences the mean peak FSC-A/FSC-H ratios in Figure 6*I*, and the mean peak FSC-W values in Figure 6*J,* The tables are split in two halves, with FWER (α) set at 0.05, the upper half shows the adjusted α-value for each comparison and the lower half the *P*-value. If significant, *P*-values are highlighted bold.

**Statistics Table 4.**
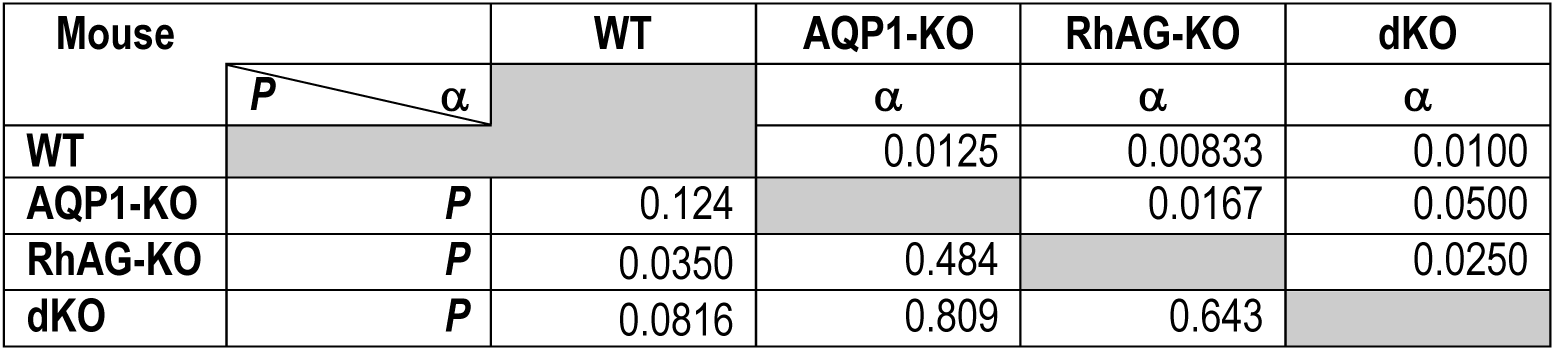
Comparison of the mean peak FSC-A/FSC-H ratios determined by light-scattering flow cytometry in Figure 6*I*.

| Mouse |  | WT | AQP1-KO | RhAG-KO | dKO |
| --- | --- | --- | --- | --- | --- |
| | $P$ $\alpha$ | | $\alpha$ | $\alpha$ | $\alpha$ |
| WT |  |  | 0.0125 | 0.00833 | 0.0100 |
| AQP1-KO | *P* | 0.124 |  | 0.0167 | 0.0500 |
| RhAG-KO | *P* | 0.0350 | 0.484 |  | 0.0250 |
| dKO | *P* | 0.0816 | 0.809 | 0.643 |  |

**Statistics Table 5.** Comparison of nucleated cell prevalence analyzed by determined by light-scattering flow cytometry in Figure 6*J*.

| Mouse |  | WT | AQP1-KO | RhAG-KO | dKO |
| --- | --- | --- | --- | --- | --- |
| | $P$ $\alpha$ | | $\alpha$ | $\alpha$ | $\alpha$ |
| WT |  |  | 0.0125 | 0.00833 | 0.0100 |
| AQP1-KO | *P* | 0.124 |  | 0.0167 | 0.0500 |
| RhAG-KO | *P* | 0.349 | 0.484 |  | 0.0250 |
| dKO | *P* | 0.0814 | 0.809 | 0.643 |  |

**Statistics Table 6 through Statistics Table 8** Tables of *P*-values for one-way ANOVA with Holm-Bonferroni post-hoc means comparison for comparisons of differences in prevalence of RBC (Statistics Table 6), reticulocytes (Statistics Table 7) and nucleated cells (Statistics Table 8) in IFC samples in Figure 7*H*. The tables are split in two halves, with FWER (α) set at 0.05, the upper half shows the adjusted α-value for each comparison and the lower half the *P*-value. If significant, *P*-values are highlighted bold.

**Statistics Table 6.** Comparison of RBC prevalence analyzed by IFC.

| Mouse |  | WT | AQP1-KO | RhAG-KO | dKO |
| --- | --- | --- | --- | --- | --- |
| | $P$ $\alpha$ | | $\alpha$ | $\alpha$ | $\alpha$ |
| WT |  |  | 0.0500 | 0.00833 | 0.0125 |
| AQP1-KO | $P$ | 0.240 | | 0.0100 | 0.0500 |
| RhAG-KO | $P$ | **$3.87 \times 10^{-4}$** | **0.00345** | | 0.0167 |
| dKO | $P$ | **0.00689** | 0.0665 | 0.133 | |

**Statistics Table 7.** Comparison of reticulocyte prevalence analyzed by IFC.

| Mouse |  | WT | AQP1-KO | RhAG-KO | dKO |
| --- | --- | --- | --- | --- | --- |
| | $P$ $\alpha$ | | $\alpha$ | $\alpha$ | $\alpha$ |
| WT |  |  | 0.0250 | 0.00833 | 0.0125 |
| AQP1-KO | $P$ | 0.396 | | 0.0100 | 0.0500 |
| RhAG-KO | $P$ | 0.0125 | 0.0625 | | 0.0167 |
| dKO | $P$ | 0.160 | 0.547 | 0.177 | |

**Statistics Table 8.** Comparison of nucleated cell prevalence analyzed by IFC.

| Mouse |  | WT | AQP1-KO | RhAG-KO | dKO |
| --- | --- | --- | --- | --- | --- |
| | $P$ $\alpha$ | | $\alpha$ | $\alpha$ | $\alpha$ |
| WT |  |  | 0.0250 | 0.0100 | 0.00833 |
| AQP1-KO | $P$ | 0.728 | | 0.0167 | 0.0125 |
| RhAG-KO | $P$ | 0.0415 | 0.0782 | | 0.0500 |
| dKO | $P$ | 0.0362 | 0.0686 | 0.942 | |

**Statistics Table 9 through Statistics Table 31** Tables of *P*-values for one-way ANOVA with Holm-Bonferroni post-hoc means comparison for comparisons of differences in inferred abundance of the SLC transporter proteins listed in Table 8 between each of the three knockout mouse strains and WT. The table is split in two halves, with FWER set at 0.05, the upper half shows the adjusted α-value for each comparison and the lower half the *P*-value. Significant *P*-values are highlighted bold.

**Statistics Table 9.** Comparison of SLC4A1 (AE1) inferred abundance.

| Mouse |  | WT | AQP1-KO | RhAG-KO | dKO |
| --- | --- | --- | --- | --- | --- |
| | $P$ $\alpha$ | | $\alpha$ | $\alpha$ | $\alpha$ |
| WT |  |  | 0.0100 | 0.0500 | 0.0250 |
| AQP1-KO | *P* | 0.760 |  | 0.0167 | 0.00833 |
| RhAG-KO | *P* | 0.869 | 0.848 |  | 0.0125 |
| dKO | *P* | 0.869 | 0.638 | 0.780 |  |

**Statistics Table 10.** Comparison of SLC42A1 (RhAG) inferred abundance.

| Mouse |  | WT | AQP1-KO | RhAG-KO | dKO |
| --- | --- | --- | --- | --- | --- |
| | $P$ $\alpha$ | | $\alpha$ | $\alpha$ | $\alpha$ |
| WT |  |  | 0.0250 | 0.0167 | 0.0125 |
| AQP1-KO | *P* | 0.898 |  | 0.0100 | 0.00833 |
| RhAG-KO | *P* | **0.00385** | **0.00269** |  | 0.0500 |
| dKO | *P* | **0.00365** | **0.00255** | 0.985 |  |

**Statistics Table 11.** Comparison of SLC16A1 (MCT1) inferred abundance.

| Mouse |  | WT | AQP1-KO | RhAG-KO | dKO |
| --- | --- | --- | --- | --- | --- |
| | $P$ $\alpha$ | | $\alpha$ | $\alpha$ | $\alpha$ |
| WT |  |  | 0.00833 | 0.0100 | 0.0125 |
| AQP1-KO | *P* | 0.605 |  | 0.0250 | 0.0167 |
| RhAG-KO | *P* | 0.673 | 0.924 |  | 0.0500 |
| dKO | *P* | 0.736 | 0.856 | 0.932 |  |

**Statistics Table 12.** Comparison of SLC42A5 (RhD) inferred abundance.

| Mouse |  | WT | AQP1-KO | RhAG-KO | dKO |
| --- | --- | --- | --- | --- | --- |
| | $P$ $\alpha$ | | $\alpha$ | $\alpha$ | $\alpha$ |
| WT |  |  | 0.0250 | 0.0100 | 0.00833 |
| AQP1-KO | *P* | 0.821 |  | 0.0167 | 0.0125 |
| RhAG-KO | *P* | 0.0103 | 0.0187 |  | 0.0500 |
| dKO | *P* | 0.00980 | 0.00179 | 0.986 |  |

**Statistics Table 13.** Comparison of SLC43A1 (LAT3) inferred abundance.

| Mouse |  | WT | AQP1-KO | RhAG-KO | dKO |
| --- | --- | --- | --- | --- | --- |
| | $P$ $\alpha$ | | $\alpha$ | $\alpha$ | $\alpha$ |
| WT |  |  | 0.0100 | 0.0500 | 0.0250 |
| AQP1-KO | $P$ | 0.783 | | 0.0167 | 0.00833 |
| RhAG-KO | $P$ | 0.928 | 0.853 | | 0.0125 |
| dKO | $P$ | 0.892 | 0.681 | 0.820 | |

**Statistics Table 14.**
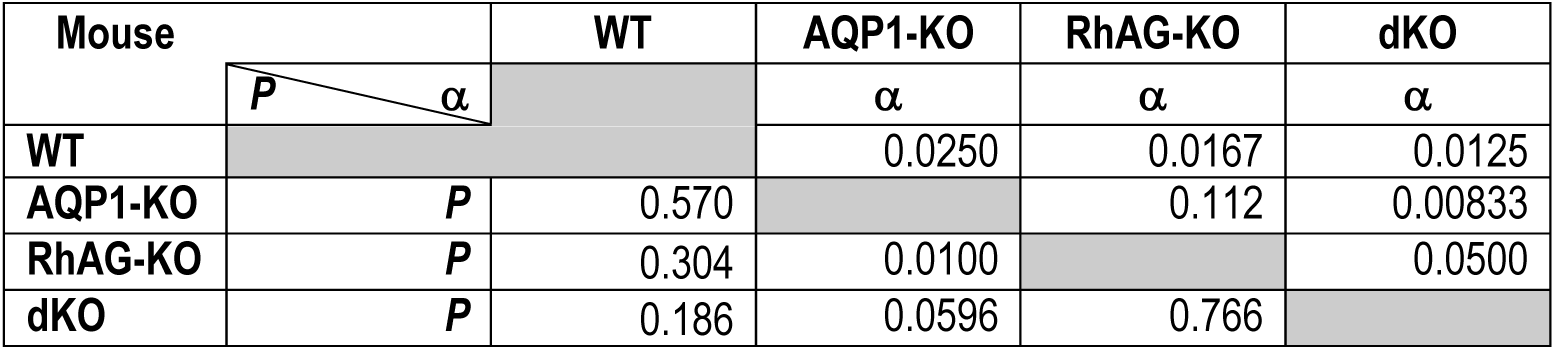
Comparison of SLC14A1 (UT-B) inferred abundance.

**Statistics Table 15.** Comparison of SLC16A10 (MCT10) inferred abundance.

| Mouse |  | WT | AQP1-KO | RhAG-KO | dKO |
| --- | --- | --- | --- | --- | --- |
| | $P$ $\alpha$ | | $\alpha$ | $\alpha$ | $\alpha$ |
| WT |  |  | 0.0167 | 0.0500 | 0.0100 |
| AQP1-KO | $P$ | 0.811 | | 0.0250 | 0.0125 |
| RhAG-KO | $P$ | 0.977 | 0.814 | | 0.00833 |
| dKO | $P$ | 0.539 | 0.675 | 0.513 | |

**Statistics Table 16.** Comparison of SLC29A1 (ENT1) inferred abundance.

| Mouse |  | WT | AQP1-KO | RhAG-KO | dKO |
| --- | --- | --- | --- | --- | --- |
| | $P$ $\alpha$ | | $\alpha$ | $\alpha$ | $\alpha$ |
| WT |  |  | 0.0125 | 0.0250 | 0.0167 |
| AQP1-KO | $P$ | 0.568 | | 0.0100 | 0.00833 |
| RhAG-KO | $P$ | 0.855 | 0.451 | | 0.0500 |
| dKO | $P$ | 0.579 | 0.261 | 0.710 | |

**Statistics Table 17.** Comparison of SLC29A2 (ENT2) inferred abundance.

| Mouse |  | WT | AQP1-KO | RhAG-KO | dKO |
| --- | --- | --- | --- | --- | --- |
| | $P$ $\alpha$ | | $\alpha$ | $\alpha$ | $\alpha$ |
| WT |  |  | 0.0500 | 0.010 | 0.0250 |
| AQP1-KO | $P$ | 0.846 | | 0.00833 | 0.0167 |
| RhAG-KO | $P$ | 0.330 | 0.251 | | 0.0125 |
| dKO | $P$ | 0.751 | 0.611 | 0.0498 | |

**Statistics Table 18.**
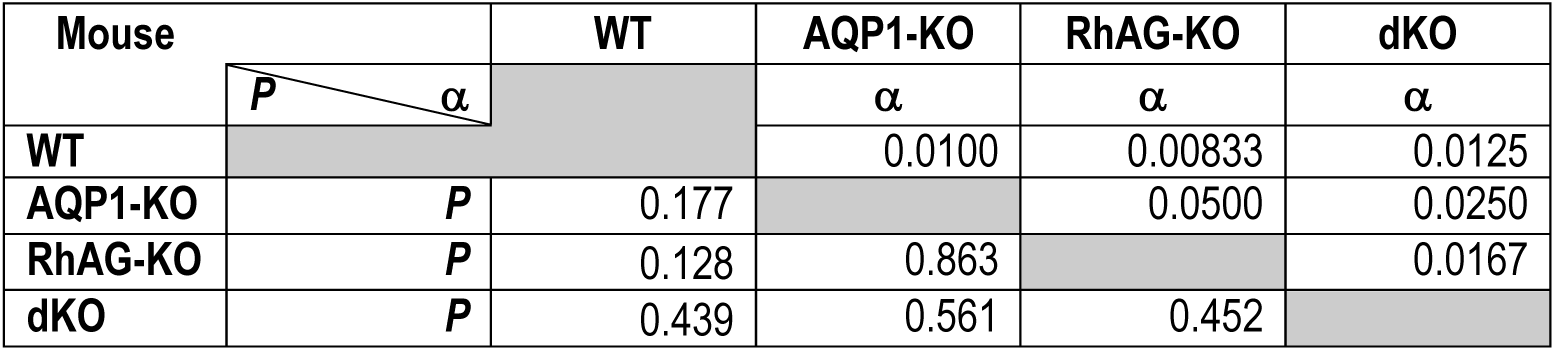
Comparison of SLC40A10 (MTP1) inferred abundance.

| Mouse |  | WT | AQP1-KO | RhAG-KO | dKO |
| --- | --- | --- | --- | --- | --- |
| | | | $\alpha$ | $\alpha$ | $\alpha$ |
| WT |  |  | 0.0100 | 0.00833 | 0.0125 |
| AQP1-KO | *P* | 0.177 |  | 0.0500 | 0.0250 |
| RhAG-KO | *P* | 0.128 | 0.863 |  | 0.0167 |
| dKO | *P* | 0.439 | 0.561 | 0.452 |  |

**Statistics Table 19.**
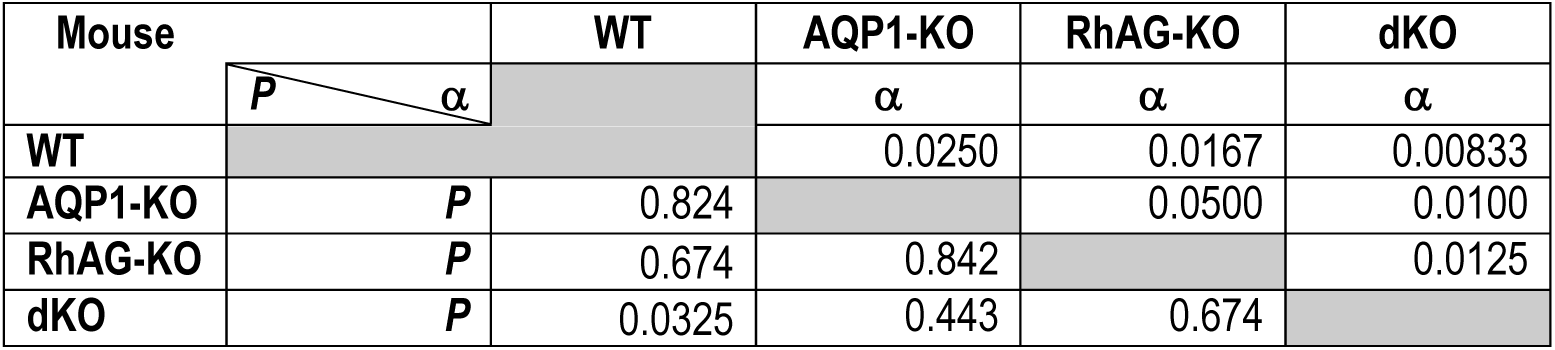
Comparison of SLC4A4 (NBCe1A) inferred abundance.

| Mouse |  | WT | AQP1-KO | RhAG-KO | dKO |
| --- | --- | --- | --- | --- | --- |
| | | | $\alpha$ | $\alpha$ | $\alpha$ |
| WT |  |  | 0.0250 | 0.0167 | 0.00833 |
| AQP1-KO | *P* | 0.824 |  | 0.0500 | 0.0100 |
| RhAG-KO | *P* | 0.674 | 0.842 |  | 0.0125 |
| dKO | *P* | 0.0325 | 0.443 | 0.674 |  |

**Statistics Table 20.**
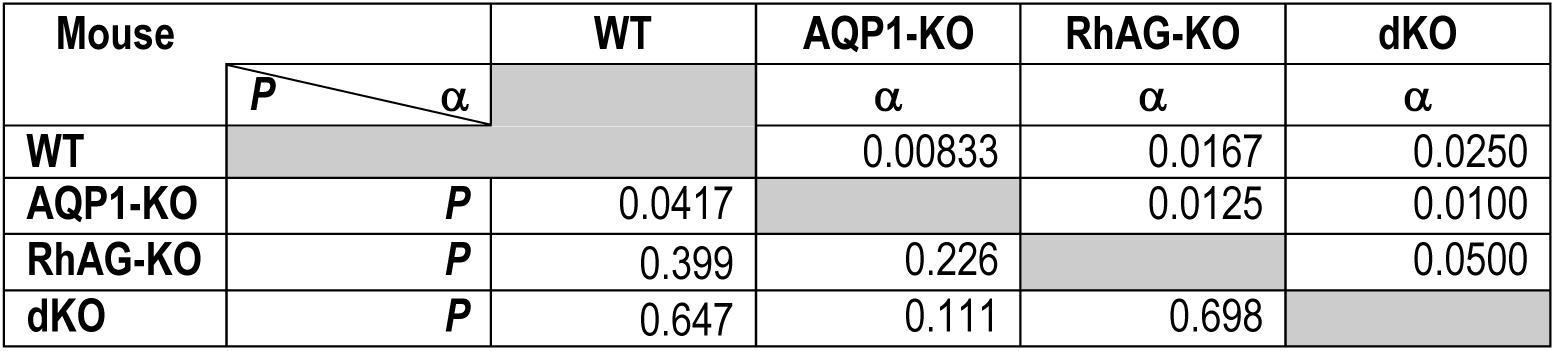
Comparison of SLC2A3 (GLUT3) inferred abundance.

| Mouse |  | WT | AQP1-KO | RhAG-KO | dKO |
| --- | --- | --- | --- | --- | --- |
| | | | $\alpha$ | $\alpha$ | $\alpha$ |
| WT |  |  | 0.00833 | 0.0167 | 0.0250 |
| AQP1-KO | *P* | 0.0417 |  | 0.0125 | 0.0100 |
| RhAG-KO | *P* | 0.399 | 0.226 |  | 0.0500 |
| dKO | *P* | 0.647 | 0.111 | 0.698 |  |

**Statistics Table 21.**
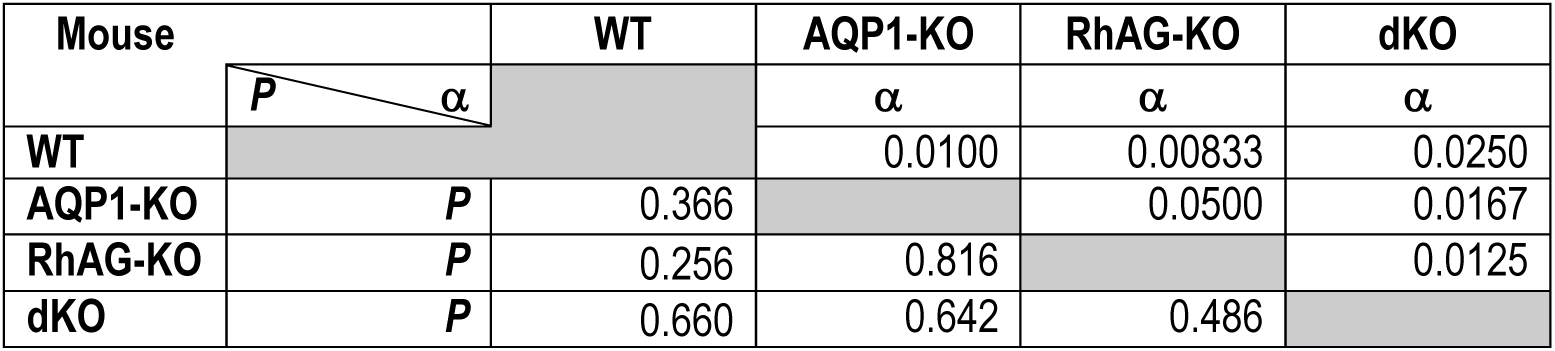
Comparison of SLC9A1 (NHE1) inferred abundance.

| Mouse |  | WT | AQP1-KO | RhAG-KO | dKO |
| --- | --- | --- | --- | --- | --- |
| | | | $\alpha$ | $\alpha$ | $\alpha$ |
| WT |  |  | 0.0100 | 0.00833 | 0.0250 |
| AQP1-KO | *P* | 0.366 |  | 0.0500 | 0.0167 |
| RhAG-KO | *P* | 0.256 | 0.816 |  | 0.0125 |
| dKO | *P* | 0.660 | 0.642 | 0.486 |  |

**Statistics Table 22.**
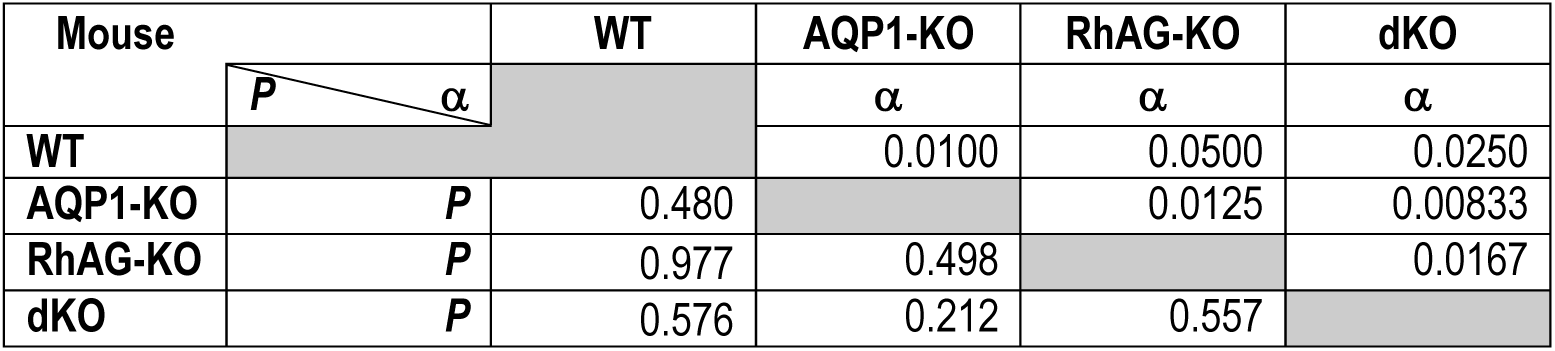
Comparison of SLC26A1 (SAT1) inferred abundance.

| Mouse |  | WT | AQP1-KO | RhAG-KO | dKO |
| --- | --- | --- | --- | --- | --- |
| | | | $\alpha$ | $\alpha$ | $\alpha$ |
| WT |  |  | 0.0100 | 0.0500 | 0.0250 |
| AQP1-KO | *P* | 0.480 |  | 0.0125 | 0.00833 |
| RhAG-KO | *P* | 0.977 | 0.498 |  | 0.0167 |
| dKO | *P* | 0.576 | 0.212 | 0.557 |  |

**Statistics Table 23.**
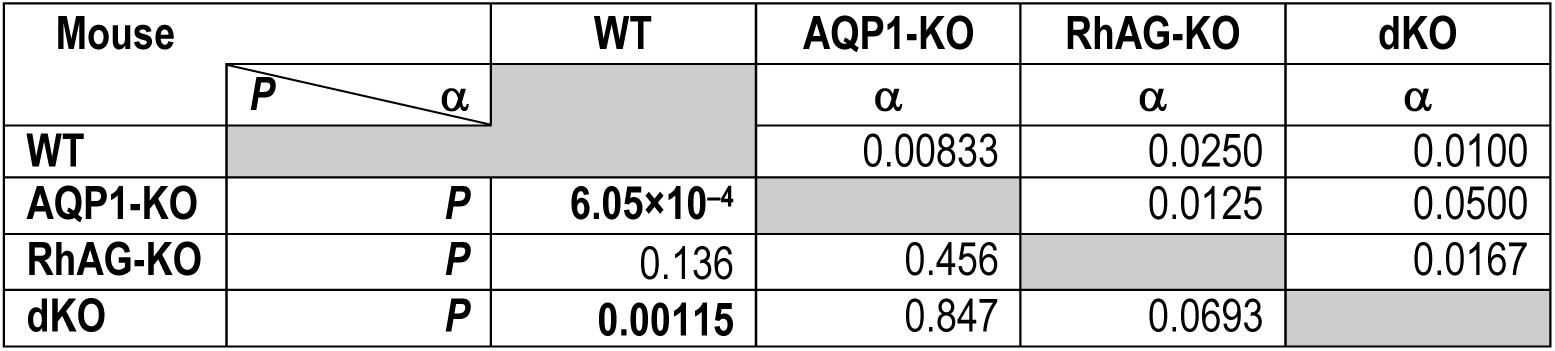
Comparison of SLC3A2 (MDU1) inferred abundance.

| Mouse |  | WT | AQP1-KO | RhAG-KO | dKO |
| --- | --- | --- | --- | --- | --- |
| | | | $\alpha$ | $\alpha$ | $\alpha$ |
| WT |  |  | 0.00833 | 0.0250 | 0.0100 |
| AQP1-KO | *P* | $6.05 \times 10^{-4}$ | | 0.0125 | 0.0500 |
| RhAG-KO | *P* | 0.136 | 0.456 |  | 0.0167 |
| dKO | *P* | 0.00115 | 0.847 | 0.0693 |  |

**Statistics Table 24.**
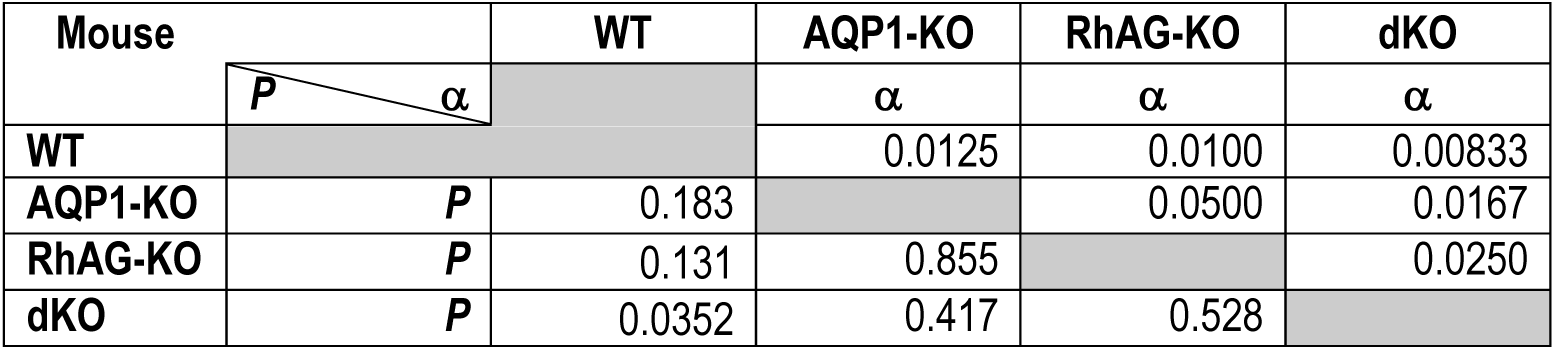
Comparison of SLC11A2 (DMT1) inferred abundance.

| Mouse |  | WT | AQP1-KO | RhAG-KO | dKO |
| --- | --- | --- | --- | --- | --- |
| | | | $\alpha$ | $\alpha$ | $\alpha$ |
| WT |  |  | 0.0125 | 0.0100 | 0.00833 |
| AQP1-KO | *P* | 0.183 |  | 0.0500 | 0.0167 |
| RhAG-KO | *P* | 0.131 | 0.855 |  | 0.0250 |
| dKO | *P* | 0.0352 | 0.417 | 0.528 |  |

**Statistics Table 25.**
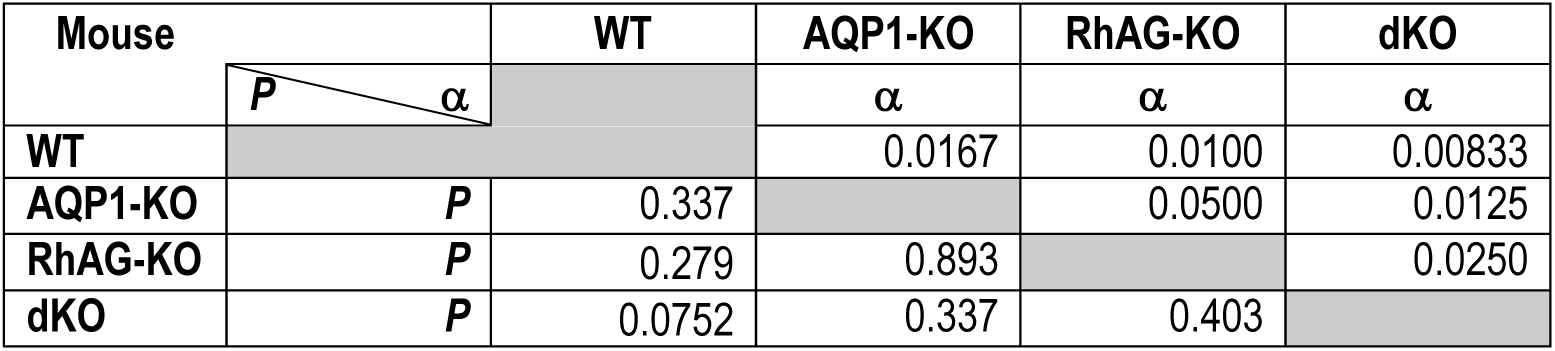
Comparison of SLC7A1 (CAT1) inferred abundance.

| Mouse |  | WT | AQP1-KO | RhAG-KO | dKO |
| --- | --- | --- | --- | --- | --- |
| | | | $\alpha$ | $\alpha$ | $\alpha$ |
| WT |  |  | 0.0167 | 0.0100 | 0.00833 |
| AQP1-KO | *P* | 0.337 |  | 0.0500 | 0.0125 |
| RhAG-KO | *P* | 0.279 | 0.893 |  | 0.0250 |
| dKO | *P* | 0.0752 | 0.337 | 0.403 |  |

**Statistics Table 26.**
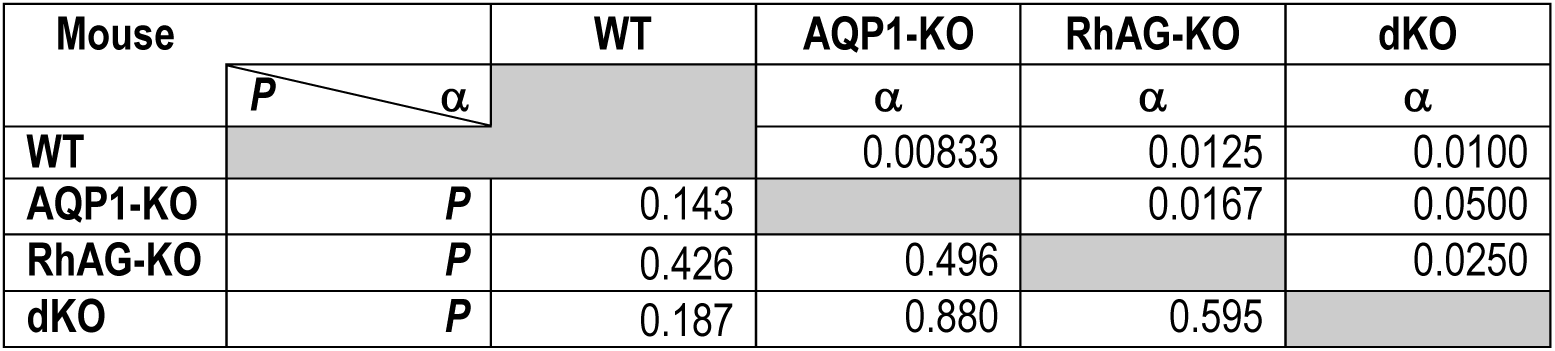
Comparison of SLC7A5 (LAT1) inferred abundance.

| Mouse |  | WT | AQP1-KO | RhAG-KO | dKO |
| --- | --- | --- | --- | --- | --- |
| | | | $\alpha$ | $\alpha$ | $\alpha$ |
| WT |  |  | 0.00833 | 0.0125 | 0.0100 |
| AQP1-KO | *P* | 0.143 |  | 0.0167 | 0.0500 |
| RhAG-KO | *P* | 0.426 | 0.496 |  | 0.0250 |
| dKO | *P* | 0.187 | 0.880 | 0.595 |  |

**Statistics Table 27.**
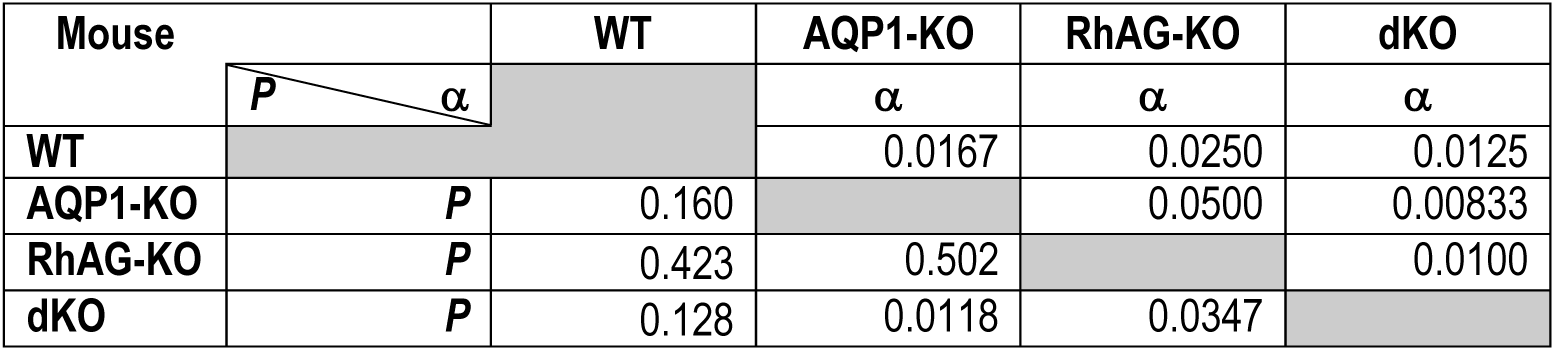
Comparison of SLC6A4 (SERT) inferred abundance.

| Mouse |  | WT | AQP1-KO | RhAG-KO | dKO |
| --- | --- | --- | --- | --- | --- |
| | | | $\alpha$ | $\alpha$ | $\alpha$ |
| WT |  |  | 0.0167 | 0.0250 | 0.0125 |
| AQP1-KO | *P* | 0.160 |  | 0.0500 | 0.00833 |
| RhAG-KO | *P* | 0.423 | 0.502 |  | 0.0100 |
| dKO | *P* | 0.128 | 0.0118 | 0.0347 |  |

**Statistics Table 28.** Comparison of SLC6A9 (GLYT1) inferred abundance.

| Mouse |  | WT | AQP1-KO | RhAG-KO | dKO |
| --- | --- | --- | --- | --- | --- |
| | $P$ $\alpha$ | | $\alpha$ | $\alpha$ | $\alpha$ |
| WT |  |  | 0.0167 | 0.0100 | 0.00833 |
| AQP1-KO | $P$ | 0.385 | | 0.0250 | 0.0125 |
| RhAG-KO | $P$ | 0.108 | 0.436 | | 0.0500 |
| dKO | $P$ | 0.0696 | 0.315 | 0.817 | |

**Statistics Table 29.** Comparison of SLC44A1 (CD92) inferred abundance.

| Mouse |  | WT | AQP1-KO | RhAG-KO | dKO |
| --- | --- | --- | --- | --- | --- |
| | $P$ $\alpha$ | | $\alpha$ | $\alpha$ | $\alpha$ |
| WT |  |  | 0.0167 | 0.0125 | 0.0250 |
| AQP1-KO | $P$ | 0.782 | | 0.00833 | 0.0500 |
| RhAG-KO | $P$ | 0.160 | 0.0958 | | 0.0100 |
| dKO | $P$ | 0.857 | 0.923 | 0.115 | |

**Statistics Table 30.** Comparison of SLC30A1 (ZnT1) inferred abundance.

| Mouse |  | WT | AQP1-KO | RhAG-KO | dKO |
| --- | --- | --- | --- | --- | --- |
| | $P$ $\alpha$ | | $\alpha$ | $\alpha$ | $\alpha$ |
| WT |  |  | 0.0500 | 0.00833 | 0.0125 |
| AQP1-KO | $P$ | 0.234 | | 0.0100 | 0.0167 |
| RhAG-KO | $P$ | $1.09 \times 10^{-04}$ | 0.00393 | | 0.0250 |
| dKO | $P$ | 0.00711 | 0.113 | 0.161 | |

**Statistics Table 31.** Comparison of SLC26A3 (DRA) inferred abundance.

| Mouse |  | WT | AQP1-KO | RhAG-KO | dKO |
| --- | --- | --- | --- | --- | --- |
| | $P$ $\alpha$ | | $\alpha$ | $\alpha$ | $\alpha$ |
| WT |  |  | 0.00833 | 0.0100 | 0.0250 |
| AQP1-KO | $P$ | 0.389 | | 0.0500 | 0.0125 |
| RhAG-KO | $P$ | 0.394 | 0.993 | | 0.0167 |
| dKO | $P$ | 0.668 | 0.654 | 0.660 | |

## Video Legends

Click to see Supporting Video #1, WT RBCs

**Video 1. Video clip showing shapes of RBCs from a WT mouse**

The video (1 frame per 5 s) follows RBCs (collected and prepared as described in <u>Methods</u>^54^, and used at an Hct of 2.5% to 3.0% as described in <u>Methods</u>^55^) of an RBC droplet as they fall freely through the plane of focus (40× objective, NA 1.35, with a 1.5× magnification selector), toward the surface of the coverslip. In the microvideo, the numerals that identify 18 specific RBCs are red when the cells are approximately in focus, but yellow either before they have fallen into focus or after they have fallen out of focus. This microvideo, which is typical of similar microvideos on a total of 3 mice, shows that the RBCs from WT mice are predominantly biconcave disks.

Click to see Supporting Video #2, *Aqp1*-/- RBCs

**Video 2. Video clip showing shapes of RBCs from an *Aqp1*–/– (i.e., AQP1-KO) mouse**

The video (1 frame per 5 s) follows RBCs (collected and prepared as described in <u>Methods</u>^56^, and used at an Hct of 2.5% to 3.0% as described in <u>Methods</u>^57^) of an RBC droplet as they fall freely through the plane of focus (40× objective, NA 1.35, with a 1.5× magnification selector), toward the surface of the coverslip. In the microvideo, the numerals that identify 14 specific RBCs are red when the cells are approximately in focus, but yellow either before they have fallen into focus or after they have fallen out of focus. This microvideo, which is typical of similar microvideos on a total of 3 mice, shows that the RBCs from AQP-KO mice are predominantly biconcave disks. Thus, one cannot attribute the observed 9% decrease in *k*_HbO_2__ value to a change in RBC shape.

Click to see Supporting Video #3, *Rhag*-/- RBCs

**Video 3. Video clip showing shapes of RBCs from an *Rhag*–/– (i.e., RhAG-KO) mouse**

The video (1 frame per 5 s) follows RBCs (collected and prepared as described in <u>Methods</u>^58^, and used at an Hct of 2.5% to 3.0% as described in <u>Methods</u>^59^) of an RBC droplet as they fall freely through the plane of focus (40× objective, NA 1.35, with a 1.5× magnification selector), toward the surface of the coverslip. In the microvideo, the numerals that identify 13 specific RBCs are red when the cells are approximately in focus, but yellow either before they have fallen into focus or after they have fallen out of focus. This microvideo, which is typical of similar microvideos on a total of 3 mice, shows that the RBCs from RhAG-KO mice are predominantly biconcave disks. Thus, one cannot attribute the observed 17% decrease in *k*_HbO_2__ value to a change in RBC shape.

Click to see Supporting Video #4, dKO RBCs

**Video 4. Video clip showing shapes of RBCs from a *Aqp1*–/–*Rhag–*/– (i.e., dKO) mouse**

The video (1 frame per 5 s) follows RBCs (collected and prepared as described in <u>Methods</u>^60^, and used at an Hct of 2.5% to 3.0% as described in <u>Methods</u>^61^) of an RBC droplet as they fall freely through the plane of focus (40× objective, NA 1.35, with a 1.5× magnification selector), toward the surface of the coverslip. In the microvideo, the numerals that identify 15 specific RBCs are red when the cells are approximately in focus, but yellow either before they have fallen into focus or after they have fallen out of focus. This microvideo, which is typical of similar microvideos on a total of 3 mice, shows that the RBCs from dKO mice are predominantly biconcave disks. Thus, one cannot attribute the observed 31% decrease in *k*_HbO_2__ value to a change in RBC shape.

## Additional Information

### Data availability statement

The data that support the findings of this study are available from the corresponding author. W.F.B., upon reasonable request.

### Author contributions

F.J.M., P.Z., & W.F.B. designed the study; F.J.M. performed mass spectrometry; F.J.M also performed the flow cytometry experiments and analyzed the data together with J.W.J.; P.Z., P.Z. performed automated hematology and analyzed the data with F.J.M.; A.I.S. & S.T. performed the microvideography of tumbling blood cells and analyzed the data with D.E.H who assisted in capturing still images from videos, P.Z. coordinated the microvideography experiments of red blood cells treated with inhibitors; A.B.W. produced ghosts for mass spectrometry; P.Z. made blood smears and H.J.M. performed the analysis; F.J.M., P.Z., R.O. & W.F.B. wrote the manuscript.

## Supporting Information

### Supporting Files

**Supporting file 1:** 2--MassSpecData.xlsx

**Supporting file 2:** 2--Suppl_Tables_MassSpec_Statistics_ForTable3.xlsx

**Supporting file 3:** 2--Suppl_Tables_MassSpec_Statistics_ForTable4&Table6.xlsx

**Supporting file 4:** 2--Suppl_Tables_MassSpec_Statistics_ForFig10&11&Table7.xlsx

**Supporting file 5:** 2--Suppl_Tables_MassSpec_Statistics_ForTable8.xlsx

**Supporting file 6:** 2--Suppl_Tables_MassSpec_Statistics_ForFig13_Tables9&10.xlsx

### Supporting Videos

**Supporting video 1:** 2--Supp_Video_1--WT.wmv

**Supporting video 2:** 2--Supp_Video_2--AQP1-KO.wmv

**Supporting video 3:** 2--Supp_Video_3--RhAG-KO.wmv

**Supporting video 4:** 2--Supp_Video_4--dKO.wmv

## Footnotes

1 As a shorthand, we will refer to the three papers as “paper #1”, “paper #2”, and “paper #3”.

2 This manuscript was first published as a preprint: Moss FJ, P Zhao, AI Salameh, S Taki, AB Wass, JW Jacobberger, DE Huffman, HJ Meyerson, R Occhipinti & WF Boron (2025). Role of channels in the O₂ permeability of murine red blood cells II. Morphological and proteomic studies. bioRxiv. https://doi.org/10.1101/2025.03.05.639962

3 See Paper #1>Methods>Mice. To aid the reader in locating the precise target, we use the “>” symbol to separate heading levels. It is understood that the reference is to a location in the present paper unless “See” is followed by “Paper #1” or “Paper #3”.

4 See Paper #1>Methods>Genotyping

5 See Paper #1>table 2

6 See Paper #1>Methods>Inhibitors

7 See Paper #1>Methods>Preparation of RBCs

8 See Paper #1>Methods>Preparation of RBCs>Processing of RBCs for assays 1–4

9 See Paper #1>Methods>Preparation of RBCs>Processing of RBCs for stopped-flow studies

10 See Methods>Still microphotography and microvideography of living RBCs

11 See Methods>Flow cytometry (workflow #6 & 6′, #15 & 15′)

12 See Paper #1>Methods>Automated hematological analyses (workflow #7 – #9)

13 See Methods>Blood smears

14 See Methods>Proteomic analysis

15 See Methods>Preparation of RBCs>Collection of blood

16 See Methods>Preparation of RBCs>Collection of blood from mice.>Processing of RBCs for assays 1–4

17 See Results>Morphometry>Light-scattering flow cytometry

18 See Methods> Flow cytometry (workflow #6 & #6′, #15 & #15′)>Sample preparation

19 See Results>Morphometry>Imaging flow cytometry (IFC)

20 See Paper #1>Methods>Automated hematological analyses (workflow #7 – #9)>Use of inhibitors in automated hematological studies

21 See Methods>Preparation of RBCs>Processing of RBCs for assays 1–4

22 See Methods>Statistical analysis>one-way analysis of variance (ANOVA)

23 See Methods>Preparation of RBCs

24 See (a) Paper #1>Results>Effect of pCMBS or DIDS on O_2_ offloading from RBCs … &… (b) ..>Effect of genetic deletions on O_2_ offloading from RBCs … & … (c) ..>Hematological and related parameters>Automated hematology

25 See Paper #3>Results

26 Performed by board-certified hematologist and co-author HJM

27 Paper #1>Methods> Stopped-flow absorbance spectroscopy (workflow #1)

28 See Methods>Still microphotography and microvideography of living RBCs

29 See Methods>Flow cytometry (workflow #6 & #6′, #15 & #15′)>Sample preparation

30 We define DRAQ-5^dim^ staining intensity as increased when compared to the negative control but <10^−4^

31 See Methods>Flow cytometry (workflow #6 & #6′, #15 & #15′)>Imaging flow cytometry (IFC)

32 See Methods>Flow cytometry>Light-scattering flow cytometry

33 The ratio of the minor axis to the major axis of an object, calculated based on the intensity of the pixels within the object in the brightfield image (Channel 1).

34 The ratio of the minor axis to the major axis of an object, calculated based on the fluorescence intensity of the pixels within the object in channel 11.

35 See Methods>Statistical analysis

36 See Results>Morphometry>Imaging of living, tumbling RBCs

37 See Results>Proteomics>Effects of knockouts on PMA proteins of greatest inferred abundance

38 To account for 100% of each class of protein, consult Supporting file 1, Tab 7, titled “Rank ALL proteins each strain”, cell coordinates “I1111” (WT).

39 See Paper #1>table 4

40 From preliminary work on human RBCs by Fraser J Moss, Pan Zhao, Richard Moon, Walter F Boron.

41 See Paper #1>figure 3*f* & figure 4*f*

42 See Paper #3>Results>Accommodation for non-BCDs

43 See Paper #3>Methods>Mathematical modeling and simulations>Calculation of RBC thickness based on the geometry of a torus

44 Kui Xu, Pan Zhao, Rossana Occhipinti, Fraser J. Moss, and Walter F. Boron (unpublished)

45 See Paper #1>Results>Accommodation for Hemolysis>Accommodation of raw-*k*_HbO_2__ for hemolysis

46 See Paper #3>Results> Predicted dependence of O_2_ offloading (*k*_HbO_2__) on the O_2_ permeability of RBC membrane (*P*_M,O_2__)

47 See Paper #1>Results>Effect of genetic deletions on O_2_ offloading from RBC>Combined effects of dKO and pCMBS.

48 Pan Zhao and Walter Boron.

49 See Paper #1>Results>Effect of genetic deletions on O2 offloading from RBC>Effects of single and double knockouts.

50 See Paper #1>Results>Effect of genetic deletions on O_2_ offloading from RBC>Combined effects of dKO and pCMBS

52 See Methods>Flow cytometry>Light-scattering flow cytometry

51 See Methods>Flow cytometry (workflow #6, 6′ & #15, 15′)>Imaging flow cytometry (IFC)

53 See Methods>Flow cytometry (workflow #6 & #6′, #15 & #15′)>Sample preparation

54 See Methods>Preparation of RBCs>Collection of blood & ..>Processing of RBCs for assays 1–4

55 See Methods>Still microphotography and microvideography of living RBCs

56 See Methods>Preparation of RBCs>Collection of blood & ..>Processing of RBCs for assays 1–4

57 See Methods>Still microphotography and microvideography of living RBCs

58 See Methods>Preparation of RBCs>Collection of blood & ..>Processing of RBCs for assays 1–4

59 See Methods>Still microphotography and microvideography of living RBCs

60 See Methods>Preparation of RBCs>Collection of blood & ..>Processing of RBCs for assays 1–4

61 See Methods>Still microphotography and microvideography of living RBCs

